# Sphingolipid-Driven Lipotoxicity Promotes Calcific Aortic Valve Disease Through ER-Stress–NF-κB–EndMT Signaling and Is Therapeutically Targetable Through Serine Palmitoyltransferase Inhibition

**DOI:** 10.64898/2026.10.07.757226

**Authors:** E. Rawish, T. Rusack, Peter Letsch, J. Inderhees, K. Händler, C. Gragoll, J.C. Voran, M. Schneider, K. Kurz, M.R. Hosen, M. Behrensen, S. Denker, A. Künstner, T. Sauer, J. Duque-Escobar, T. Schmidt, D. Frank, T. Stiermaier, W. Raasch, O.J. Müller, T. Zeller, H. Busch, T. Gemoll, C. Knosalla, M. Spielmann, Z. Aherrharou, S.A. Summers, I. Eitel, C. Frerker

## Abstract

**Background:** Calcific aortic valve disease (CAVD) lacks disease-modifying therapy. LDL-cholesterol lowering does not attenuate valvular calcification, suggesting cholesterol-independent drivers. Thus, we tested whether ceramide accumulation drives CAVD and whether inhibiting sphingolipid de novo synthesis is therapeutic.

**Methods:** We integrated lipidomics and proteomics of calcified and non-calcified human aortic valves with matched plasma. Mechanistic effects of ceramides were tested in primary human valvular interstitial cells (VICs). Therapeutic efficacy of serine palmitoyltransferase (SPT) inhibition with Myriocin was evaluated in ApoE^−/−^ mice with PCSK9^DY^-AAV–driven hyperlipidemia on Western diet using preventive treatment and therapeutic initiation after echocardiographically verified gradient elevation. Myriocin was additionally tested in a normolipidemic wild-type aortic valve wire-injury model. Hemodynamics, valve structure, lipid composition, and cell-state transitions were assessed by echocardiography, histology, lipidomics, and single-cell RNA sequencing.

**Results\:** Calcified human valves showed enrichment of long-chain ceramides linked to NF-κB, endothelial-to-mesenchymal transition (EndMT), and ER-stress/unfolded protein response (UPR) programs, whereas matched plasma sphingolipids were not elevated. In hVICs, long-chain C18 ceramide induced GRP78/BiP, CHOP, and ATF4, activated NF-κB, and promoted dose-dependent calcification. *In vivo*, SPT inhibition with Myriocin lowered valvular ceramides, reduced leaflet inflammation and mineralization, and improved cusp separation and transvalvular hemodynamics without lowering plasma LDL-cholesterol. In a therapeutic protocol initiated after gradient elevation, Myriocin halted further hemodynamic progression, providing the first evidence that de novo sphingolipid synthesis can be targeted after established CAVD onset *in vivo*. Myriocin also attenuated injury-induced stenosis in normolipidemic mice. Single-cell RNA sequencing showed reduced NF-κB/EndMT and ER-stress signatures and shifted VIC states away from osteogenic phenotypes.

**Conclusions:** Human valve lipidomics and human VIC experiments identify ceramide-driven lipotoxicity as a driver of an ER-stress–NF-κB–EndMT axis in CAVD. *In vivo*, de novo sphingolipid synthesis is an actionable, LDL-independent target capable of modifying established hemodynamic disease progression, supporting SPT inhibition as a tractable strategy for medical CAVD therapy.

**What is new?:**

- Calcified human aortic valves show local long-chain ceramide enrichment linked to NF-κB, EndMT, and ER-stress/UPR programs, whereas matched plasma lacks a corresponding sphingolipid increase.
- C18 ceramide directly promotes ER-stress– and NF-κB-dependent calcification in primary human valvular interstitial cells.
- To our knowledge, this is the first demonstration that pharmacologic inhibition of de novo sphingolipid synthesis, initiated after echocardiographically verified gradient elevation, attenuates further hemodynamic progression of CAVD *in vivo*.

What are the clinical implications?

- CAVD progression may involve valve-local ceramide lipotoxicity rather than LDL cholesterol alone.
- SPT inhibition provides preclinical proof-of-concept for LDL-independent disease modification, including established hemodynamic disease.
- Clinically translatable SPT inhibitors or targeted delivery strategies warrant evaluation as medical approaches to slow calcific aortic stenosis progression.

## Introduction

Calcific aortic valve disease (CAVD) is the most common valvular heart disease requiring intervention in high-income countries [1]. Once clinically significant, untreated aortic stenosis carries substantial mortality, emphasizing the need for disease-modifying therapies beyond valve replacement [2]. Despite this burden, no medical therapy slows CAVD progression: low-density lipoprotein (LDL-)cholesterol lowering, effective in atherosclerosis, was ineffective in randomized controlled trials [3, 4]. Sphingolipids have been implicated in cardiometabolic disease through lipotoxic mechanisms [5]. When neutral lipid storage is exceeded, bioactive ceramides accumulate and engage inflammatory programs [5–7]. In human atherosclerosis, ceramide burden correlates with plaque vulnerability [8], and pharmacological inhibition of serine palmitoyltransferase (SPT), the rate-limiting enzyme of de novo sphingolipid biosynthesis, reduces atherosclerosis in ApoE-deficient mice [9–11]. By contrast, while statins mitigate atherosclerosis via LDL lowering in rodents [12], murine CAVD models show no meaningful reduction of valvular calcification or stenosis with statins [13, 14]. Although CAVD and vascular atherosclerosis share chronic inflammation [15], CAVD features fibro-calcific remodeling in largely avascular leaflets, indicating distinct tissue biology [16].

Taken together, LDL-dependent anti-atherosclerotic effects do not translate to CAVD and motivate evaluation of cholesterol-independent lipotoxic pathways – particularly sphingolipid metabolism – in degenerative aortic valve calcification. Human valve lipidomic and biochemical studies support sphingolipid dysregulation: ceramides concentrate in heavily calcified regions and extracellular particles from stenotic valves exhibit an increased sphingomyelin-to-phosphatidylcholine ratio [17, 18]. However, the causal relevance of sphingolipids to degenerative calcification in native valves, the mechanistic links to inflammatory and osteogenic programs, and the therapeutic tractability via upstream SPT inhibition remain undefined. Establishing such links would constitute a conceptual shift beyond lipoprotein-centric models with potential therapeutic implications in valvular heart disease.

Here, we show that ceramide-driven lipotoxicity engages ER stress and NF-κB signaling and promotes endothelial-to-mesenchymal transition (EndMT) to drive calcific remodeling in native degenerative aortic valves. Integrating multi-omics analyses of human valve tissue with mechanistic experiments in human valvular interstitial cells (hVICs) and complementary *in vivo* models, we delineate a cholesterol-independent ceramide–ER stress–NF-κB–EndMT axis. Pharmacologic inhibition of serine palmitoyltransferase (SPT) with Myriocin lowers valvular ceramide accumulation and attenuates calcification in a metabolically driven ApoE^−/−^+PCSK9^DY^-AAV model without lowering plasma LDL-cholesterol, prevents calcifying valve stenosis in a normolipidemic wild-type wire-injury model, and halts progression when initiated after established stenosis. Across species, a long chain ceramide signature tracks calcification severity, underscoring translational relevance.

Together, our findings provide proof-of-concept that targeting ceramide accumulation, rather than cholesterol, can modify key fibro-calcific and inflammatory programs in CAVD. They position SPT inhibition as an actionable upstream strategy to disrupt ceramide-driven valvular lipotoxicity and support ceramides as pharmacologically targetable mediators of calcific remodeling beyond LDL-centric pathways.

## Methods

Expanded Methods are provided in the Supplemental Material.

### Animal Studies

All animal care and procedures were conducted in accordance with NIH guidelines and approved by the local regulatory authority (Reference Number 49-9/23, Ministerium für Landwirtschaft, ländliche Räume, Europa und Verbraucherschutz des Landes Schleswig-Holstein) and reported according to ARRIVE. For a metabolically driven CAVD model, male and female ApoE^−/−^ mice (B6.129P2-Apoetm1Unc/J) were injected at 7 weeks with AAV-PCSK9^DY^ (2×10^11^ viral particles/mouse) [19] and fed a cholesterol-enriched Western diet. Mice received vehicle, Myriocin (0.3 mg/kg/day) or GW4869 (30 mg/kg/day) [20] in drinking water with concentrations adjusted to water consumption; AAV transduction was confirmed by measuring plasma PCSK9. For preventive experiments, treatment was started with diet/AAV and continued for 21 weeks. For therapeutic experiments, mice were maintained on Western diet for 17 weeks, aortic valve stenosis was verified by echocardiography, and Myriocin or vehicle was initiated thereafter and continued until week 24. In a non-hyperlipidemic model, C57BL/6J mice on chow diet underwent aortic valve wire injury [21] and were treated with Myriocin or vehicle until endpoint. Terminal blood and tissues were collected under anesthesia.

### Mouse Echocardiography

Transthoracic echocardiography was performed under ketamine-based sedation at prespecified timepoints (week 21 in preventive experiments; weeks 17 and 24 in therapeutic experiments, and at endpoint in the wire-injury model). Doppler-derived peak transvalvular velocity (Vmax) and mean pressure gradient (dpmean) were quantified alongside leaflet morphology (thickness/cusp separation) and left ventricular systolic/diastolic function (LVEF, E/E′, IVRT).

### Single-cell RNA Sequencing

Aortic valves were enzymatically dissociated (n=4/group), processed with 10x Genomics single-cell 3′ chemistry, sequenced, and analyzed using Cell Ranger and Seurat with Harmony integration. Pathway-level differences were assessed by GSEA and ssGSEA (details in the Supplemental Material).

### Histology

Aortic valves were fixed, embedded, and sectioned (5 µm). Calcification, fibrosis, ceramide deposition, and immune cell infiltration were assessed by histology/immunostaining (eg, calcein, Masson’s trichrome, anti-ceramide staining, MOMA-2) (details in the Supplemental Material).

### Human Aortic Valves and Plasma Samples

Human aortic valves and plasma were obtained from patients with CAVD and non-calcified controls during surgery (ethics approval EA4/028/12, Charité, Berlin). A validation plasma cohort from TAVI patients was collected at the Christian-Albrechts-Universität Kiel (D 506/22; AZ F 01/24); fasting plasma was obtained and CT-based calcification scores were quantified.

### Lipidomics and Proteomics

Lipids were extracted from plasma and valve tissue and quantified by LC-MS/MS. Valve tissue proteomes were analyzed by LC-MS/MS with data-independent acquisition; downstream analyses included differential abundance testing, WGCNA, and integrative correlation analyses (details in the Supplemental Material).

### Culture of Primary Human Valvular Interstitial Cells

Primary hVICs (passages 2–4) were cultured in valvular interstitial cell medium and subjected to calcification medium for up to 21 days. Cells were stimulated with C2 ceramide and long-chain C18 ceramide, as indicated. Calcification was assessed by Alizarin Red staining and calcium quantification; inflammatory signaling was assessed by IL-6 ELISA and an NF-κB pathway array.

### Immunoblotting and RT-qPCR

ER stress/UPR activation in hVICs after ceramide stimulation was assessed by immunoblotting (eg, GRP78/BiP) and RT-qPCR for UPR genes (including DDIT3/CHOP, HSPA5/GRP78, ATF4 and related markers).

### Statistical Analysis

Animal sample sizes were determined by power analysis based on the transvalvular mean gradient. Data were analyzed using GraphPad Prism 10 or SPSS 25. Parametric or non-parametric tests were applied depending on normality and variance homogeneity. Omics data were analyzed using DESeq2, edgeR, and limma-trend with multiple testing correction. A p-value < 0.05 was considered statistically significant.

## Results

### Lipidome Characterization of Human Aortic Valve Tissue Reveals Sphingolipid Accumulation in CAVD

We profiled sphingolipids by liquid chromatography–mass spectrometry (LC-MS) in aortic valve tissue and plasma from 29 patients with or without valve calcification to distinguish local from systemic changes (for clinical characteristics see (Table 1). Analysis of 748 lipid species in aortic valves revealed a significantly altered lipidome, as 3D sPLS-DA (sparse Partial Least Squares Discriminant Analysis) showed clear group discrimination between patients with or without CAVD (Fig. 1A) (Supplementary Tables 1-2). Several sphingolipid species—including ceramides (Cer), hexosylceramides (HexCer), and sphingomyelins (SM)—as well as ether-linked lysophosphatidylcholines (LPC O) were significantly increased in calcified valve tissues, while triglycerides (TG) were decreased (Fig. 1A). Similarly, fatty acids (FA) and acylcarnitines (CAR) were increased in aortic valve tissue of CAVD patients (Fig. 1A), consistent with dysfunctional mitochondrial fatty acid oxidation (FAO), as CAR constitute a transport form of FA utilized for energy production in mitochondria [22].

**Fig. 1.**
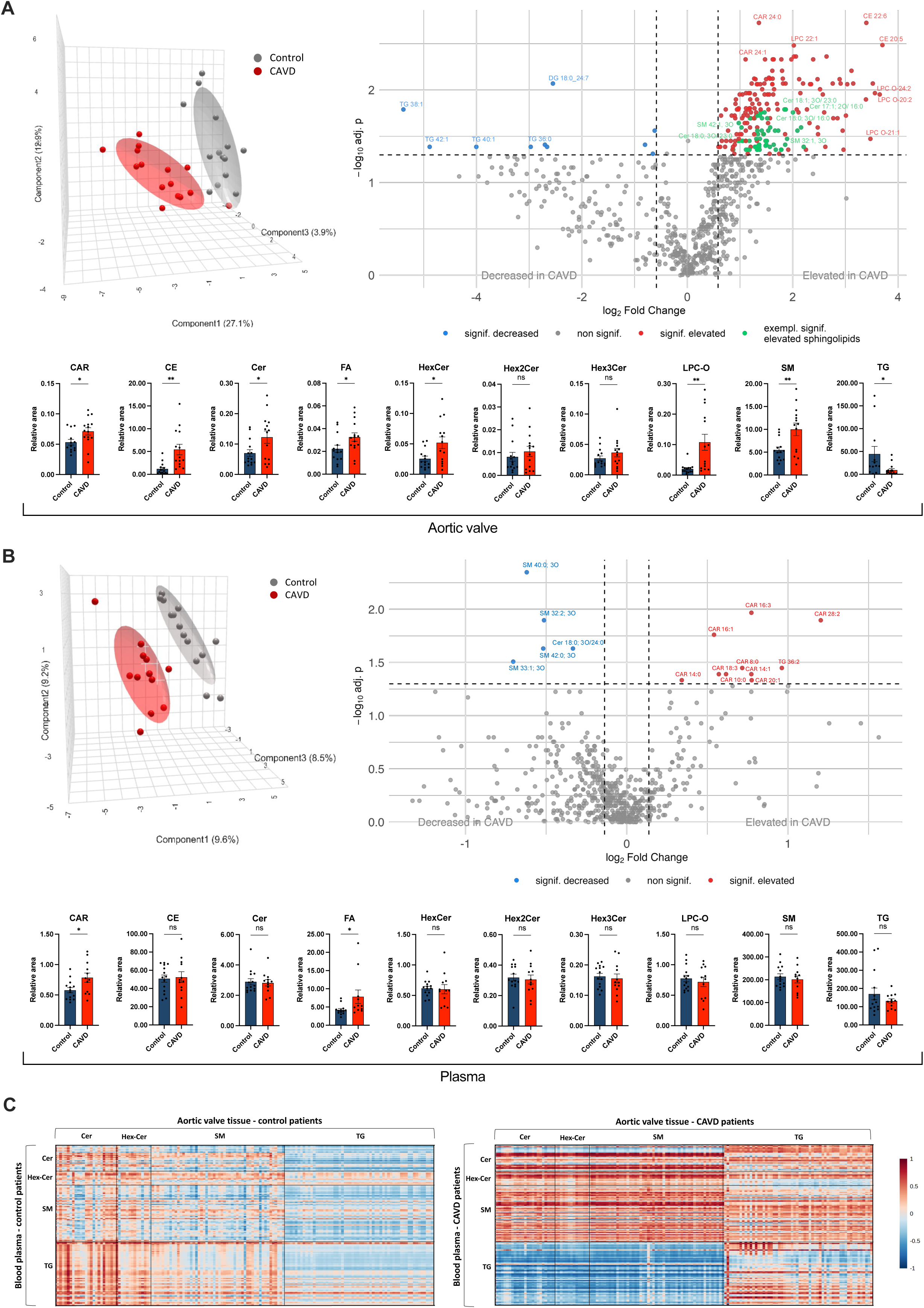
Human aortic valve and plasma lipidomics reveal local sphingolipid enrichment in CAVD and a divergent circulating lipid signature. A, Top left: 3D sparse partial least squares discriminant analysis (sPLS-DA) of 748 detected lipid species in aortic valve tissue from control patients (n=14) and patients with CAVD (n=15). Top right: volcano plot of tissue lipids (CAVD versus control). Bottom: lipid classes in valve tissue. Cer, HexCer, SM, ether-linked lysophosphatidylcholines (LPC-O), fatty acids, and acylcarnitines were increased, whereas triglycerides were decreased. B, Top left: 3D sPLS-DA of 609 plasma lipid species from control patients (n=14) and patients with CAVD (n=12). Top right: volcano plot of plasma lipids. Bottom: plasma lipid classes. Circulating acylcarnitines and fatty acids were increased, whereas total plasma sphingolipid classes were not increased. C, Correlation matrices of lipid classes in aortic valve tissue versus matched plasma shown separately for controls and CAVD. Bar charts show mean ± SEM. Tissue comparisons in A were analyzed by Wilcoxon rank-sum test for volcano plots (fold-change threshold 1.5; FDR-adjusted P<0.05) and Student’s t test for lipid-class summaries; plasma comparisons in B used Wilcoxon rank-sum test for volcano plots (fold-change threshold 1.1; P<0.05) and Student’s t test for lipid-class summaries. Pearson correlation coefficients are shown in C. *P<0.05, **P<0.005. Abbreviations: CAR, acylcarnitine; Cer, ceramide; CE, cholesteryl ester; DG, diacylglycerol; FA, fatty acid; HexCer, hexosylceramide; LPC-O, ether-linked lysophosphatidylcholine; SM, sphingomyelin; TG, triglyceride.

**Table 1:** Characteristics of CAVD patients and Controls for aortic valve Omics analysis.

| Parameter (mean ± SD or %) | Control (n=14) | CAVD (n=15) | P-Value |
| --- | --- | --- | --- |
| Age | 66.0 ± 10.7 | 69.4 ± 7.5 | 0.35 |
| Female | 33.3% | 35.3% | 1 |
| BMI | 26.1 ± 3.0 | 27.8 ± 4.3 | 0.22 |
| Dpmean aortic valve [mmHg] | 9.1 ± 4.5 | 49.6 ± 18.8 | <0.001 |
| Severe aortic valve stenosis | 0.0% | 100.0% | <0.001 |
| Severe aortic valve insufficiency | 78.6% | 20.0% | 0.003 |
| Aortic Root Dilation | 64.3% | 0.0% | <0.001 |
| LV-EF [%] | 55.9 ± 9.9 | 58.5 ± 6.0 | 0.44 |
| eGRF [ml/min/1,72m<sup>2</sup>] | 73.0 ± 16.8 | 71.4 ± 22.7 | 0.84 |
| LDL Cholesterol [mg/dL] | 136.0 ± 24.4 | 114.8 ± 37.3 | 0.38 |
| Triglycerides [mg/dL] | 141.0 ± 56.2 | 128.3 ± 12.55 | 0.67 |
| Hypertension | 50.0% | 60.0% | 0.72 |
| Type 2 Diabetes | 21.4% | 26.7% | 1 |
| Coronary artery disease | 35.7% | 46.7% | 0.71 |
| Medication |  |  |  |
| Beta Blockers | 57.1% | 66.7% | 0.71 |
| ACE Inhibitors/AT1 Blockers | 50.0% | 53.3% | 1 |
| Diuretics | 50.0% | 33.3% | 0.46 |
| Statins | 35.7% | 46.7% | 0.71 |
| Calcium Channel Blockers | 35.7% | 26.7% | 0.7 |

Plasma analysis also rendered good group distinction and revealed elevated levels of medium-and long-chain CAR (Fig. 1B). In contrast to aortic valve tissue, levels of several sphingomyelin species and Cer 18:0;O3/24:0 were reduced in plasma of CAVD patients, whereas total plasma sphingolipids showed no significant increases compared with controls (Fig. 1B). Nevertheless, a correlation analysis of lipid species in aortic valve tissue with plasma lipids in CAVD patients and controls revealed distinct patterns: CAVD patients showed a stronger positive correlation between ceramides, hexosylceramides, and sphingomyelins in aortic valve tissue and plasma compared to controls (Fig. 1C). Additionally, CAVD patients exhibited a negative correlation between triglycerides in plasma and sphingolipids in valve tissue. Hence, our lipidomic analysis identifies an accumulation of sphingolipids in calcified aortic valves together with a systemic plasma signature consistent with FAO stress rather than generalized plasma sphingolipid elevation.

### Bioinformatic Integration of Lipidomics and Proteomics Data Connects Local Ceramide Accumulation to Activation of Pro-Inflammatory and Osteogenic Pathways In Aortic Valve Tissue

To substantiate the lipidomics findings and elucidate their pathophysiological relevance, we performed proteomic analyses on aortic valve specimens. In CAVD patients, protein abundance of 94 genes was upregulated and 125 were downregulated (Fig. 2A). Unsupervised hierarchical clustering highlighted an upregulation of pro-inflammatory proteins, particularly those involved in NFκB signaling (Fig. 2B), alongside a suppression of HSPA2, a cytoprotective chaperone, indicating endoplasmic reticulum (ER) stress (Fig. 2A) [23]. Weighted gene co-expression network analysis (WGCNA) was used to develop gene correlation networks, allowing module clustering (Supplementary Fig. 1A-I), which identified ME1, ME3, and ME4 as leading modules associated with CAVD traits. Gene set enrichment analysis (GSEA) revealed enrichment of NFκB signaling in ME1 and Endothelial-to-Mesenchymal Transition (EndMT)-associated pathways in ME3, reinforcing recent single-cell transcriptomics findings linking EndMT to aortic valve thickening [24]. Regarding intersections with lipid metabolism, impaired FAO – which was indicated by our lipidomics analysis – has been shown to activate EndMT [25], and ME4 exhibited depletion of apical junction components, potentially altering endothelial lipid permeability. Key FA and TG metabolism proteins were altered in calcified valve tissue, notably a reduction in Perilipin-1 (PLIN1), which protects intracellular LDs from lipases [26]. Its downregulation may contribute to TG depletion (Fig. 1A), intracellular FFA excess, and subsequent lipotoxicity and inflammation. Additionally, Toll-like receptor 2 (TLR2) was significantly upregulated (Fig. 2A), a regulator of ceramide de novo synthesis in response to excess saturated FFAs [27–29].

**Fig. 2.**
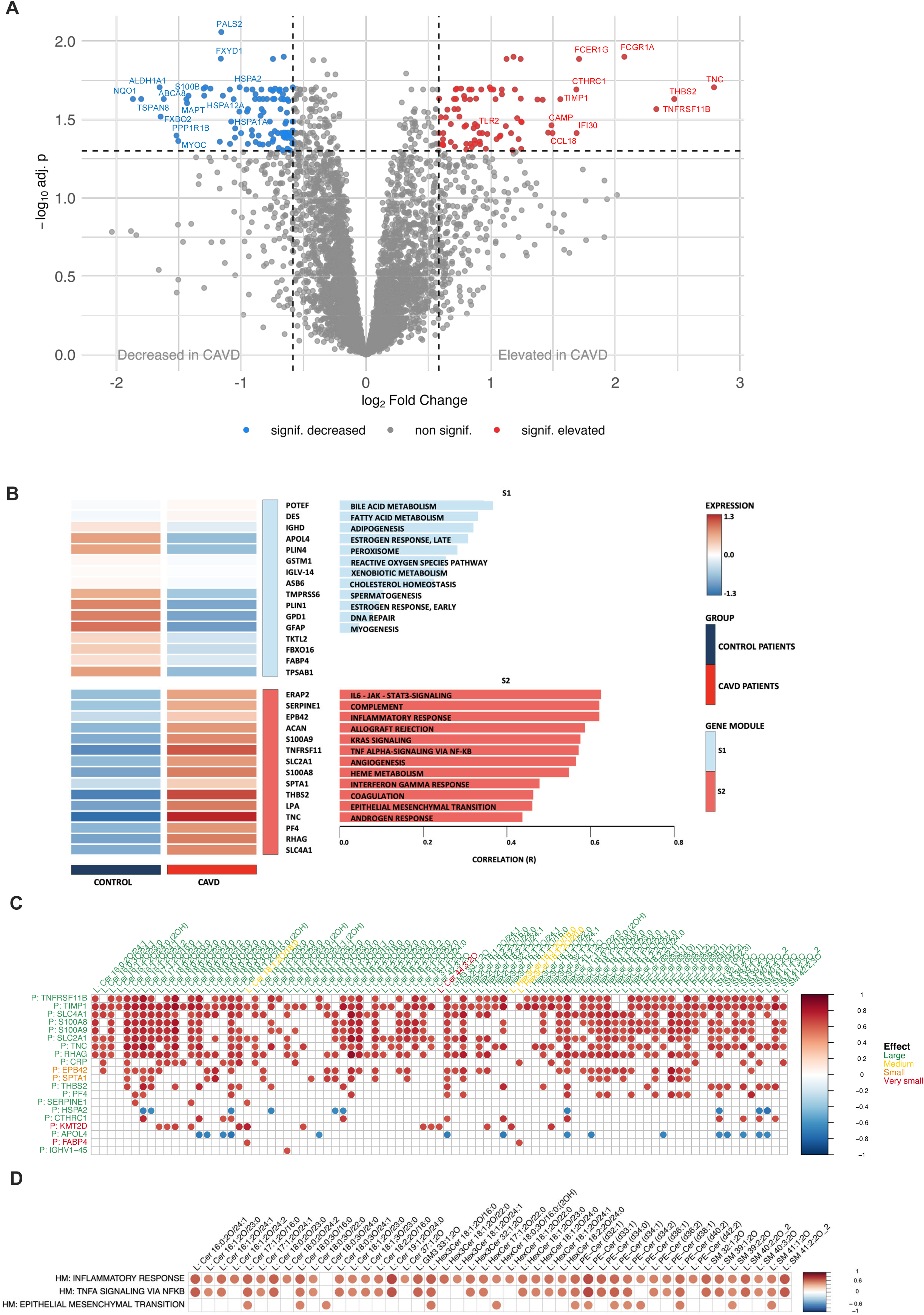
Human valve proteomics links ceramide enrichment to inflammatory, stress, and pro-calcific programs in CAVD. A, Volcano plot of differentially abundant proteins in CAVD versus control aortic valves. Proteins involved in inflammation (eg, FCGR1A, IFI30, CCL18, TLR2), ER/oxidative stress (eg, HSPA2, NQO1), and osteogenic remodeling (eg, TNFRSF11B, TNC, THBS2, TIMP1, CTHRC1) are highlighted. B, Unsupervised hierarchical clustering analysis and functional annotation of protein modules show upregulation of pro-inflammatory proteins, while fatty acid metabolism proteins are downregulated in CAVD. C, Partial correlation analysis between selected lipids and proteins demonstrates positive associations of long-chain ceramides and related sphingolipids with pro-inflammatory and pro-calcific proteins and inverse associations with HSPA2. Label color indicates Cohen’s d effect size. D, Partial correlation analysis between selected lipids and hallmark gene-set scores derived from the proteomic dataset reveals positive associations of long-chain ceramides with inflammatory response, TNFα signaling via NF-κB, and epithelial/EndMT-related programs. Differential protein abundance in A was assessed using DESeq2 (Wald test), edgeR (QLF test), and limma-trend, with the maximum q-value of the 3 methods used as aggregate q-value (fold-change threshold 0.5; FDR-adjusted P<0.05). Correlations in C and D were calculated by Spearman partial correlation adjusting for age and sex; only associations with P<0.05 are shown.

To explore the relationship between proteome changes and sphingolipidome alteration, we performed partial correlation analysis, revealing strong correlations between ceramide species and pro-calcific proteins, including Osteoprotegerin (TNFRSF11B), Tenascin C (TNC), Tissue Inhibitor of Metalloproteinases 1 (TIMP1), Thrombospondin 2 (THBS2), and Collagen Triple Helix Repeat Containing 1 (CTHRC1). Furthermore, several canonical ceramides (Cer 18:1;O2/X) and their precursor dihydroceramides (Cer 18:0;O2/X)), as well as less abundant ceramides comprising a 16:1 sphingoid base (Cer 16:1;O2/X), correlated positively with the pro-inflammatory S100A8/A9 complex but negatively with HSPA2 (Fig. 2C). These findings underscore the interplay between inflammation, sphingolipid dysregulation, and CAVD pathogenesis, further supported by partial correlation with hallmark inflammatory gene sets (Fig. 2D).

### Inhibition of Ceramide Synthesis Restores Aortic Valve Sphingolipidome and Leaflet Morphology in a Metabolic Mouse Model of CAVD Independent of Plasma Cholesterol Levels

Building upon our human multi-omics findings, we next investigated the therapeutic potential of modulating ceramide accumulation *in vivo*. To address both metabolic and non-metabolic drivers of CAVD and to disentangle systemic lipid confounding from valve-intrinsic mechanisms, we employed complementary mouse models and preventive as well as therapeutic treatment paradigms.

First we employed a metabolic CAVD model (ApoE^−/−^+ PCSK9^DY^-AAV) that induces chronic hyperlipidemia and recapitulates macrophage infiltration and leaflet sclerosis [30] (Fig. 3A). Extending prior work that lacked hemodynamic assessment [30], we demonstrate hemodynamically significant aortic stenosis at 21 weeks in ApoE^−/−^+PCSK9^DY^-AAV mice compared with wild-type controls (Fig. 5B-C).

**Fig. 3.**
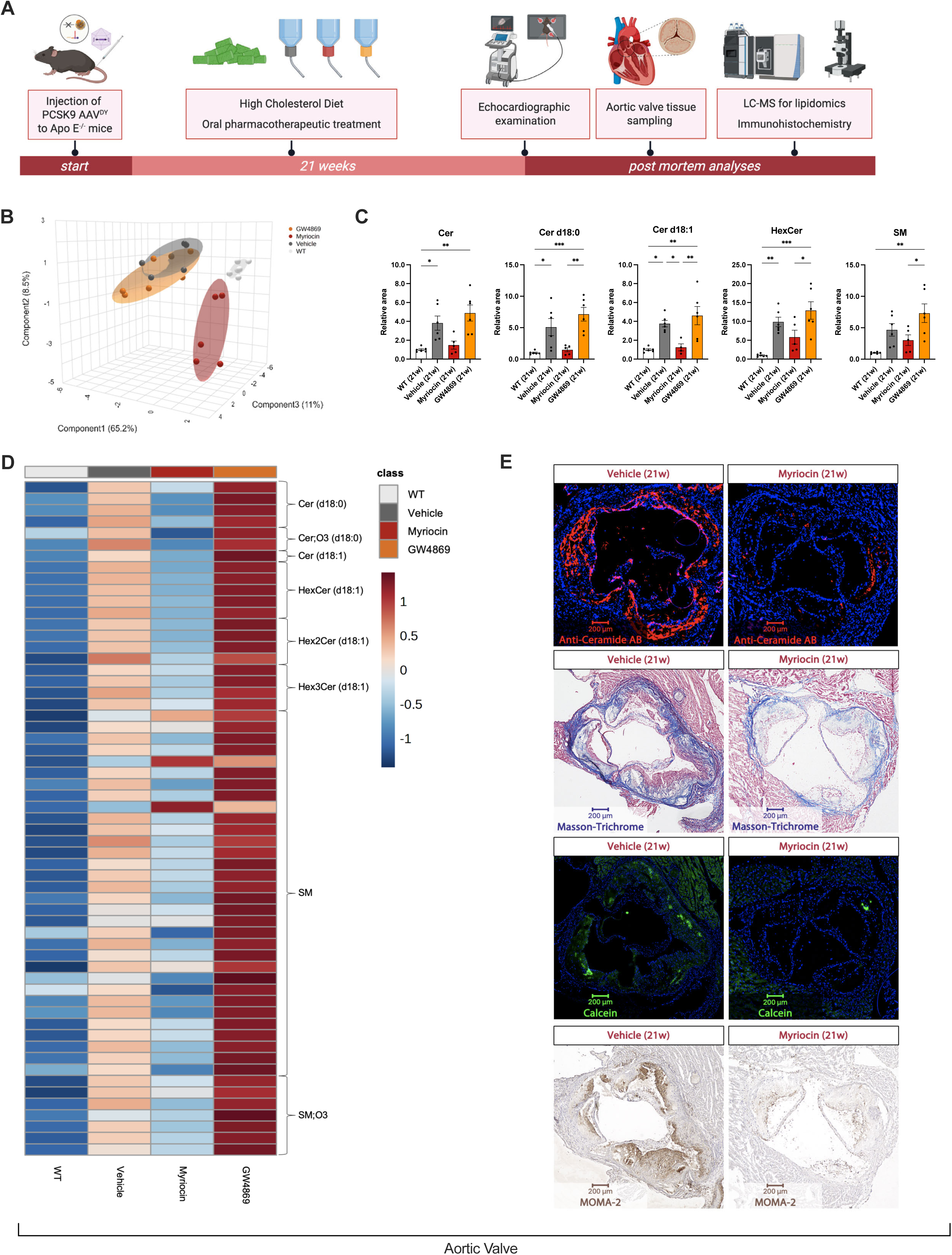
SPT inhibition restores the valvular sphingolipidome and attenuates leaflet pathology in a metabolic mouse model of CAVD. A, Preventive study design in ApoE^−/−^ mice injected with PCSK9^DY^-AAV and fed Western diet for 21 weeks with oral vehicle, Myriocin, or GW4869. Echocardiography was performed before terminal valve harvesting for lipidomics and histology. B, 3D sPLS-DA of valve-tissue sphingolipid profiles across wild-type (WT), vehicle-, Myriocin-, and GW4869-treated groups. C, Valvular sphingolipid classes/species showing reduction of total ceramides, Cer d18:0, Cer d18:1, HexCer, and SM with Myriocin. D, Heat map of significantly altered sphingolipid species in aortic valve tissue; full annotation is provided in Supplementary Table 3. E, Representative anti-ceramide immunostaining, Masson’s trichrome, Calcein, and MOMA-2 staining of aortic valves from vehicle– and Myriocin-treated mice at week 21. Bar charts show mean ± SEM. Statistical analyses were performed by 1-way ANOVA followed by Bonferroni post hoc correction (C). *P<0.05, **P<0.01, ***P<0.001.

**Fig. 4.**
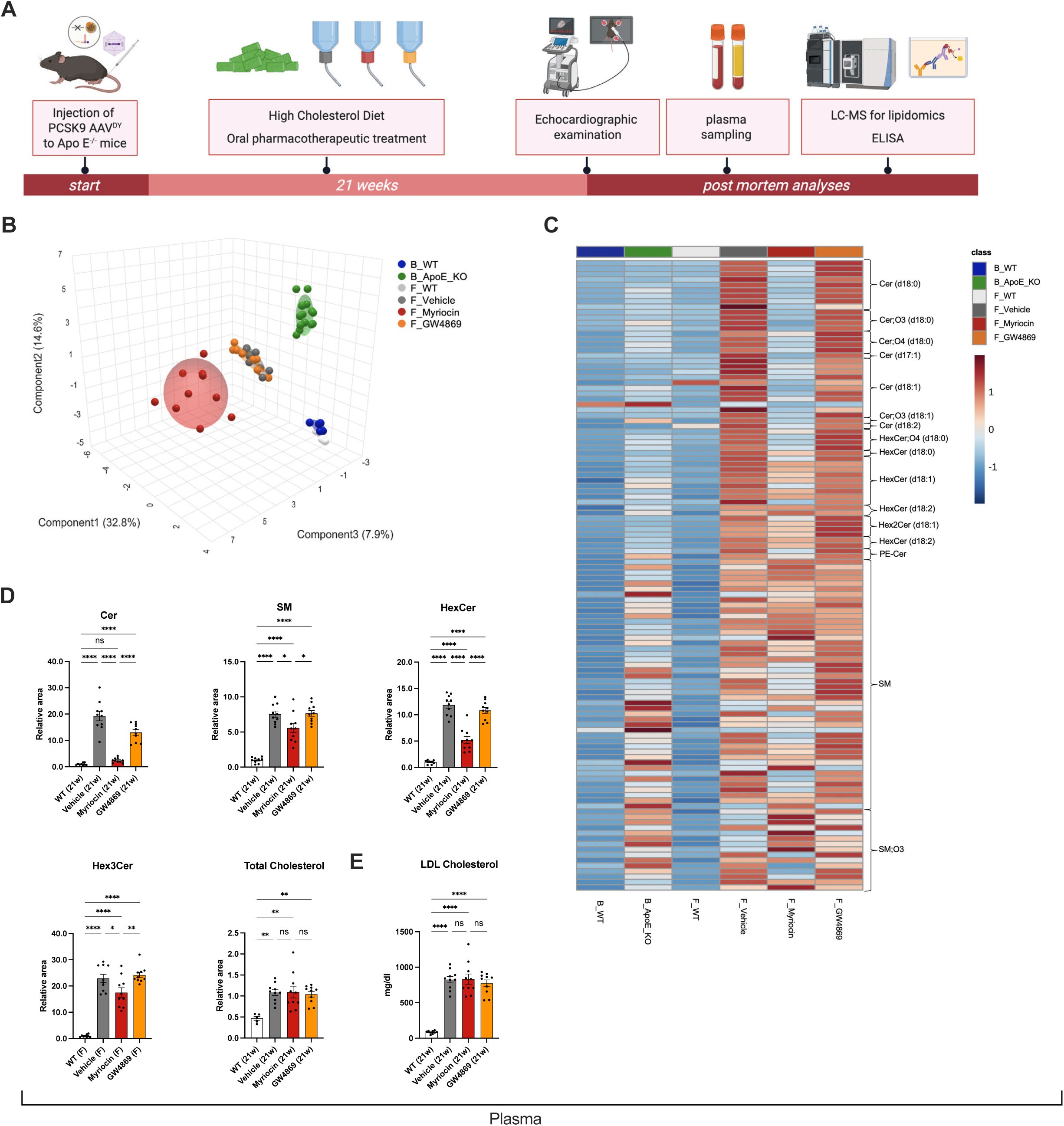
Myriocin lowers circulating sphingolipids without reducing plasma cholesterol in the metabolic CAVD model. A, Study workflow for plasma sampling, lipidomics, and LDL-cholesterol ELISA in ApoE^−/−^+PCSK9^DY^-AAV mice after 21 weeks of Western diet and treatment. B, 3D sPLS-DA of plasma lipidome profiles, including baseline (B_) and final (F_) samples from WT and ApoE KO groups and final samples from vehicle-, Myriocin-, and GW4869-treated ApoE KO+PCSK9^DY^-AAV mice. C, Heat map of significantly altered plasma sphingolipid species; full annotation is provided in Supplementary Table 4. D, Circulating sphingolipid classes showing reduced Cer, SM, HexCer, and Hex3Cer with Myriocin, whereas total cholesterol remained elevated across all ApoE KO+PCSK9^DY^-AAV treatment groups. E, ELISA-based LDL-cholesterol measurements confirm persistently elevated LDL-cholesterol despite Myriocin treatment. Bar charts show mean ± SEM. Statistical analyses were performed by 1-way ANOVA followed by Bonferroni post hoc correction (D-E). *P<0.05, **P<0.01, ***P<0.001, ****P<0.0001.

**Fig. 5.**
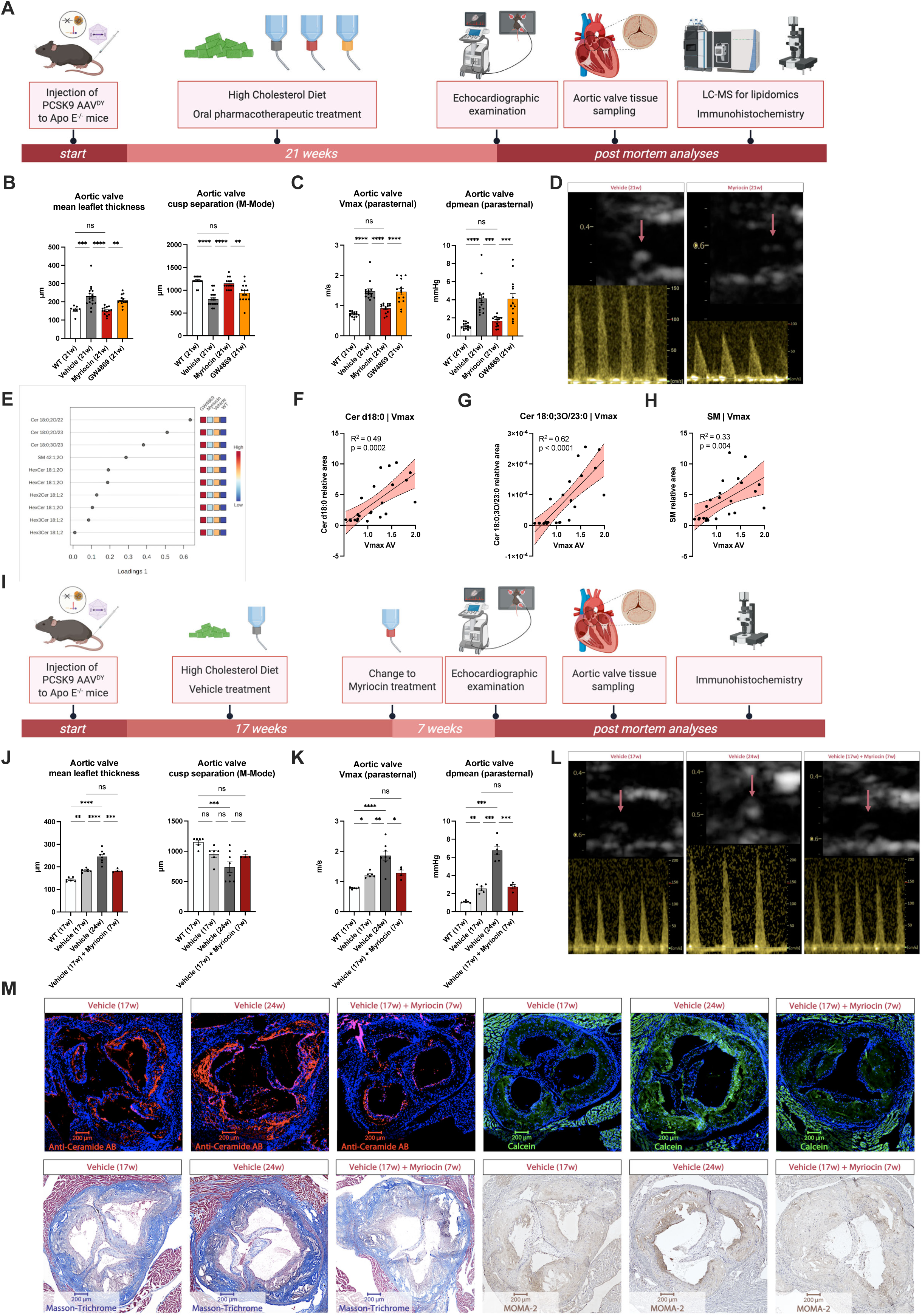
SPT inhibition preserves valve hemodynamics, identifies Cer 18:0;O3/23:0 as a cross-species correlate, and attenuates progression of established aortic stenosis. A, Preventive study design. B, Mean leaflet thickness and cusp separation measured by echocardiography after 21 weeks. C, Peak transvalvular velocity (Vmax) and mean pressure gradient (dpmean) after 21 weeks. D, Representative echocardiographic images and Doppler traces from vehicle– and Myriocin-treated mice. E, Loadings plot identifying Cer 18:0;O3/23:0 and related sphingolipids as principal drivers of group separation. F-H, Correlation of Vmax with total Cer d18:0, Cer 18:0;O3/23:0, and SM relative area in aortic valve tissue. I, Therapeutic study design in ApoE^−/−^+PCSK9^DY^-AAV mice maintained on Western diet for 17 weeks to establish stenosis, then randomized to Myriocin or vehicle for 7 additional weeks. J, Mean leaflet thickness and cusp separation across WT at week 17, vehicle at week 17, vehicle at week 24, and Myriocin from week 17 to week 24. K, Vmax and dpmean across the same groups. L, Representative echocardiographic images and Doppler traces for the therapeutic study before and after treatment with Myriocin or vehicle. M, Anti-ceramide immunostaining, Calcein, Masson’s trichrome, and MOMA-2 staining in the therapeutic study before treatment initiation at week 17 and after Myriocin treatment or vehicle at week 24. Bar charts show mean ± SEM. Statistical analyses were performed by 1-way ANOVA followed by Bonferroni post hoc correction (B-C, J-K). Correlation panels (F-H) show linear regression with 95% confidence bands. *P<0.05, **P<0.01, ***P<0.001, ****P<0.0001.

Importantly, lipidomic profiling of valve tissue confirmed substantial sphingolipid accumulation, including pro-inflammatory long-chain Cer 18 species (Cer 18:0 and Cer 18:1) [31], in vehicle-treated *ApoE KO + PCSK9^DY^ AAV* animals. This valvular ceramide burden was markedly reduced towards wild-type levels upon pharmacological inhibition of SPT with Myriocin, whereas GW4869 failed to normalize the sphingolipidome (Fig. 3B-D, full annotation in Supplementary Table 3). These findings were corroborated by histology, showing reduced ceramide deposition in the aortic valve under Myriocin, accompanied by reduced calcification (Calcein), less fibrosis (Trichrome), and decreased macrophage infiltration (Fig. 3E). Accordingly, SPT inhibition reduced leaflet thickness and improved cusp separation compared with vehicle, restoring structural indices towards wild-type levels (Fig. 5B). In contrast, inhibition of neutral sphingomyelinase with GW4869 failed to improve sphingolipidomic alterations.

As expected, analysis of the plasma lipidome revealed that long-chain ceramide species were significantly reduced in Myriocin-treated mice, reaching levels comparable to wild-type controls, whereas the plasma lipidomes of GW4869– and vehicle-treated mice remained indistinguishable (Fig. 4A-D, full annotation of 4C in Supplementary Table 4). Myriocin modestly attenuated body weight gain (Supplementary Fig. 2B) but did not lower plasma total cholesterol; plasma total cholesterol remained elevated across all *ApoE KO + PCSK9^DY^ AAV* treatment groups (Fig. 4D), which was also confirmed by ELISA-based LDL-cholesterol analysis (Fig. 4E). These data indicate that the protective effects of SPT inhibition occurred independently of systemic cholesterol lowering. Reduced plasma triglycerides and acylcarnitines may reflect secondary metabolic adaptation upon alleviation of ceramide overload and lipotoxic stress (Supplementary Fig. 2C). To further explore circulating acylcarnitines as indicators of impaired FAO in humans, we performed plasma lipidomics in a validation cohort of 70 non-diabetic CAVD patients without coronary artery disease to minimize confounders (Supplementary Table 5). All patients underwent transcatheter aortic valve implantation (TAVI) for severe aortic stenosis. The aortic valve calcification score, derived from CT scans, was correlated with plasma lipid species levels. Among all analyzed lipids, CAR 20:4 emerged as the only species significantly associated with the calcification score (Supplementary Fig. 3), supporting its role as an indicator of impaired fatty acid oxidation (FAO) and suggesting a potential link to subsequent pro-calcific lipid accumulation within the aortic valve

### Echocardiographic Assessment Reveals the Hemodynamic Benefits of Serine Palmitoyltransferase Inhibition Highlighting the Cross-Species Pathogenic Relevance of Cer 18:0;O3/23:0 in CAVD

Regarding echocardiographic parameters, SPT inhibition maintained aortic valve cusp separation, Vmax and dpmean near wild-type levels (Fig. 5A-D, Supplementary Video 1), whereas GW4869 treatment did not yield improvements in the severity of aortic valve stenosis. Echocardiographic assessment revealed no differences in left ventricular ejection fraction (LV-EF) between groups (Supplementary Fig. 2D). However, measures of diastolic dysfunction, including E/E’ ratio and isovolumic relaxation time (IVRT), were elevated in vehicle-treated *ApoE KO + PCSK9^DY^ AAV* mice compared to wild-type, indicating diastolic impairment in CAVD (Supplementary Fig. 2D). Notably, Myriocin normalized both E/E’ ratio and IVRT, whereas GW4869 had no effect, mirroring its lack of structural and hemodynamic benefits. To mechanistically link these functional improvements to sphingolipid modulation, we analyzed individual lipid species by loadings plot analysis identifying Cer 18:0;O3/23:0 as a principal driver of group variance (Fig.5E). Its accumulation in murine valve tissue strongly correlated with Vmax, even outperforming total Cer 18:0 species, while sphingomyelins showed weaker associations (Fig. 5F-H). These findings were further validated in human CAVD tissue, where Cer 18:0;O3/23:0 was highly upregulated and positively correlated with osteoprotegerin (TNFRSF11B) and TIMP1 (Fig. 2C). Overall, these data underscore the cross-species relevance of sphingolipid metabolism in aortic valve calcification and highlight that SPT inhibition improves hemodynamically relevant valve function independent of LDL-cholesterol lowering.

### Therapeutic SPT inhibition halts progression of established aortic stenosis

To model clinically relevant intervention after disease onset, *ApoE KO + PCSK9^DY^ AAV* mice were maintained on Western diet to establish aortic stenosis before treatment initiation (Fig. 5I). Echocardiography at week 17 confirmed increased aortic valve thickness and elevated transvalvular gradients (Fig. 5J-K, Supplementary Video 2), after which mice were randomized to receive Myriocin or vehicle for the remaining study period. Whereas vehicle-treated animals showed progressive worsening of valve hemodynamics, therapeutic SPT inhibition attenuated the progression of Vmax and dpmean and preserved cusp separation at week 24 (Fig. 5J-L, Supplementary Video 2). Endpoint histology indicated reduced leaflet calcification, fibrosis, and inflammatory cell infiltration in Myriocin-treated mice compared with vehicle (Fig. 5M). Body weight trajectories were not changed under Myriocin in the therapeutic approach (Supplementary Fig. 2E).

SPT inhibition prevents calcifying valve stenosis in a normolipidemic wire-injury model To disentangle valvular lipotoxic signaling from systemic hyperlipidemia and atherosclerosis-related confounding, we employed an aortic valve wire-injury model in wild-type mice maintained on normal chow (Supplementary Fig. 6A-B). Wire injury induced increased transvalvular gradients. Under normolipidemic conditions and without detectable differences in body weight, Myriocin treatment preserved valve hemodynamics (Supplementary Fig. 6D). Together with the therapeutic intervention study, these data demonstrate that the protective effects of SPT inhibition extend beyond preventive treatment and hyperlipidemic settings, supporting a direct role of ceramide-driven valve biology.

### Cellular Heterogeneity Uncovered by Single-Cell RNA Sequencing Reflects Anti-Calcifying Effect of Ceramide de novo Synthesis Inhibition

To investigate the cellular mechanisms linking ceramide-driven calcification and the therapeutic effects of ceramide de novo synthesis inhibition, we performed single-cell RNA sequencing (scRNA-seq) on murine aortic valve tissue from vehicle– and Myriocin-treated *ApoE KO + PCSK9^DY^ AAV* mice (n=4 per group). After quality filtering, 55,976 cells were classified into 13 clusters and visualized via Uniform Manifold Approximation and Projection (UMAP) (Fig. 6A-C). Clusters were grouped into five main cell lineages based on canonical marker gene expression: VICs, proliferative VICs, valvular endothelial cells (VECs), neutrophils, and other leukocytes (macrophages, dendritic cells, T cells, B cells). While all cell types were present in both *ApoE KO + PCSK9^DY^ AAV* groups (vehicle– and Myriocin-treated), notable compositional differences emerged (Fig. 6D). VICs were dominant, but Myriocin-treated mice exhibited a shift towards quiescent and myofibroblastic VICs, with reduced chondrogenic and osteogenic VICs, aligning with its anti-calcifying effects. Among leukocytes, macrophages were the most prevalent, with higher immune cell infiltration in vehicle-treated mice, supporting ceramide-driven inflammation in CAVD.

**Fig. 6.**
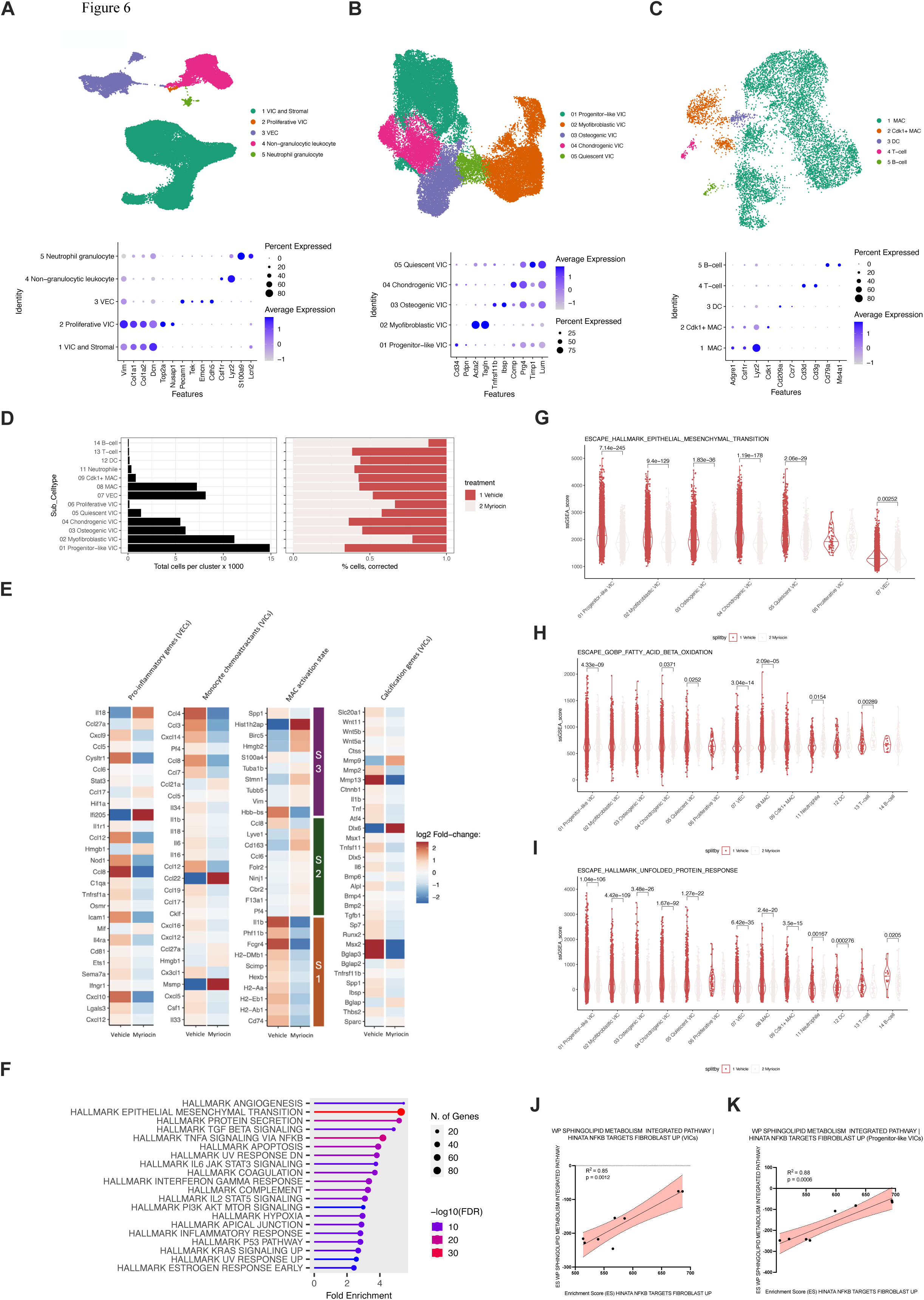
Single-cell RNA sequencing reveals anti-inflammatory, anti-EndMT, and ER-stress-attenuating effects of SPT inhibition in murine CAVD. A, UMAP of 55,976 aortic valve cells (n=4 mice per group) colored by major cell lineage with canonical-marker dot plot. B, UMAP of VIC subclusters with marker dot plot identifying progenitor-like, myofibroblastic, osteogenic, chondrogenic, and quiescent VIC states. C, UMAP of immune subclusters with marker dot plot identifying macrophages, proliferating macrophages, dendritic cells, T cells, and B cells. D, Absolute cell numbers and treatment-corrected cell fractions across subclusters. E, Heat maps showing selected pro-inflammatory genes in VECs, monocyte chemoattractant genes in VICs, macrophage activation-state genes, and calcification-related genes in VICs comparing vehicle and Myriocin. F, GSEA of differentially expressed genes in Myriocin-versus vehicle-treated valves highlights HALLMARK EPITHELIAL MESENCHYMAL TRANSITION and TNFA SIGNALING VIA NFKB pathways were most significantly enriched in vehicle-treated mice. G, ssGSEA scores for HALLMARK EPITHELIAL MESENCHYMAL TRANSITION. H, ssGSEA scores for fatty acid β-oxidation. I, ssGSEA scores for HALLMARK UNFOLDED PROTEIN RESPONSE. J-K, Correlation between WP sphingolipid metabolism and HINATA NF-κB targets fibroblast-up enrichment scores in total VICs and progenitor-like VICs. Two-sided Wilcoxon tests were used for ssGSEA comparisons; exact P values are indicated in the panels.

Further transcriptional analysis revealed that Myriocin treatment reduced pro-inflammatory gene expression in VECs (Fig. 6E, Supplementary Fig. 4A) and chemoattractant gene expression in VICs (Fig. 6E), consistent with lower macrophage infiltration. Additionally, Myriocin altered macrophage phenotype, reducing the pro-inflammatory Lyve1+MHC-II⁺⁺ subset (S1) while promoting an anti-inflammatory Lyve1+MHC-II⁻ subset (S2) [30] (Fig. 6E, Supplementary Fig. 4B). This was accompanied by the downregulation of calcification-related genes in VICs (Fig. 6E, Supplementary Fig. 4C). These findings underscore the interplay between ceramide accumulation, immune cell recruitment, and VIC differentiation in CAVD progression, highlighting the protective effects of SPT inhibition.

### Gene Set Enrichment Analysis Indicates Protection Against Valvular Endothelial to Mesenchymal Transition by Serine Palmitoyltransferase Inhibition

To elucidate the molecular mechanisms underlying SPT inhibition’s protective effects, we performed Gene Set Enrichment Analysis (GSEA) on scRNA-seq data from murine aortic valve tissue. The HALLMARK EPITHELIAL MESENCHYMAL TRANSITION and TNFA SIGNALING VIA NFKB pathways exhibited the highest –log10(FDR) values and the most enriched genes (Fig. 6F), mirroring key pathways distinguishing CAVD from control valves in human samples (Fig. 2). Myriocin treatment attenuated the enrichment of these pathways, indicating a therapeutic effect in counteracting central mechanisms of CAVD pathogenesis. As EndMT-transformed VECs serve as a precursor pool for activated VICs that further differentiate into osteogenic VICs [32], we analyzed single-sample GSEA (ssGSEA) scores for the HALLMARK EPITHELIAL MESENCHYMAL TRANSITION pathway. Myriocin-treated mice exhibited significant downregulation of EndMT scores in VECs (Fig. 6G), along with a marked reduction in EndMT enrichment in CD34-expressing progenitor-like VICs [33], underscoring Myriocin’s potential to mitigate EndMT-associated VIC activation in CAVD.

### SPT inhibition mitigates sphingolipid-driven lipotoxicity, ER stress and NF-κB-dependent VIC activation

The ssGSEA analysis further suggested an improvement in fatty acid β-oxidation in VECs following Myriocin treatment (Fig. 6H), consistent with FAO as an emerging regulator of EndMT progression [25]. Lipotoxicity has been linked to oxidative stress and ER stress in metabolic syndrome [34]. At single-cell resolution, Myriocin was associated with reduced enrichment of reactive oxygen species (ROS)-related pathways and attenuation of the unfolded protein response (UPR), a hallmark of ER stress, alongside decreased enrichment of NF-κB signaling pathways (Fig. 6I, Supplementary Fig. 4D). Notably, correlation analysis between enrichment scores for WP Sphingolipid Metabolism Integrated Pathway and HINATA NF-κB Targets Fibroblast Up revealed a strong positive association, particularly in progenitor-like VICs (Fig. 6J-K), underscoring the interplay between sphingolipid metabolism, NF-κB activation, and pathogenic VIC differentiation. Taken together, these findings support a model in which SPT inhibition disrupts ceramide-associated lipotoxic stress programs that amplify inflammatory activation and osteogenic differentiation of VICs.

### Long-chain ceramides induce ER stress and NF-κB-dependent calcification in human VICs

To experimentally validate the mechanistic programs in a human cell system, we employed primary human valvular interstitial cells (hVICs). In addition to short-chain C2 ceramide, we stimulated hVICs with a biologically relevant long-chain ceramide species (C18), reflecting the predominant ceramide signatures detected in calcified human valves. Both C2 and C18 ceramides promoted dose-dependent calcification, as quantified by calcium deposition and Alizarin Red staining, without evidence for cytotoxicity at concentrations up to 5 µM (Fig. 7A-E). Pro-calcific effects were compared by alkaline phosphatase (ALP) activity and qPCR analysis (Fig. 7F-G). Mechanistically, ceramide stimulation activated NF-κB signaling, with upregulation of key pathway components including IκB alpha, IKK gamma (NEMO), NFKB2, and Rela/p65, and increased IL-6 release (Fig. 7H-I). NF-κB inhibition by ammonium pyrrolidine dithiocarbamate (PDTC) and SC75741 abolished ceramide-induced calcification, identifying NF-κB as a key mediator of this process (Fig. 7J). Partial suppression of calcification with the NLRP3 inhibitor MCC950 suggests an additional contribution of inflammasome activation.

**Fig. 7.**
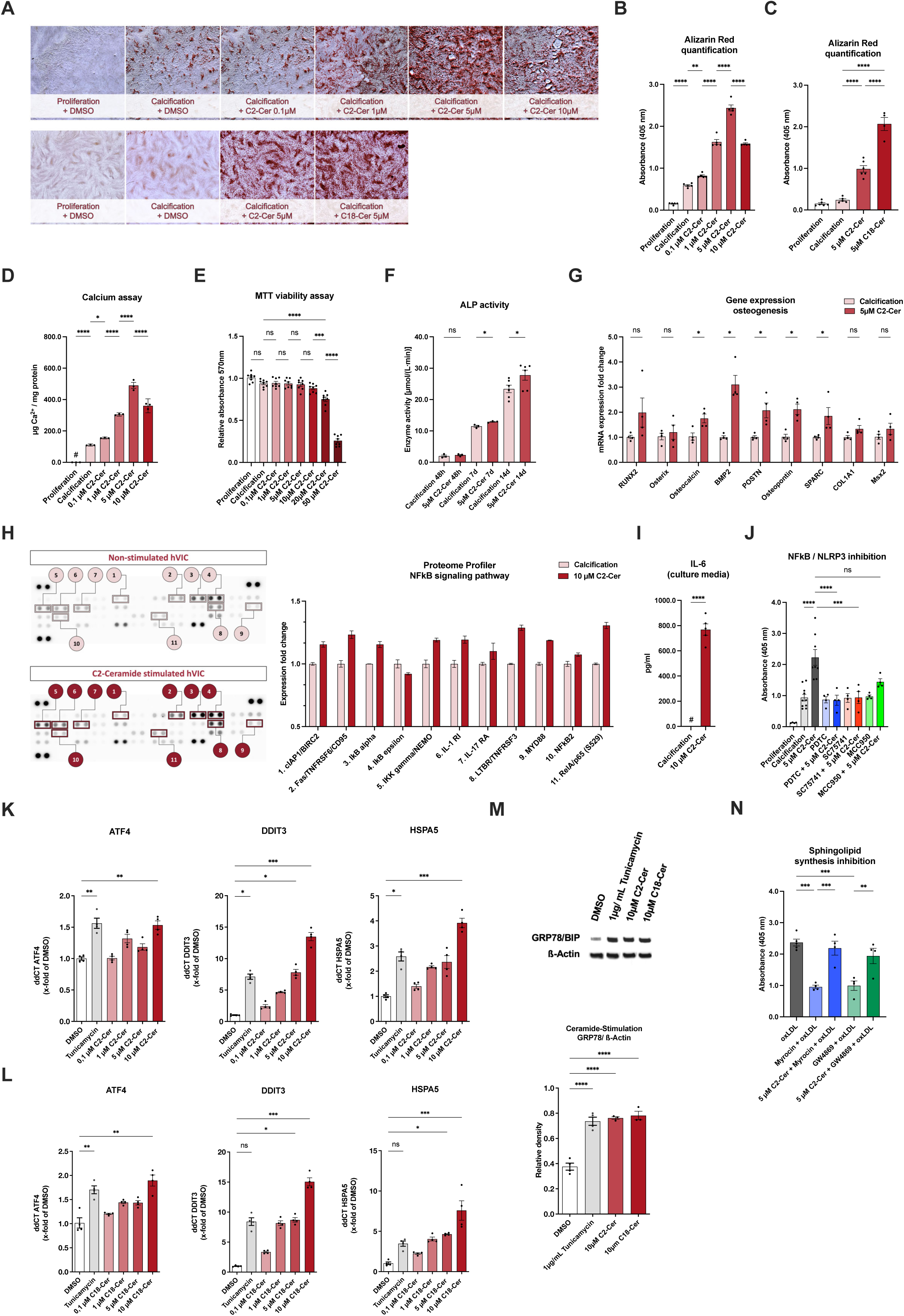
Short and long-chain ceramides induce ER stress and NF-κB-dependent calcification in human valvular interstitial cells. A, Representative Alizarin Red staining of hVICs cultured in proliferation or calcification medium with increasing C2 ceramide or 5 µM C18 ceramide. B, Quantification of Alizarin Red staining after graded C2 ceramide exposure. C, Quantification of Alizarin Red staining after 5 µM C2 or 5 µM C18 ceramide. D, Calcium assay after graded C2 ceramide exposure. E, MTT viability assay. F, Alkaline phosphatase (ALP) activity at 48 hours, 7 days, and 14 days in calcification medium with or without 5 µM C2 ceramide. G, Expression of osteogenic genes after 5 µM C2 ceramide. H, Representative Proteome Profiler membranes and densitometric quantification of selected NF-κB pathway proteins after 10 µM C2 ceramide. I, IL-6 concentration in culture medium after 10 µM C2 ceramide. J, Effects of PDTC, SC75741, and MCC950 on C2 ceramide-induced calcification. K, UPR gene expression after C2 ceramide stimulation. L, UPR gene expression after C18 ceramide stimulation. M, Representative GRP78/BiP immunoblot and quantification after DMSO, Tunicamycin, C2 ceramide, or C18 ceramide. N, OxLDL-induced calcification with rescue experiments using Myriocin or GW4869 with or without C2 ceramide reconstitution. All bar charts show mean ± SEM. Statistical analyses were performed by 1-way ANOVA followed by Bonferroni post hoc correction (B-F, J-N) or Student’s t test (G, I). *P<0.05, **P<0.01, ***P<0.001, ****P<0.0001.

Consistent with the cross-species enrichment of unfolded protein response/ ER-stress pathways, ceramide exposure induced ER stress in hVICs, evidenced by increased GRP78/BiP protein abundance and dose-dependent induction of canonical UPR transcripts including CHOP, GRP78, and ATF4 (Fig. 7K-M). Finally, pharmacologic inhibition of de novo ceramide synthesis attenuated endogenous ceramide-dependent calcification (Fig. 7N), supporting ceramide accumulation as a translatable actionable upstream driver of pro-inflammatory and pro-osteogenic VIC activation in human derived cells.

## Discussion

Here we define ceramide-driven lipotoxicity as a cholesterol-independent upstream mechanism of CAVD. Integrating lipidomics and proteomics of human aortic valve tissue with single-cell transcriptomics, mechanistic experiments in primary hVICs, and complementary *in vivo* models, we show that long-chain ceramides accumulate locally in calcified valves and align with NF-κB activation, EndMT, and ER stress programs. Pharmacologic inhibition of SPT with Myriocin reduced valvular ceramide burden and mitigated fibro-calcific remodeling and hemodynamically relevant stenosis in preventive and therapeutic settings without lowering plasma LDL-cholesterol, including in a normolipidemic wire-injury model. These findings provide mechanistic and translational rationale for targeting sphingolipid biosynthesis as a disease-modifying strategy in CAVD.

CAVD is prevalent and lethal once advanced, yet pharmacologic disease modification remains elusive. Trials of LDL-cholesterol lowering failed to slow aortic stenosis progression [3, 4] [35], suggesting that LDL-centric paradigms incompletely capture key drivers of fibro-calcific remodeling. The valve leaflet is a bradytrophic, largely avascular tissue [16, 36], which likely shapes how metabolic stressors translate into chronic inflammation, EndMT, and osteogenic VIC differentiation.

Sphingolipids, particularly ceramides, are established mediators of cardiometabolic inflammation and vascular disease [5–8, 37]. Extending prior descriptive studies of spatial lipid distribution in stenotic valves [17], we compared entirely non-calcified control valves with fully calcified CAVD specimens and integrated lipidomics with unbiased proteomics. This approach revealed a pronounced local long-chain ceramide signature in CAVD tissue, together with depletion of triglycerides and increases in fatty acids and acylcarnitines, consistent with local lipotoxic pressure in the setting of systemic FAO stress. Notably, plasma sphingolipid classes were not increased, suggesting that valve ceramide accumulation reflects local tissue remodeling rather than generalized circulating ceramide elevation.

Across omics layers, ceramide enrichment aligned with NF-κB signaling, EndMT, and ER-stress/UPR programs. Mechanistic experiments in hVICs support causality: long-chain C18 ceramide, reflecting species accumulating in human valves, activated the UPR and promoted calcification, while NF-κB inhibition abrogated ceramide-induced calcification. Single-cell RNA-seq connected sphingolipid metabolism to inflammatory and stress programs at cellular resolution, with SPT inhibition attenuating TNFα/NF-κB, EndMT, UPR, and ROS-associated signatures and shifting VIC states away from osteogenic phenotypes.

To test therapeutic tractability, we targeted SPT, a druggable rate-limiting step of de novo sphingolipid biosynthesis. In the *ApoE KO + PCSK9^DY^ AAV* model, Myriocin normalized the valvular sphingolipidome, reduced calcification, fibrosis, and macrophage infiltration, and improved hemodynamically relevant stenosis while leaving plasma LDL-cholesterol elevated—demonstrating cholesterol-independent valve protection. In contrast, inhibition of neutral sphingomyelinase with GW4869 failed to reduce valvular ceramide burden or improve stenosis, consistent with its limited systemic effect and the restricted availability of circulating sphingomyelin substrate within the bradytrophic valve milieu [36].

Importantly, the therapeutic cohort moves the study beyond prevention. To our knowledge, this is the first demonstration that pharmacological inhibition of sphingolipid metabolism, initiated only after echocardiographically verified elevation of transvalvular gradients, attenuates further hemodynamic progression of calcific aortic valve stenosis *in vivo*. This distinction is critical because most experimental CAVD interventions are tested prophylactically, whereas patients present after valve gradients have already risen. In parallel, in normolipidemic wild-type mice maintained on chow, Myriocin prevented wire-injury–induced calcifying stenosis, indicating that its protective effects are not restricted to hypercholesterolemic or diet-driven disease contexts. Together with the unchanged LDL-cholesterol levels observed in the Myriocin treated *ApoE KO + PCSK9^DY^ AAV* model, these findings support a mechanism that extends beyond systemic cholesterol lowering. Direct leaflet trauma is known to elicit a robust fibro-inflammatory response characterized by oxidative stress, cytokine induction, and osteochondrogenic signalling that culminates in calcification and stenosis [21]. Pattern-recognition receptor signalling can intersect with saturated fatty-acid availability to amplify de novo ceramide synthesis and downstream inflammatory programs [38], providing a plausible mechanistic basis for SPT inhibition even in the absence of general hyperlipidemia.

Our pharmacologic findings extend recent genetic work implicating ceramide synthase 5 (CerS5) in wire-injury–induced aortic valve stenosis under Western diet [39]. Whereas CerS5 deficiency targets a single ceramide synthase and demonstrated benefit predominantly in the context of diet-induced lipotoxic stress, our study integrates human multi-omics data, identifies a conserved long-chain ceramide correlate (Cer 18:0;O3/23:0) across species, and demonstrates that upstream inhibition of sphingolipid de novo synthesis confers protection in preventive and therapeutic settings as well as in normolipidemic injury-driven disease.

Furthermore, our findings complement recent soluble guanylyl cyclase (sGC)-reactivation work with ataciguat, which provided preclinical and early clinical evidence for slowing fibrocalcific aortic valve stenosis progression [40], but identify a distinct upstream metabolic mechanism: valve-local de novo sphingolipid synthesis and ceramide-driven lipotoxicity.

However, several limitations warrant consideration. Human samples were derived from end-stage disease requiring surgery, precluding direct inference on when ceramides begin to accumulate during earlier stages of CAVD. *In vivo*, Myriocin was administered systemically and modestly attenuated weight gain in the preventive hyperlipidemic cohort; in the therapeutic cohort (Myriocin initiated at week 17), endpoint body weight was comparable between Myriocin-and vehicle-treated mice despite marked differences in transvalvular gradients. LDL-cholesterol was unchanged and valve protection was also observed in the wire-injury model without weight differences, making body weight an unlikely sole driver of benefit. Nevertheless, future studies leveraging tissue– and cell type-specific genetic perturbations, including conditional knockout models, are needed to delineate the most relevant sources and targets of ceramide accumulation within the aortic valve niche. Pharmacological translation of Myriocin is challenged by its potential gastrointestinal toxicity in humans [41], highlighting the need for therapeutic alternatives. Recently, a safe and orally bioavailable SPT inhibitor (ALT-007) was reported [42], and emerging targeted delivery platforms –including lipid nanoparticles [43], AAVs [44], or magnetic nanocarriers [45] –may enable organ– or cell-selective modulation of ceramide metabolism, potentially minimizing off-target effects.

In conclusion, our integrative multi-omics and reverse-translational data define a cholesterol-independent ceramide–ER stress–NF-κB–EndMT axis, demonstrate direct ceramide-driven calcification in human VICs with physiologic long-chain species, and provide therapeutic proof-of-concept across complementary *in vivo* models, including an established disease stage. Together, these findings position ceramide metabolism as a tractable upstream driver of valve calcification and support targeting sphingolipid metabolism as a potential disease-modifying strategy in CAVD

## Conflicts of interests

S.A.S.: Founder and shareholder of Centaurus Therapeutics.

T.G.: Member of the BigOmics Analytics Advisory Board. E.R.: None

T.R.: None

J.I.: None

K.H.: None

C.G.: None

J.C.V.: None

M.S.: None

K.K.: None

M.B.: None

S.D.: None

A.K.: None

T.Sa.: None

T.Sc.: None

D.F.: None

T.St.: None

W.R.: None

O.J.M.: None

T.Z.: None

H.B.: None

C.K.: None

M.Sp.: None

Z.A.: None

I.E.: None

C.F.: None

## Acknowledgments

We thank Anke Constantz, Annett Liebers, Julia Horn and Alexander Haschke for their excellent technical assistance. We thank Thomas Eschenhagen for proofreading the revised manuscript.

## Non-standard abbreviations and acronyms

ALP: Alkaline Phosphatase
ApoE KO: Apolipoprotein E Knockout
AAV: Adeno-Associated Virus
CAR: Acylcarnitine
CAVD: Calcific Aortic Valve Disease
CE: Cholesterol Ester
Cer: Ceramide
CT: Computed Tomography
dpmean: Mean Pressure Gradient
ECM: Extracellular Matrix
EndMT: Endothelial-to-Mesenchymal Transition
ER: Endoplasmic Reticulum
FA: Fatty Acid
FAO: Fatty Acid Oxidation
FC: Fold Change
hVICs: Human Valvular Interstitial Cells
HexCer: Hexosylceramide
IVRT: Isovolumic Relaxation Time
LD: Lipid Droplets
LC-MS: Liquid Chromatography Mass Spectrometry
LPC O: Ether-linked lysophosphatidylcholines
LV-EF: Left Ventricular Ejection Fraction
MALDI-MSI: Matrix-assisted laser desorption/ionization mass spectrometry imaging
ME: Module Eigenprotein
NF-κB: Nuclear Factor kappa-light-chain-enhancer of activated B cells
PLIN1: Perilipin-1
RANKL: Receptor Activator of Nuclear Factor Kappa-Β Ligand
ROS: Reactive Oxygen Species
sPLS-DA: Sparse Partial Least Squares Discriminant Analysis
SM: Sphingomyelin
TG: Triglyceride
TLR2: Toll-like Receptor 2
TAVI: Transcatheter Aortic Valve Implantation
UPR: Unfolded Protein Response
VECs: Valvular Endothelial Cells
Vmax: Maximum Velocity
VICs: Valvular Interstitial Cells
WD: Western Diet
WGCNA: Weighted Gene Co-expression Network Analysis

---

**Supplementary Fig. 1. Network topology, module-trait relationships, and partial-correlation networks from valve proteomics.** A, Scale independence and mean connectivity analysis of network topology for various soft-thresholding powers used for weighted gene co-expression network analysis (WGCNA). B, Protein dendrogram and topological overlap matrix (TOM) heatmap. C, Module-trait relationship heat map for ME1-ME7 in control and CAVD samples. D-F, Enrichment analysis of ME1, ME3, and ME4, respectively. G-I, Partial correlation graphs for ME1, ME3, and ME4. Green edges correspond to positive partial correlation and red edges to negative partial correlation; edge width is proportional to correlation strength.

**Supplementary Fig. 2. PCSK9 transduction, ancillary metabolic changes, and cardiac functional parameters in the preventive and therapeutic mouse studies.** A, Plasma PCSK9 levels after 21 weeks confirm successful PCSK9^DY^-AAV transduction. B, Body-weight gain in the preventive hyperlipidemic cohort. C, Plasma triglycerides (TG), fatty acids (FA), and acylcarnitines (CAR) in the preventive cohort. D, Heart rate, left ventricular ejection fraction (LV-EF), E/E’, and isovolumic relaxation time (IVRT) in the preventive cohort. E, Body-weight gain in the therapeutic cohort. F, Heart rate, LV-EF, E/E’, and IVRT in the therapeutic cohort. Bar charts show mean ± SEM. Statistical analyses were performed by 1-way ANOVA followed by Bonferroni post hoc correction. *P<0.05, **P<0.01, ***P<0.001, ****P<0.0001.

**Supplementary Fig. 3. Plasma CAR 20:4 correlates with CT-derived valve calcification in the TAVI validation cohort.** A, Correlation between plasma CAR 20:4 and computed tomography–derived aortic valve calcification score in 70 non-diabetic TAVI patients without coronary artery disease.

**Supplementary Fig. 4. Myriocin suppresses inflammatory, macrophage-activation, calcification, and stress signatures in scRNA-seq data.** A, Violin plot of selected pro-inflammatory genes in valvular endothelial cells (VECs). B, Violin plot of the pro-inflammatory S1 macrophage activation-state score. C, Violin plot of calcification-related genes in VICs. D, Heat map of pathway-enrichment scores across the major valve cell populations for sphingolipid metabolism, NF-κB signaling, unfolded protein response, inflammatory response, reactive oxygen species, ceramide pathway activity, and fatty acid β-oxidation.

**Supplementary Fig. 5. Full immunoblot of GRP78/BiP after ceramide stimulation in hVICs.** A, Full Western blot for GRP78/BiP and β-actin in DMSO-treated cells and cells exposed to Tunicamycin, C2 ceramide, or C18 ceramide.

**Supplementary Fig. 6. Myriocin prevents wire-injury-induced aortic valve stenosis under normolipidemic conditions.** A, Study design for the wild-type wire-injury model. B, Exemplary echocardiographic image showing wire position relative to the left ventricle (LV) and aortic valve (AV). C, Dpmean in uninjured and wire-injured mice treated with vehicle or Myriocin for 6 weeks. Bar charts show mean ± SEM. Statistical analyses were performed by 1-way ANOVA followed by Bonferroni post hoc correction. ****P<0.0001.

## Notes

### Funding

- German Centre for Cardiovascular Research: 81X3700107, 81X1700114, 81Z0700107
- Horizon Europe: Grant No. 101156555
- University of Lübeck: CS06-2024

## References

1. Coffey, S., B.J. Cairns, and B. Iung, The modern epidemiology of heart valve disease. Heart, 2016. 102(1): p. 75–85.

2. Gupta, A., T. Liu, C. Pounds, R.P. Sharma, C. Yong, H.M. Krumholz, et al., Predictors of increased mortality in untreated moderate aortic stenosis. European Heart Journal, 2021. 42(Supplement_1): p. ehab724.1573.

3. Chan, K.L., K. Teo, J.G. Dumesnil, A. Ni, and J. Tam, Effect of Lipid lowering with rosuvastatin on progression of aortic stenosis: results of the aortic stenosis progression observation: measuring effects of rosuvastatin (ASTRONOMER) trial. Circulation, 2010. 121(2): p. 306–14.

4. Rossebø, A.B., T.R. Pedersen, K. Boman, P. Brudi, J.B. Chambers, K. Egstrup, et al., Intensive lipid lowering with simvastatin and ezetimibe in aortic stenosis. N Engl J Med, 2008. 359(13): p. 1343–56.

5. Choi, R.H., S.M. Tatum, J.D. Symons, S.A. Summers, and W.L. Holland, Ceramides and other sphingolipids as drivers of cardiovascular disease. Nat Rev Cardiol, 2021. 18(10): p. 701–711.

6. Hannun, Y.A. and L.M. Obeid, Sphingolipids and their metabolism in physiology and disease. Nat Rev Mol Cell Biol, 2018. 19(3): p. 175–191.

7. Rawish, E., L. Nickel, F. Schuster, I. Stölting, A. Frydrychowicz, K. Saar, et al., Telmisartan prevents development of obesity and normalizes hypothalamic lipid droplets. J Endocrinol, 2020. 244(1): p. 95–110.

8. Edsfeldt, A., P. Dunér, M. Ståhlman, I.G. Mollet, G. Asciutto, H. Grufman, et al., *Sphingolipids Contribute to Human Atherosclerotic Plaque Inflammation.* Arteriosclerosis, Thrombosis, and Vascular Biology, 2016. 36(6): p. 1132–1140.

9. Hojjati, M.R., Z. Li, H. Zhou, S. Tang, C. Huan, E. Ooi, et al., Effect of myriocin on plasma sphingolipid metabolism and atherosclerosis in apoE-deficient mice. J Biol Chem, 2005. 280(11): p. 10284–9.

10. Yu, Z., Q. Peng, S. Li, H. Hao, J. Deng, L. Meng, et al., Myriocin and d-PDMP ameliorate atherosclerosis in ApoE−/− mice via reducing lipid uptake and vascular inflammation. Clinical Science, 2020. 134(5): p. 439–458.

11. Glaros, E.N., W.S. Kim, C.M. Quinn, W. Jessup, K.A. Rye, and B. Garner, Myriocin slows the progression of established atherosclerotic lesions in apolipoprotein E gene knockout mice. J Lipid Res, 2008. 49(2): p. 324–31.

12. Zadelaar, S., R. Kleemann, L. Verschuren, J. de Vries-Van der Weij, J. van der Hoorn, H.M. Princen, et al., *Mouse Models for Atherosclerosis and Pharmaceutical Modifiers.* Arteriosclerosis, Thrombosis, and Vascular Biology, 2007. 27(8): p. 1706–1721.

13. Isoda, K., T. Matsuki, H. Kondo, Y. Iwakura, and F. Ohsuzu, Deficiency of interleukin-1 receptor antagonist induces aortic valve disease in BALB/c mice. Arterioscler Thromb Vasc Biol, 2010. 30(4): p. 708–15.

14. Sider, K.L., M.C. Blaser, and C.A. Simmons, Animal models of calcific aortic valve disease. Int J Inflam, 2011. 2011: p. 364310.

15. Hulin, A., A. Hego, P. Lancellotti, and C. Oury, Advances in Pathophysiology of Calcific Aortic Valve Disease Propose Novel Molecular Therapeutic Targets. Front Cardiovasc Med, 2018. 5: p. 21.

16. Moncla, L.-H.M., M. Briend, Y. Bossé, and P. Mathieu, Calcific aortic valve disease: mechanisms, prevention and treatment. Nature Reviews Cardiology, 2023. 20(8): p. 546–559.

17. Prabutzki, P., M. Wölk, J. Böttner, Z. Ni, S. Werner, H. Thiele, et al., Sex-specific lipidomic signatures in aortic valve disease reflect differential fibro-calcific progression. Nat Commun, 2025. 16(1): p. 5163.

18. Lehti, S., R. Käkelä, S. Hörkkö, O. Kummu, S. Helske-Suihko, M. Kupari, et al., Modified lipoprotein-derived lipid particles accumulate in human stenotic aortic valves. PLoS One, 2013. 8(6): p. e65810.

19. Achner, L., T. Klersy, B. Fels, T. Reinberger, C.X. Schmidt, N. Groß, et al., AFM-based nanoindentation indicates an impaired cortical stiffness in the AAV-PCSK9(DY) atherosclerosis mouse model. Pflugers Arch, 2022. 474(9): p. 993–1002.

20. Wu, Q., L. Sun, X. Hu, X. Wang, F. Xu, B. Chen, et al., Suppressing the intestinal farnesoid X receptor/sphingomyelin phosphodiesterase 3 axis decreases atherosclerosis. J Clin Invest, 2021. 131(9).

21. Honda, S., T. Miyamoto, T. Watanabe, T. Narumi, S. Kadowaki, Y. Honda, et al., A novel mouse model of aortic valve stenosis induced by direct wire injury. Arterioscler Thromb Vasc Biol, 2014. 34(2): p. 270–8.

22. Makrecka-Kuka, M., E. Sevostjanovs, K. Vilks, K. Volska, U. Antone, J. Kuka, et al., Plasma acylcarnitine concentrations reflect the acylcarnitine profile in cardiac tissues. Scientific Reports, 2017. 7(1): p. 17528.

23. Kondo, T., T. Matsuda, M. Tashima, H. Umehara, N. Domae, K. Yokoyama, et al., Suppression of Heat Shock Protein-70 by Ceramide in Heat Shock-induced HL-60 Cell Apoptosis*. Journal of Biological Chemistry, 2000. 275(12): p. 8872–8879.

24. Xu, K., S. Xie, Y. Huang, T. Zhou, M. Liu, P. Zhu, et al., *Cell-Type Transcriptome Atlas of Human Aortic Valves Reveal Cell Heterogeneity and Endothelial to Mesenchymal Transition Involved in Calcific Aortic Valve Disease.* Arteriosclerosis, Thrombosis, and Vascular Biology, 2020. 40(12): p. 2910–2921.

25. Xiong, J., H. Kawagishi, Y. Yan, J. Liu, Q.S. Wells, L.R. Edmunds, et al., A Metabolic Basis for Endothelial-to-Mesenchymal Transition. Molecular Cell, 2018. 69(4): p. 689–698.e7.

26. Sztalryd, C. and D.L. Brasaemle, The perilipin family of lipid droplet proteins: Gatekeepers of intracellular lipolysis. Biochim Biophys Acta Mol Cell Biol Lipids, 2017. 1862(10 Pt B): p. 1221–1232.

27. Huang, S., J.M. Rutkowsky, R.G. Snodgrass, K.D. Ono-Moore, D.A. Schneider, J.W. Newman, et al., Saturated fatty acids activate TLR-mediated proinflammatory signaling pathways[S]. Journal of Lipid Research, 2012. 53(9): p. 2002–2013.

28. Serbulea, V., C.M. Upchurch, K.W. Ahern, G. Bories, P. Voigt, D.E. DeWeese, et al., Macrophages sensing oxidized DAMPs reprogram their metabolism to support redox homeostasis and inflammation through a TLR2-Syk-ceramide dependent mechanism. Mol Metab, 2018. 7: p. 23–34.

29. Fucho, R., N. Casals, D. Serra, and L. Herrero, Ceramides and mitochondrial fatty acid oxidation in obesity. Faseb j, 2017. 31(4): p. 1263–1272.

30. Lee, S.H., N. Kim, M. Kim, S.-H. Woo, I. Han, J. Park, et al., Single-cell transcriptomics reveal cellular diversity of aortic valve and the immunomodulation by PPARγ during hyperlipidemia. Nature Communications, 2022. 13(1): p. 5461.

31. Spaggiari, R., S. Angelini, A. Di Vincenzo, G. Scaglione, S. Morrone, V. Finello, et al., Ceramides as Emerging Players in Cardiovascular Disease: Focus on Their Pathogenetic Effects and Regulation by Diet. Advances in Nutrition, 2024. 15(7): p. 100252.

32. Ma, X., D. Zhao, P. Yuan, J. Li, Y. Yun, Y. Cui, et al., Endothelial-to-Mesenchymal Transition in Calcific Aortic Valve Disease. Acta Cardiol Sin, 2020. 36(3): p. 183–194.

33. Decano, J.L., Y. Iwamoto, S. Goto, J.Y. Lee, J.T. Matamalas, A. Halu, et al., A disease-driver population within interstitial cells of human calcific aortic valves identified via single-cell and proteomic profiling. Cell Reports, 2022. 39(2).

34. Chaurasia, B. and S.A. Summers, Ceramides in Metabolism: Key Lipotoxic Players. Annu Rev Physiol, 2021. 83: p. 303–330.

35. Zhao, Y., R. Nicoll, Y.h. He, and M.Y. Henein, The effect of statins on valve function and calcification in aortic stenosis: A meta-analysis. Atherosclerosis, 2016. 246: p. 318–324.

36. Miller, J.D., R.M. Weiss, D.D. Heistad, and D.A. Towler, Calcific Aortic Valve Stenosis: Methods, Models, and Mechanisms. Circulation Research, 2011. 108(11): p. 1392–1412.

37. Bismuth, J., P. Lin, Q. Yao, and C. Chen, Ceramide: a common pathway for atherosclerosis? Atherosclerosis, 2008. 196(2): p. 497–504.

38. Schilling, J.D., H.M. Machkovech, L. He, R. Sidhu, H. Fujiwara, K. Weber, et al., Palmitate and lipopolysaccharide trigger synergistic ceramide production in primary macrophages. J Biol Chem, 2013. 288(5): p. 2923–32.

39. Reese, L., S.T. Niepmann, P. Düsing, L. Hänschke, T. Beiert, S. Zimmer, et al., Loss of ceramide synthase 5 inhibits the development of experimentally induced aortic valve stenosis. Acta Physiol (Oxf), 2024. 240(5): p. e14140.

40. Zhang, B., M. Enriquez-Sarano, H.V. Schaff, H.I. Michelena, C.M. Roos, M.A. Hagler, et al., Reactivation of Oxidized Soluble Guanylate Cyclase as a Novel Treatment Strategy to Slow Progression of Calcific Aortic Valve Stenosis: Preclinical and Randomized Clinical Trials to Assess Safety and Efficacy. Circulation, 2025. 151(13): p. 913–930.

41. Monasterio, B.G., N. Jiménez-Rojo, A.B. García-Arribas, H. Riezman, F.M. Goñi, and A. Alonso, Plasma membrane effects of sphingolipid-synthesis inhibition by myriocin in CHO cells: a biophysical and lipidomic study. Sci Rep, 2022. 12(1): p. 955.

42. Poisson, J., I. Daskalaki, V. Potluri, J.-D. Morel, S. Rodriguez-Lopez, A. De Masi, et al., Safe and Orally Bioavailable Inhibitor of Serine Palmitoyltransferase Improves Age-Related Sarcopenia. ACS Pharmacology & Translational Science, 2025. 8(1): p. 203–215.

43. Labonia, M.C.I., M. Estape Senti, P.H. Van Der Kraak, M.A.D. Brans, A.K. Deshantri, I. Dokter, et al., Effective cardiac mRNA delivery using lipid nanoparticles. European Heart Journal, 2023. 44(Supplement_2): p. ehad655.3301.

44. Wong, F.F., M.L. Ho, M. Yamagami, M.T. Lam, K.J. Grande-Allen, and J. Suh, Effective Gene Delivery to Valvular Interstitial Cells Using Adeno-Associated Virus Serotypes 2 and 3. Tissue Eng Part C Methods, 2015. 21(8): p. 808–15.

45. Chen, J., T. Ren, L. Xie, H. Hu, X. Li, M. Maitusong, et al., Enhancing aortic valve drug delivery with PAR2-targeting magnetic nano-cargoes for calcification alleviation. Nature Communications, 2024. 15(1): p. 557.

