## Supplemental Materials and Figures for "Sphingolipid-Driven Lipotoxicity Promotes Calcific Aortic Valve Disease Through ER-Stress–NF-κB–EndMT Signaling and Is Therapeutically Targetable Through Serine Palmitoyltransferase Inhibition"

#### Extended Methods

##### Animal Studies

All animal care and experimental procedures were performed in accordance with the NIH guidelines for the care and use of laboratory animals and were approved by the animal ethics committee of the local regulatory authority (Reference Number 49-9/23, Ministerium für Landwirtschaft, ländliche Räume, Europa und Verbraucherschutz des Landes Schleswig-Holstein). The animal studies are reported according to the ARRIVE guidelines.

B6.129P2-Apoetm1Unc/J (ApoE KO) mice were obtained from Charles River Laboratories (Sulzfeld, Germany). To enhance hypercholesterolemia and valvular calcification, promoting CAVD, adeno-associated virus (AAV) encoding $PCSK9^{DY}$ (Proprotein convertase subtilisin/kexin type 9) was administered at the age of seven weeks via tail vein injection at a dose of $2x10^{11}$ viral particles per mouse (N=44) [1]. Sufficient $PCSK9^{DY}$ AAV injection was later confirmed by plasma PCSK9 measurement (Supplementary Fig. 1A). C57BL/6J mice served as non-diseased controls (N=14). Male and female mice were fed a cholesterol-enriched Western Diet (EF acc. D12108) to accelerate CAVD. Before initiation of Western Diet and $PCSK9^{DY}$-AAV injection, baseline blood samples were collected by tail vein puncture for lipidomic analysis. Throughout the study, mice were treated chronically for a total of 21 weeks with either vehicle, Myriocin (0.3 mg/kg/d), or GW4869 (30 mg/kg/d) [2] dissolved in drinking water. The concentrations were adjusted based on continuous monitoring of water consumption to ensure accurate and consistent delivery of the compounds. For therapeutic intervention, an independent ApoE KO + PCSK9$^{DY}$-AAV cohort was maintained on Western diet for 24 weeks; after echocardiography at week 17 to document established transvalvular gradients, mice were randomized to receive Myriocin or vehicle for the remaining 7 weeks (weeks 17–24). At the end of each experiment, terminal blood samples (500 µl) were collected under anesthesia using ketamine (100 mg/kg, i.p.) and xylazine (10 mg/kg i.p.) via the retro-orbital venous plexus, and plasma was prepared for further analyses.

#### Wire Injury Model in Wild-Type Mice

Aortic valve wire injury was performed in wild-type C57BL/6J mice (Charles River Laboratories, Sulzfeld, Germany) maintained on standard chow to induce fibro-calcific aortic valve disease under normolipidemic conditions, following established protocols [3]. Mice were randomized to receive Myriocin (0.3 mg/kg/day) or vehicle in drinking water for the duration of the experiment, with concentrations adjusted according to water consumption. Echocardiography was performed at baseline and predefined follow-up time points to assess transvalvular gradients and valve morphology. At endpoint, valves and blood were collected for histological and molecular analyses.

#### Mouse Echocardiography

To assess the hemodynamic impact of aortic valve disease, transthoracic echocardiography was performed longitudinally in all mouse cohorts. In the preventive metabolic model cohort, imaging was performed in week 21. In the therapeutic cohort, echocardiography was performed at week 17 prior to treatment randomization and at week 24 at endpoint. In the wire-injury cohort, echocardiography was performed at baseline and six weeks post-injury. Sedation was achieved with an intraperitoneal injection of 100 mg/kg ketamine dissolved in 0.9% saline, minimizing movement artifacts during imaging [4]. Each mouse was positioned on a heated stage (37°C), heart rate and electrocardiogram (ECG) were continuously monitored. The chest area was shaved using depilatory cream to facilitate ultrasound imaging. Echocardiography was conducted using a GE i13L probe on a GE Vivid E9 system, capturing standard views, including parasternal long-axis and short-axis views, as well as the three-chamber view for additional gradient measurements across the aortic valve. Key parameters evaluated included peak transvalvular velocity (Vmax), mean pressure gradient (dpmean) across the aortic valve, and mean aortic valve thickness in the parasternal long axis. In addition, systolic and diastolic markers of left ventricular function were evaluated (including left ventricular ejection fraction, LVEF; E/E'; and isovolumic relaxation time, IVRT).

#### Single-cell RNA Sequencing

Following euthanasia, aortic valves were harvested from the mice (n=4 mice per group). Each valve was processed into a single-cell suspension by enzymatic digestion. The digestion was performed at 37°C for 45 minutes using a rotating incubator with $Ca^{2+}/Mg^{2+}$ Dulbecco's phosphate-buffered saline ($Ca^{2+}/Mg^{2+}$ DPBS, Welgene, #LB 001-01) containing collagenase II (1000 U/mL, Sigma-Aldrich, #C6885) and DNase I (90 U/mL, Roche, #10104159001) [5]. The resulting single-cell suspension was filtered through a 40 µm cell strainer to remove debris and large aggregates. Cell viability was assessed, and cells were quantified in a Neubauer chamber with Trypan blue staining. Subsequent single-cell RNA sequencing (scRNA-seq) library prep was performed using the Chromium GEM-X Single Cell 3' v4 Gene Expression kit on a Chromium X device (10X Genomics) following the manufacturer's protocols. Libraries were quantified, and library fragment size distribution was determined using a Qubit 1x dsDNA HS assay (Invitrogen) and an Agilent High Sensitivity DNA Kit on a Bioanalyzer 2100 device (Agilent), respectively.

Libraries were multiplexed and sequenced using XLEAP-SBS P4 (100 Cycles) reagents on a NextSeq 2000 device (Illumina). Sequencing data was demultiplexed and converted to FASTQ format using bcl2fastq2 v2.20 (Illumina), and count matrices were generated with Cellranger v7.0.0 (10X Genomics) using the mouse reference transcriptome mm10-2020-A (10x Genomics). The rest of the analysis used Seurat v4 [6] and standard R packages. Custom filtering at the cell level was performed with a minimum of 400 UMI counts and between 200 and 2,000 distinct features (genes), a maximum of 25% mitochondrial and 25% ribosomal reads, as well as a doublet score (scrublet-0.2.3 [7]) below 0.22. Individual samples were integrated using harmony [8]. Cluster annotation was performed based on differentially expressed marker genes from published mouse aortic valve data [5] (Fig. 6A-D). Clusters expressing markers Syt1 & Kcnip4 (neural cell contamination from sympathetic nervous system of the aorta), Myl2 and Tnnt2 (contaminating cardiomyocytes), Adipoq (contaminating Adipocytes), Wt1 and Upk3b (epicardial cells), and Plac8 and Ly6c2 (monocytes) were removed before UMAP projection (Fig. 6 A-D) and ignored for subsequent analysis.

Differentially expressed genes between Myriocin treated and vehicle samples were used as an input for GSEA for the Hallmark gene sets [9] using ShinyGO [10] (Fig. 6F). Single-sample Gene Set Enrichment Analysis (ssGSEA) was applied to scRNA-seq data using the escape R-package [11]. The msigdbr and getGeneSets functions were used to fetch and filter the corresponding gene sets of Mus musculus from the MSigDB [9, 12]. EnrichIt with default parameters, except for using 10,000 groups and a variable number of cores, was performed on the Seurat object (Supplementary Fig. 4D). A double-sided Wilcoxon test (rstatix::wilcox_test) with default parameters was applied to statistically compare scores between cells treated with Myriocin or Vehicle (Fig. 6E, G-J).

#### Histology

Following euthanasia, aortic valves were excised and fixed in 4% formaldehyde for 24 hours at 4°C followed by a second fixation step for 24 h in Tissue Tek (Sakura). After fixation, tissues were dehydrated in a graded ethanol series, embedded in paraffin, and sectioned into 5 µm thick slices using a kryotome at -20°C. The sections were placed on glass slides for histological staining to assess ceramide deposition, calcification, fibrosis, and immune cell infiltration.

For visualising ceramide accumulation, cryosections of murine aortic valves were permeabilised in 0.3% Triton X-100 in PBS for 30 minutes, followed by three washes in PBS. Samples were then incubated in 5% donkey normal serum in PBS for 1 hour to block non-specific binding (Southern Biotech, #0030-01). After removing the excess blocking solution, the samples were incubated overnight by 4°C with the primary anti-ceramide antibody at a dilution of 1:50 (Sigma-Aldrich, #C8104-50TST). The samples were then washed three times in PBS and incubated for one hour with the secondary antibody donkey anti-mouse Alexa Fluor 568 at a dilution 1:1000 in PBS (Invitrogen, #A10037). After antibody staining, samples were rinsed in distilled water and incubated for 20 minutes with DAPI for nuclear counterstaining. After a final wash in distilled water, the samples were covered with Dako fluorescence mounting medium (#Dako, S3023).

To assess the degree of calcification, Calcein staining was used. Sections were incubated with 1% calcein solution (Sigma-Aldrich, #C0875) for 30 minutes at room temperature, followed by washing with PBS. Calcified areas exhibited strong green fluorescence under fluorescence microscopy, indicating mineral deposition. Masson's Trichrome staining was performed to evaluate fibrosis and collagen deposition. Sections were rehydrated, and stained according to the standard protocol using Bouin's fixative, Weigert's iron hematoxylin, Biebrich scarlet-acid fuchsin, phosphotungstic/phosphomolybdic acid, and aniline blue. Fibrotic tissue and collagen appeared blue, while muscle fibers and cytoplasm stained red. Monocyte and macrophage infiltration into the valve tissue was assessed using MOMA-2 immunohistochemistry. Following antigen retrieval, sections were incubated with MOMA-2 primary antibody (Bio-Rad, #MCA519GA) overnight at 4°C. After washing with PBS, sections were incubated with a HRP-conjugated secondary antibody, followed by detection using DAB (3,3′-

diaminobenzidine) substrate. Sections were counterstained with hematoxylin, and macrophages and monocytes were visualized as brown-stained cells.

All stained sections were imaged using a BZ-X810 microscope (Keyence), which provided high-resolution brightfield and fluorescence imaging.

#### Human Aortic Valves and Plasma Samples

Following informed consent and ethics committee approval (Reference Number EA4/028/12, Ethics Committee at the Virchow-Klinikum Campus of Charité, Berlin), human aortic valve specimens were obtained during surgical valve replacement procedures for severe aortic valve stenosis due to CAVD, in accordance with the principles of the Declaration of Helsinki. The valves were harvested with a cold ischemia time of less than 5 minutes to ensure minimal tissue degradation. After excision, the valve tissues were snap-frozen in liquid nitrogen. Control group valves were obtained from patients without aortic valve calcification, as confirmed by echocardiography and computed tomography (CT). These patients underwent valve replacement due to aortic aneurysm, where valve-sparing surgery was not possible, or due to aortic regurgitation. For omics analyses, the frozen valve tissues were pulverized using a Cellcrusher Mini device. The processed samples were stored at -80°C until further lipidomics and proteomics analyses. Fasting plasma samples of the surgical valve replacement patients were collected in EDTA tubes and stored at -80°C after centrifugation at 10,000g for 10 minutes. The validation cohort of patients with CAVD, undergoing transcatheter aortic valve replacement, provided written informed consent prior to participation. Sample collection and analyses were approved by the Ethics Committee of the Christian-Albrechts-University Kiel (file number D 506/22; AZ F 01/24) and performed in compliance with the Declaration of Helsinki. Plasma samples were collected after overnight fasting on the morning preceding transcatheter aortic valve replacement at the University Medical Centre Schleswig-Holstein – Campus Kiel. Corresponding aortic valve calcification data were obtained from pre-procedural computed tomography scans and quantitatively analyzed using the software "3mensio Structural Heart" (3mensio Medical Imaging BV, Bilthoven, Netherlands). Plasma samples, drawn into EDTA tubes, were centrifuged at 10,000 g for 10 minutes, aliquoted, and stored at -80 °C until lipidomics analyses were conducted.

#### Lipidomics Analyses

All lipidomics experiments used a Dionex Ultimate 3000 RS LC-system coupled to an Orbitrap mass spectrometer (QExactive, ThermoFisher Scientific, Bremen, Germany) equipped with a heated-electrospray ionization (HESI-II) probe. All solvents were of LC-MS grade quality and were purchased from Merck (Darmstadt, Germany). Lipid profiling was performed as described previously [13]. Briefly, lipids were extracted by adding 1 ml of methanol/methyl tertiary-butyl ether/chloroform (1.33:1:1, *v/v*) containing butylated hydroxytoluene (100 mg/L, Sigma-Aldrich, Schnelldorf, Germany) and 2.5 ml/L of SPLASH internal standard mix (Avanti Polar Lipids, Alabaster, AL, USA), to 50 µl of serum or tissue homogenate (10 – 60 mg) in 50 µl water. After incubation and centrifugation, supernatants were dried under vacuum and resuspended in 50 µl of methanol/isopropyl alcohol (1:1, *v/v*) for LC-MS/MS analysis. Extracted lipids were separated on an Accucore C30 RP column (150 × 2.1 mm, 2.6 µm) using acetonitrile/water 6:4 (*v/v*) as eluent A and isopropyl alcohol/acetonitrile (9:1, *v/v*) as eluent B; 10 mM ammonium acetate (Sigma-Aldrich, Schnelldorf, Germany) and 0.1 % formic acid (Biosolve, Dieuze, France) was added to both eluents. Ionization and data acquisition with data-dependent $MS^2$ scans (top 15) were performed as described previously [14]. Lipid species were identified according to exact mass, isotopic pattern, retention time, and fragmentation ions using Compound Discoverer 3.3 (Thermo Fisher Scientific) and two in silico databases [15]. The area under the peak was normalized to the internal standard. For tissue, samples were normalized additionally to the weight. An extraction blank and quality control samples were used to ensure data quality and linearity.

#### Proteomic Analyses

The global label-free proteome was analyzed by liquid chromatography-coupled mass spectrometry. Proteins were extracted using the EasyPep Mini MS Sample Prep Kit (Thermo Fisher) and prepared (100 µg each) via the FASP-Protocol. Proteins were processed into dried pellets and reconstituted in 0.1% formic acid in water for mass spectrometric analysis. Extracted peptides were loaded onto C18 EvoTip disposable trap columns. Chromatographic separation of 200 ng per sample was performed on an EvoSep One (EvoSep, Odense, Denmark) using a C18 Performance column (EV1137, 15cm x 150 µm, 1.5 µm) in combination with the extended 15 SPD method (88 min gradient, 220 nl/min). Mobile phases A and B were water with 0.1% (v/v) formic acid and 0.1% (v/v) acetonitrile, respectively. The liquid chromatography system was coupled to a hybrid TIMS quadrupole TOF mass spectrometer (Bruker timsTOF Flex HT) via a CaptiveSpray nano-electrospray ion source. The mass spectrometer was operated in the diaPASEF mode [16]. The MS settings were set to a scan range of 100-1700 m/z in positive ion polarity. The TIMS settings were set to an ion mobility range of 0.6-1.6 1/K0, a ramp time of 100 ms, an accumulation time of 100 ms, and a ramp rate of 9.43 Hz. The MS/MS settings were set to a mass range of 400 to 1201.0 Da and a mobility range of 0.6-1.6 1/K0. We use a fixed dia-PASEF window width of 26.0 Da, with 1.0 Da mass overlap and 0.0 1/K0 mobility overlap. The method was set to 32 mass steps and 1.0 mobility window per cycle. The entire method had an estimated cycle time of 1.80 seconds.

##### *diaPASEF data processing*

The acquired diaPASEF mass spectra were processed using the DIA-NN v1.8.1 software tool [17]. The software was used in the high precision LC mode, with RT-dependent cross normalization enabled, smart profiling for Library generation, and single pass mode as neural network classifier. Mass accuracy and MS1 accuracy were set to 10.0, scan window was set to 0 (DIA-NN automatically determines scan window width). The 'Match between runs' option was enabled. The human UniProtKB/swiss-prot database (downloaded on 12/22/2021) was used for deep learning based on *in silico* spectral library generation. The protease setting was set to trypsin/P, while 1.0 missed cleavage was accepted. The maximum number of variable modifications was set to 0.0, N-terminal methionine excision and cysteine carbamidomethylation enabled as fixed modifications. The peptide length range was set to 7-30, precursor charge range to 2-4, precursor m/z range to 100-1700, and fragment ion m/z range to 400-1201. The 'pg_matrix' (protein groups) output, was used for downstream analysis.

For further data processing, Perseus software platform by Max-Planck-Institute of Biochemistry was used to ensure reliable and interpretable results [18]. The pre-processing workflow started with the explicit group annotation. After filtering for valid values, $\log_2$ transformation and imputation addressed data standardization and missing values. Data were exported as text files for further use on BigOmics analytics platform. Statistical significance for Volcano plot of proteins for Fig. 2A was assessed using three independent statistical methods: DESeq2 (Wald test) [19], edgeR (QLF test) [20] and limma-trend [21]. Test results were exported and visualized using the R package ggplot2 (v3.5.1). The maximum q-value of the three methods was taken as aggregate q-value, which corresponds to taking the intersection of significant proteins from all three tests. For the clustered heatmap shown in Fig. 2B, hierarchical clustering was performed using an unsupervised two-way approach to simultaneously cluster both proteins and samples. The heatmap displays the abundance matrix of proteins, which was scaled to an average relative expression range to highlight differences between study and control groups. The clustering was based on protein expression levels across the samples, using a top 30 protein selection ranked by standard deviation (Top N = 30). Proteins were grouped into two modules (K modules = 2), and the color scale represents relative expression levels, ranging from blue (low expression) to red (high expression). Functional annotation of protein modules was performed to identify enriched biological processes and pathways for the clusters, with correlations (R values) visualized in separate scatter plots for modules S1 and S2. Annotation scores were calculated, highlighting processes such as adipogenesis and immune response, and the clustering results were summarized based on phenotypic grouping. Weighted correlation network analysis (WGCNA) for Fig. 2C-G was performed using the respective BigOmics module [22]. Briefly, modules were identified as branches in a hierarchical clustering tree, with dynamic branch cutting applied to optimize module assignment (Supplementary Fig. 2B). The eigengene of each module, representing the first principal component of the standardized protein abundance profiles, was calculated to summarize the module's characteristics. Relationships between module eigengenes and phenotypic traits were assessed using Pearson correlation coefficients, which range from -1 (fully negative correlation) to 1 (fully positive correlation) (Fig. 2C). To investigate key interactions within modules, partial correlation networks were generated, by computing regularized partial correlation matrix is using the 'graphical lasso' (Glasso) with BIC model selection, displaying connections among the most correlated features, with edges colored based on the correlation direction (positive or negative) (Fig. 2G). Functional enrichment analysis of identified modules was performed using Fisher's exact test to determine overrepresented biological pathways and functions, providing insights into the biological relevance of the modules (Fig. 2D-F).

#### Correlation Analysis of Omics Data

Partial correlation analysis between proteins and lipids from tissue measurements was performed in R (v4.4.1) with PPCOR (v1.1) [23], correcting for sex and age effects using Spearman's rank correlation. 53 target proteins and 93 target lipids were analyzed (Supplementary Table 6). Adjusted p-values were calculated by applying Benjamini-Hochberg correction, and all correlations with values below 0.05 were considered for Fig. 2H. The analysis was restricted to patients where both tissue lipid and protein measurements were present ($n_{total} = 19$; $n_{CAVDpatients} = 9$, $n_{controls} = 10$). For partial correlation against selected human

MSigDB hallmark gene sets [9], a gene set variation analysis with Gaussian kernel was performed with GSVA (v1.52.3) on the protein dataset. The partial correlation against the lipids was done, as explained above. All correlations with $p < 0.05$ were plotted in Fig. 2I for an explorative overview. Effect sizes were calculated as Cohen's *d* using EFFECT SIZE (v0.8.9) [9] for all target lipids and target proteins; values below 0.2 were considered as a very small effect size, between 0.2 and 0.5 a small effect size, between 0.5 and 0.8 as medium effect size and values above 0.8 were considered as large effect sizes, respectively. The calculated effect sizes of the lipids and proteins are represented in the correlation Fig. by the color of the molecule names.

#### Culture of Primary Human Valvular Interstitial Cells

Primary human valvular interstitial cells (hVICs) were purchased from Innoprot (Innoprot, ref: P10977). Cells were cultured in Valvular Interstitial Cell Medium (Innoprot, ref: P60177), supplemented with 5% fetal bovine serum (FBS) and 1% penicillin-streptomycin (Thermo Fisher Scientific). Cells were maintained at 37°C in a humidified atmosphere containing 5% $CO_2$. The culture medium was refreshed every 2-3 days, and cells were passaged upon reaching 80-90% confluence. All experiments were conducted with cells between passages 2 and 5.

For calcification assays, hVICs were seeded in 6-well plates at a density of $1x10^5$ cells per well. After reaching confluence, cells were treated with a calcification medium consisting of 10 mM β-glycerophosphate, 50 µg/mL ascorbic acid, and 100 nM dexamethasone, supplemented with either C2 ceramide (Sigma-Aldrich, dissolved in DMSO) or DMSO alone as a control with the same interval of media refreshment for 21 days to induce calcification.

#### Calcium assay and Alizarin Red Staining

To assess calcification in hVICs, both a calcium assay and Alizarin Red staining were performed. Following the treatment period with calcification medium supplemented with C2 ceramide, C18 ceramide, or DMSO, hVICs were washed twice with phosphate-buffered saline

(PBS) and fixed with 4% paraformaldehyde for 15 minutes at room temperature. For Alizarin Red staining, fixed cells were incubated with 2% Alizarin Red S solution (pH 4.2, Sigma-Aldrich) for 20 minutes at room temperature with gentle agitation. Excess dye was removed by washing the cells thoroughly with distilled water. Calcified nodules stained bright red, indicating calcium deposition. The stained wells were imaged under a light microscope (Keyence BZ-X810), and images were captured for qualitative analysis. For quantification, the Alizarin Red stain was solubilized with 10% acetic acid for 30 minutes, followed by neutralization with 10% ammonium hydroxide. Absorbance was measured at 405 nm using a microplate reader, and the intensity of the signal was used to quantify the amount of calcium deposited in the cells.

The calcium assay was conducted to measure the calcium content in the cells. Cells were decalcified by incubating with 0.6 N HCl for 24 hours at 4°C. The supernatant was collected, and the calcium content was quantified using a Calcium Colorimetric Assay Kit (Sigma-Aldrich, MAK022) according to the manufacturer's instructions. Absorbance was measured at 575 nm, and calcium levels were normalized to total protein content, which was determined using the BCA protein assay.

#### RNA isolation and quantitative real-time PCR

hVICs were stimulated for 4 h with short-chain C2 ceramide (C2-Cer) or long-chain C18 ceramide (C18-Cer) at concentrations of 0.1, 1, 5, and 10 µM (n = 4 independent experiments per condition), alongside DMSO (negative control, same final concentration as in other experimental conditions) and 1 µg/mL tunicamycin (positive control). Cells were lysed in RLT buffer (Qiagen) supplemented with 1% β-mercaptoethanol, and total RNA was extracted using the RNeasy® Mini Kit (Qiagen) and reverse-transcribed with the High-Capacity cDNA Reverse Transcription Kit (Applied Biosystems™). Quantitative PCR was performed using 5 ng cDNA with TaqMan Gene Expression Assays on an Applied Biosystems™ QuantStudio 7

Flex system. Reference gene stability (GUSB, HPRT, RPL37A) was evaluated using RefFinder, identifying GUSB (Hs00939627_m1) as the most stable gene for normalization. Relative expression of ATF4 (Hs00909569_g1), DDIT3 (Hs99999172_m1), and HSPA5 (Hs99999174_m1) was calculated using the $2^{-\Delta\Delta Ct}$ method.

#### Western blotting

hVICs were stimulated for 16 hours with DMSO as negative control, tunicamycin (1 µg/mL) as positive control, or C2- and C18-ceramide (10 µM each) prior to protein extraction. Cells were lysed in ice-cold RIPA buffer (Abcam, ab156034) supplemented with protease and phosphatase inhibitors (Fisher scientific, PI78440). Lysates were clarified by centrifugation at 10,000 rpm for 20 minutes at 4°C, and protein concentrations were determined using a colorimetric protein assay with bovine serum albumin standards (Fisher scientific, PI23225). Equal amounts of protein, typically 10 µg per lane, were denatured under reducing conditions of 500mM DTT (30x Biolabs B7705S) and separated on 4–12% Bis-Tris SDS-polyacrylamide gels (ThermoFisher, NP0322BOX) using MES running buffer (NuPage 20x NP0002). Proteins were transferred onto PVDF membranes (Amersham IPVH 0001), which were subsequently blocked in 5% non-fat dry milk (Roth T145.2) in Tris-buffered saline containing 0.1% Tween-20. Membranes were incubated with primary antibody for GRP78/BiP (Abcam, ab21685) at a dilution of 1:5000 overnight at 4°C, followed by horseradish peroxidase–conjugated secondary antibody (Cell signaling, 7076 S) at a dilution of 1:5000 for 60 minutes at room temperature. Immunoreactive bands were visualized using enhanced chemiluminescence (Amersham, RPN 2232) and imaged with ChemiDoc imaging system Azure 400 (Azure Biosystems). Densitometric quantification was performed using ImageJ software, and target protein expression was normalized to β-actin (Merck, A1978).

#### Enzyme-linked Immunosorbent Assays

For the detection of IL-6 in the cell culture supernatant and LDL-cholesterol as well as PCSK9 in mouse blood, enzyme-linked immunosorbent assays (ELISA) were performed according to the manufacturer's protocols. The IL-6 ELISA (ThermoFisher, KHC0061) was carried out on supernatants collected from hVIC cultures following 24-hour treatment with calcification medium supplemented with C2 ceramide (10 µM) or DMSO. Supernatants were centrifuged at 300 x g for 10 minutes at 4°C to remove cellular debris before analysis. PCSK9 levels (Abcam, ab215538) were measured in plasma samples collected from the mice to verify successful AAV injection in the murine model. LDL-cholesterol levels were analyzed in murine plasma samples (Novus Biologicals, NBP2-81135). Blood was collected via retro-orbital venous plexus, and plasma was separated by centrifugation at 10,000 x g for 10 minutes.

#### Proteome Profiler

To evaluate the activation of the NF-κB signaling pathway in hVICs following C2 ceramide treatment, we utilized the Proteome Profiler Human NF-κB Pathway Array (R&D Systems, #ARY029). Cell lysates were prepared from hVICs treated with C2 ceramide (10 µM, Sigma-Aldrich) for 24 hours and from control cells treated with DMSO. After treatment, cells were washed with ice-cold PBS and scraped into RIPA buffer (Thermo Fisher Scientific) containing HALT protease and phosphatase inhibitor cocktail (Thermo Fisher Scientific) using cell scrapers. The cell suspensions were incubated on ice for 60 minutes to allow for efficient lysis. The lysates were then centrifuged at 14,000 x g for 10 minutes at 4°C to remove cellular debris, and the supernatants containing the proteins were collected. Protein concentrations were measured using a BCA protein assay. For the proteome analysis, 300 µg of total protein from each sample was incubated with the Proteome Profiler membrane overnight at 4°C. The membrane, containing antibodies specific for 45 proteins involved in the NF-κB signaling pathway, was washed and incubated with a cocktail of biotinylated detection antibodies. After washing, the membrane was treated with streptavidin-HRP, and the signal was visualized using chemiluminescence. Spot intensity was quantified using Quick Spots image analysis software, by R&D Systems.

#### Data availability

The lipidomics data are available at the NIH Common Fund's National Metabolomics Data Repository (NMDR) (Study ID ST003682 and ST003684) [24]. For peer review, the data are temporarily available at https://dev.metabolomicsworkbench.org:22222/data/DRCCMetadata.php?Mode=Study&StudyID=ST003682&Access=MgzJ8575 and https://dev.metabolomicsworkbench.org:22222/data/DRCCMetadata.php?Mode=Study&StudyID=ST003684&Access=CkpJ9804.

The proteomics data have been deposited to the ProteomeXchange Consortium via the PRIDE [25] partner repository with the dataset identifier PXD058987. For peer review please log in to the PRIDE website (https://www.ebi.ac.uk/pride/login) using the following details: Project accession: PXD058987 and Token: hMm2l8uiSeJR. Single-cell transcriptomic are available at the GEO repository (GEO accession GSE294099, https://www.ncbi.nlm.nih.gov/geo/query/acc.cgi?acc=GSE294099) using the following reviewer token: gpmtacqchhefbin.

#### Statistical analysis

All Omics Data were analyzed as indicated above. Other data were analyzed using GraphPad Prism 10 (GraphPad Software) or SPSS 25 (IBM). Sample sizes for animal experiments were planned based on a power analysis by G*Power using the transvalvular mean aortic valve gradient as primary outcome parameter. Significance was considered when $p < 0.05$. Depending on the dataset and experimental design, different statistical methods were used as indicated in the Figure legends. Parametric statistics (e.g., t-Test, ANOVA) were only applied if assumptions were met, i.e., datasets were examined for Gaussian distribution by D'Agostino-Pearson test, aided by visual inspection of the data, and homogeneity of variances by Brown-Forsythe, Levene's or F-test (depending on the statistical method used). If assumptions for parametric procedures were not met or could not be reliably assumed due to small sample size, non-parametric methods were used as indicated. Two-tailed tests were applied if not indicated otherwise. For multiple comparisons following ANOVA, Bonferroni-adjusted post hoc tests were applied.

**Supplementary Table 1:** Lipidome of human aortic valves – Fold Change CAVD patients vs. Controls

| Lipid | Fold Change (FC) | log2(FC) | raw.pval | p.adjusted | -LOG10(p) |
|---|---|---|---|---|---|
| CAR 10:0 | 2.1651 | 1.1144 | 0.00076461 | 0.010791 | 1.9669 |
| CAR 12:0 | 1.6405 | 0.71413 | 0.02025 | 0.060588 | 1.2176 |
| CAR 12:1 | 2.4121 | 1.2703 | 0.00022148 | 0.0075304 | 2.1232 |
| CAR 13:0 | 2.3534 | 1.2348 | 0.05109 | 0.10857 | 0.96431 |
| CAR 13:1 | 1.9655 | 0.97488 | 0.0016 | 0.015748 | 1.8028 |
| CAR 14:0 | 1.089 | 0.12297 | 0.23401 | 0.33468 | 0.47537 |
| CAR 14:1 | 1.7011 | 0.76647 | 0.017844 | 0.056316 | 1.2494 |
| CAR 14:2 | 1.9564 | 0.96823 | 0.0019061 | 0.016202 | 1.7904 |
| CAR 15:0 | 0.96389 | -0.053052 | 0.45087 | 0.55378 | 0.25666 |
| CAR 16:0 | 1.276 | 0.35168 | 0.056773 | 0.11635 | 0.93425 |
| CAR 16:1 | 1.1337 | 0.18109 | 0.28978 | 0.38984 | 0.40911 |
| CAR 16:2 | 1.4943 | 0.57945 | 0.036742 | 0.085617 | 1.0674 |
| CAR 16:3 | 2.4757 | 1.3078 | 0.0026755 | 0.01888 | 1.724 |
| CAR 17:0 | 1.0545 | 0.076511 | 0.81318 | 0.8519 | 0.06961 |
| CAR 17:1 | 0.91099 | -0.13449 | 0.40049 | 0.50348 | 0.29802 |
| CAR 18:0 | 1.5737 | 0.65417 | 0.041102 | 0.093733 | 1.0281 |
| CAR 18:1 | 0.87038 | -0.20028 | 0.71483 | 0.76494 | 0.11637 |
| CAR 18:2 | 1.0143 | 0.02046 | 0.71483 | 0.76494 | 0.11637 |
| CAR 18:3 | 1.4109 | 0.49666 | 0.093216 | 0.16924 | 0.77151 |
| CAR 19:0 | 1.4303 | 0.51635 | 0.14562 | 0.23526 | 0.62846 |
| CAR 19:1 | 1.1222 | 0.16639 | 0.53255 | 0.61759 | 0.2093 |
| CAR 20:0 | 1.5418 | 0.62464 | 0.036742 | 0.085617 | 1.0674 |
| CAR 20:1 | 1.0075 | 0.010818 | 0.53255 | 0.61759 | 0.2093 |
| CAR 20:2 | 1.1684 | 0.22458 | 0.21718 | 0.31666 | 0.4994 |
| CAR 20:3 | 1.3414 | 0.42376 | 0.056773 | 0.11635 | 0.93425 |
| CAR 20:4 | 1.7102 | 0.77421 | 0.0050625 | 0.02744 | 1.5616 |

| | | | | | |
|---|---|---|---|---|---|
| CAR 20:5 | 1.4528 | 0.53888 | 0.025885 | 0.068659 | 1.1633 |
| CAR 22:0 | 1.7712 | 0.82472 | 0.0079031 | 0.035398 | 1.451 |
| CAR 22:1 | 1.2791 | 0.3551 | 0.21718 | 0.31666 | 0.4994 |
| CAR 22:4 | 1.2005 | 0.26366 | 0.2012 | 0.3004 | 0.5223 |
| CAR 24:0 | 2.5655 | 1.3593 | 5.03E-06 | 0.0018806 | 2.7257 |
| CAR 24:1 | 2.1533 | 1.1066 | 8.73E-05 | 0.0046651 | 2.3311 |
| CAR 24:2 | 1.6848 | 0.75259 | 0.041102 | 0.093733 | 1.0281 |
| CAR 26:0 | 2.2667 | 1.1806 | 0.0016 | 0.015748 | 1.8028 |
| CAR 26:1 | 2.7566 | 1.4629 | 0.00042051 | 0.0085011 | 2.0705 |
| CAR 26:2 | 2.5849 | 1.3701 | 0.022923 | 0.063978 | 1.194 |
| CAR 8:0 | 2.058 | 1.0412 | 0.0031534 | 0.02016 | 1.6955 |
| CE 16:1 | 4.8258 | 2.2708 | 0.0019061 | 0.016202 | 1.7904 |
| CE 17:0 | 1.322 | 0.40275 | 0.069659 | 0.13712 | 0.86291 |
| CE 17:1 | 3.7753 | 1.9166 | 0.012021 | 0.044075 | 1.3558 |
| CE 20:1 | 1.8715 | 0.9042 | 0.045875 | 0.10004 | 0.99982 |
| CE 20:2 | 1.8077 | 0.85416 | 0.036742 | 0.085617 | 1.0674 |
| CE 20:4 | 6.2448 | 2.6427 | 3.10E-05 | 0.0043427 | 2.3622 |
| CE 20:5 | 13.023 | 3.703 | 1.31E-05 | 0.0032662 | 2.486 |
| CE 22:2 | 1.5635 | 0.64473 | 0.25171 | 0.35325 | 0.45192 |
| CE 22:3 | 1.1684 | 0.22457 | 0.47731 | 0.57771 | 0.23829 |
| CE 22:4 | 1.0858 | 0.11882 | 0.25171 | 0.35325 | 0.45192 |
| CE 22:5 | 5.095 | 2.3491 | 0.00051551 | 0.0086482 | 2.0631 |
| CE 22:6 | 10.539 | 3.3976 | 3.58E-06 | 0.0018806 | 2.7257 |
| CE 24:2 | 1.0952 | 0.13125 | 0.55569 | 0.64197 | 0.19249 |
| CE 24:3 | 1.6121 | 0.68893 | 0.45807 | 0.56078 | 0.25121 |
| CE 24:4 | 0.69425 | -0.52648 | 0.65162 | 0.71259 | 0.14716 |
| CE 24:5 | 0.50726 | -0.9792 | 0.62087 | 0.68699 | 0.16305 |
| CE 26:2 | 1.6703 | 0.74014 | 0.59074 | 0.66648 | 0.17621 |
| CE 26:4 | 1.443 | 0.52906 | 0.94862 | 0.95758 | 0.018823 |
| CE 28:5 | 0.69262 | -0.52987 | 0.71483 | 0.76494 | 0.11637 |
| Cer 16:0;O2/24:1 | 2.4219 | 1.2761 | 0.0031534 | 0.02016 | 1.6955 |
| Cer 16:0;O3/24:1 | 2.0262 | 1.0188 | 0.056773 | 0.11635 | 0.93425 |
| Cer 16:1;O2/23:0 | 2.1644 | 1.114 | 0.012021 | 0.044075 | 1.3558 |
| Cer 16:1;O2/24:0;(2OH) | 2.6265 | 1.3931 | 0.0022621 | 0.017444 | 1.7584 |
| Cer 16:1;O2/24:1 | 1.6304 | 0.70524 | 0.015684 | 0.052844 | 1.277 |
| Cer 16:1;O2/24:2 | 1.6604 | 0.73152 | 0.1337 | 0.22224 | 0.65318 |
| Cer 17:1;O2/16:0 | 4.0154 | 2.0055 | 0.0022621 | 0.017444 | 1.7584 |
| Cer 17:1;O2/24:1 | 1.7432 | 0.80178 | 0.076919 | 0.14787 | 0.83012 |
| Cer 18:0;O2/20:0 | 1.8884 | 0.91719 | 0.036742 | 0.085617 | 1.0674 |
| Cer 18:0;O2/22:0 | 2.2666 | 1.1806 | 0.012021 | 0.044075 | 1.3558 |
| Cer 18:0;O2/23:0 | 2.8465 | 1.5092 | 0.0079031 | 0.035398 | 1.451 |
| Cer 18:0;O2/24:0 | 1.9682 | 0.97687 | 0.0037044 | 0.022712 | 1.6437 |
| Cer 18:0;O2/24:2 | 1.8367 | 0.87708 | 0.18608 | 0.28464 | 0.5457 |
| Cer 18:0;O3/16:0 | 2.9748 | 1.5728 | 0.0043372 | 0.025346 | 1.5961 |

| | | | | | |
|---|---|---|---|---|---|
| Cer 18:0;O3/22:0 | 2.9149 | 1.5434 | 0.025885 | 0.068659 | 1.1633 |
| Cer 18:0;O3/23:0 | 2.265 | 1.1795 | 0.0079031 | 0.035398 | 1.451 |
| Cer 18:0;O3/24:0 | 2.2811 | 1.1897 | 0.0031534 | 0.02016 | 1.6955 |
| Cer 18:0;O3/24:1 | 2.962 | 1.5666 | 0.010482 | 0.041049 | 1.3867 |
| Cer 18:1;O2/16:0;(2OH) | 1.3085 | 0.38794 | 0.18608 | 0.28464 | 0.5457 |
| Cer 18:1;O2/18:0 | 0.87452 | -0.19344 | 0.50454 | 0.59904 | 0.22254 |
| Cer 18:1;O2/20:0 | 1.3891 | 0.47418 | 0.18608 | 0.28464 | 0.5457 |
| Cer 18:1;O2/22:0 | 1.2929 | 0.37062 | 0.12252 | 0.20877 | 0.68034 |
| Cer 18:1;O2/23:0 | 2.9012 | 1.5367 | 0.0050625 | 0.02744 | 1.5616 |
| Cer 18:1;O2/24:0;(2OH) | 4.4671 | 2.1593 | 0.0043372 | 0.025346 | 1.5961 |
| Cer 18:1;O2/24:1 | 1.6492 | 0.72177 | 0.21718 | 0.31666 | 0.4994 |
| Cer 18:1;O2/24:1;(2OH) | 3.5898 | 1.8439 | 0.0019061 | 0.016202 | 1.7904 |
| Cer 18:1;O2/26:0 | 1.5636 | 0.64485 | 0.18608 | 0.28464 | 0.5457 |
| Cer 18:1;O2/26:1 | 1.3863 | 0.47127 | 0.37659 | 0.47988 | 0.31887 |
| Cer 18:1;O3/22:0 | 2.8027 | 1.4868 | 0.0094011 | 0.039955 | 1.3984 |
| Cer 18:1;O3/23:0 | 3.6523 | 1.8688 | 0.0019061 | 0.016202 | 1.7904 |
| Cer 18:1;O3/24:0 | 2.5335 | 1.3411 | 0.0058902 | 0.029972 | 1.5233 |
| Cer 18:1;O3/24:1 | 3.1712 | 1.665 | 0.0050625 | 0.02744 | 1.5616 |
| Cer 18:2;O2/16:0 | 2.6636 | 1.4134 | 0.0050625 | 0.02744 | 1.5616 |
| Cer 18:2;O2/18:0 | 1.1983 | 0.26096 | 0.23401 | 0.33468 | 0.47537 |
| Cer 18:2;O2/20:0 | 1.4661 | 0.55198 | 0.11207 | 0.1945 | 0.71109 |
| Cer 18:2;O2/22:0 | 1.4576 | 0.54359 | 0.14562 | 0.23526 | 0.62846 |
| Cer 18:2;O2/23:0 | 2.161 | 1.1117 | 0.0079031 | 0.035398 | 1.451 |
| Cer 18:2;O2/24:0 | 1.9413 | 0.95701 | 0.10231 | 0.18134 | 0.7415 |
| Cer 18:2;O2/24:2 | 1.4438 | 0.52988 | 0.65162 | 0.71259 | 0.14716 |
| Cer 18:2;O2/26:0 | 2.2437 | 1.1659 | 0.062953 | 0.12624 | 0.89879 |
| Cer 19:1;O2/22:0 | 1.6898 | 0.75684 | 0.05109 | 0.10857 | 0.96431 |
| Cer 19:1;O2/24:0 | 1.599 | 0.6772 | 0.11207 | 0.1945 | 0.71109 |
| Cer 37:1;O2 | 1.4521 | 0.53818 | 0.37659 | 0.47988 | 0.31887 |
| Cer 42:3;O2 | 1.2209 | 0.2879 | 0.40049 | 0.50348 | 0.29802 |
| Cer 44:3;O2 | 0.96498 | -0.051428 | 0.84669 | 0.8784 | 0.056307 |
| CL 70:6 | 0.52675 | -0.9248 | 0.093216 | 0.16924 | 0.77151 |
| CL 70:7 | 0.44772 | -1.1593 | 0.05109 | 0.10857 | 0.96431 |
| CL 72:5 | 0.6372 | -0.65018 | 0.11207 | 0.1945 | 0.71109 |
| CL 74:10 | 0.31876 | -1.6494 | 0.017844 | 0.056316 | 1.2494 |
| CL 74:6 | 0.73959 | -0.4352 | 0.093216 | 0.16924 | 0.77151 |
| CL 74:9 | 0.41942 | -1.2535 | 0.022923 | 0.063978 | 1.194 |
| CoQ10 | 1.3478 | 0.4306 | 0.53255 | 0.61759 | 0.2093 |
| CoQ9 | 2.8867 | 1.5294 | 0.045875 | 0.10004 | 0.99982 |
| DG 16:0_16:0 | 0.67048 | -0.57674 | 0.50454 | 0.59904 | 0.22254 |
| DG 16:0_18:1 | 0.8159 | -0.29353 | 0.81318 | 0.8519 | 0.06961 |
| DG 16:0_18:2 | 0.85886 | -0.21951 | 0.65162 | 0.71259 | 0.14716 |
| DG 16:0_20:4 | 0.58523 | -0.77293 | 0.1337 | 0.22224 | 0.65318 |
| DG 16:0_22:6 | 0.92012 | -0.12011 | 0.2703 | 0.37166 | 0.42985 |

| | | | | | |
|---|---|---|---|---|---|
| DG 16:0_24:7 | 0.35025 | -1.5135 | 0.05109 | 0.10857 | 0.96431 |
| DG 16:1_18:1 | 0.57706 | -0.7932 | 0.71483 | 0.76494 | 0.11637 |
| DG 18:0_18:1 | 0.65518 | -0.61004 | 0.35355 | 0.45675 | 0.34032 |
| DG 18:0_20:4 | 0.54233 | -0.88276 | 0.076919 | 0.14787 | 0.83012 |
| DG 18:0_22:4 | 0.66462 | -0.58941 | 0.2012 | 0.3004 | 0.5223 |
| DG 18:0_22:6 | 0.38334 | -1.3833 | 0.1337 | 0.22224 | 0.65318 |
| DG 18:0_24:7 | 0.17086 | -2.5491 | 0.00042051 | 0.0085011 | 2.0705 |
| DG 18:1_18:1 | 0.52716 | -0.9237 | 0.81318 | 0.8519 | 0.06961 |
| DG 18:1_18:2 | 0.91325 | -0.13092 | 1 | 1 | 0 |
| DG 18:1_20:4 | 1.2355 | 0.30506 | 0.53255 | 0.61759 | 0.2093 |
| DG 18:1_22:5 | 0.99222 | -0.011266 | 0.65162 | 0.71259 | 0.14716 |
| DG 18:1_22:6 | 1.5184 | 0.60254 | 0.65162 | 0.71259 | 0.14716 |
| DG 32:1 | 0.74058 | -0.43328 | 0.50454 | 0.59904 | 0.22254 |
| DG 32:3 | 0.6117 | -0.70911 | 0.31014 | 0.41206 | 0.38504 |
| FA 18:1 | 1.3605 | 0.44418 | 0.11207 | 0.1945 | 0.71109 |
| FA 18:2 | 1.4202 | 0.50608 | 0.084761 | 0.1585 | 0.79996 |
| FA 20:4 | 1.3436 | 0.42615 | 0.14562 | 0.23526 | 0.62846 |
| FA 22:6 | 2.2888 | 1.1946 | 0.0011149 | 0.012636 | 1.8984 |
| GM3 33:1;O2 | 2.7218 | 1.4445 | 0.0058902 | 0.029972 | 1.5233 |
| GM3 41:2;O2 | 2.2826 | 1.1907 | 0.0043372 | 0.025346 | 1.5961 |
| GM3 44:1;O2 | 1.3444 | 0.42696 | 0.56129 | 0.64197 | 0.19249 |
| Hex2Cer 18:1;O2/24:0 | 1.0394 | 0.055708 | 0.94862 | 0.95758 | 0.018823 |
| Hex2Cer 18:1;O2/24:1 | 1.4502 | 0.53629 | 0.28978 | 0.38984 | 0.40911 |
| Hex2Cer 18:2;O2/16:0 | 2.6715 | 1.4176 | 0.0031534 | 0.02016 | 1.6955 |
| Hex2Cer 18:2;O2/24:0 | 1.3726 | 0.45691 | 0.2012 | 0.3004 | 0.5223 |
| Hex2Cer 18:2;O2/24:1 | 1.898 | 0.92445 | 0.17179 | 0.26826 | 0.57144 |
| Hex2Cer 32:1;O2 | 2.2513 | 1.1708 | 0.036742 | 0.085617 | 1.0674 |
| Hex3Cer 16:1;O2/24:1 | 1.4229 | 0.50879 | 0.14562 | 0.23526 | 0.62846 |
| Hex3Cer 18:1;O2/16:0 | 2.2196 | 1.1503 | 0.02025 | 0.060588 | 1.2176 |
| Hex3Cer 18:1;O2/18:0 | 0.99426 | -0.0083018 | 0.84669 | 0.8784 | 0.056307 |
| Hex3Cer 18:1;O2/20:0 | 1.0869 | 0.12016 | 0.65162 | 0.71259 | 0.14716 |
| Hex3Cer 18:1;O2/22:0 | 1.2703 | 0.34517 | 0.53255 | 0.61759 | 0.2093 |
| Hex3Cer 18:1;O2/24:0 | 1.2538 | 0.32635 | 0.50454 | 0.59904 | 0.22254 |
| Hex3Cer 18:1;O2/24:1 | 1.1744 | 0.23198 | 0.50454 | 0.59904 | 0.22254 |
| Hex3Cer 18:2;O2/24:0 | 1.205 | 0.26909 | 0.59074 | 0.66648 | 0.17621 |
| Hex3Cer 32:1;O2 | 3.2021 | 1.679 | 0.012021 | 0.044075 | 1.3558 |
| Hex3Cer 41:1;O2 | 1.4035 | 0.489 | 0.15831 | 0.25088 | 0.60053 |
| Hex3Cer 44:1;O2 | 1.4188 | 0.50466 | 0.62087 | 0.68699 | 0.16305 |
| HexCer 17:1;O2/22:0 | 2.6362 | 1.3985 | 0.022923 | 0.063978 | 1.194 |
| HexCer 18:0;O3/16:0;(2OH) | 3.7163 | 1.8939 | 0.0068331 | 0.033626 | 1.4733 |
| HexCer 18:1;O2/20:0 | 1.6341 | 0.70847 | 0.35355 | 0.45675 | 0.34032 |
| HexCer 18:1;O2/22:0 | 1.7609 | 0.81629 | 0.045875 | 0.10004 | 0.99982 |
| HexCer 18:1;O2/23:0 | 2.5976 | 1.3772 | 0.0068331 | 0.033626 | 1.4733 |
| HexCer 18:1;O2/24:0 | 2.2405 | 1.1638 | 0.022923 | 0.063978 | 1.194 |

| | | | | | |
|---|---|---|---|---|---|
| HexCer 18:1;O2/24:1 | 1.6417 | 0.71522 | 0.076919 | 0.14787 | 0.83012 |
| HexCer 18:2;O2/22:0 | 2.26 | 1.1763 | 0.10231 | 0.18134 | 0.7415 |
| HexCer 18:2;O2/24:0 | 2.1354 | 1.0945 | 0.069659 | 0.13712 | 0.86291 |
| LNAPE 16:0/N-18:0 | 1.049 | 0.069054 | 0.81318 | 0.8519 | 0.06961 |
| LNAPE 18:1/N-20:0 | 1.1551 | 0.20799 | 0.25171 | 0.35325 | 0.45192 |
| LNAPE 18:1/N-20:1 | 0.99496 | -0.0072836 | 0.94862 | 0.95758 | 0.018823 |
| LNAPE 18:2/N-16:0 | 6.0544 | 2.598 | 0.0022621 | 0.017444 | 1.7584 |
| LNAPE 18:2/N-17:0 | 1.3586 | 0.44217 | 0.59074 | 0.66648 | 0.17621 |
| LNAPE 18:2/N-20:0 | 1.4089 | 0.49452 | 0.062953 | 0.12624 | 0.89879 |
| LNAPE 18:2/N-20:1 | 1.0067 | 0.0096832 | 0.98286 | 0.9855 | 0.0063434 |
| LNAPE 20:2/N-20:0 | 0.8431 | -0.24623 | 0.81318 | 0.8519 | 0.06961 |
| LNAPE 20:3/N-18:0 | 1.6117 | 0.68862 | 0.025885 | 0.068659 | 1.1633 |
| LNAPE 20:3/N-20:0 | 1.9144 | 0.9369 | 0.0058902 | 0.029972 | 1.5233 |
| LNAPE 20:4/N-16:0 | 7.8098 | 2.9653 | 0.012021 | 0.044075 | 1.3558 |
| LNAPE 20:4/N-18:0 | 1.0849 | 0.11761 | 0.77999 | 0.82523 | 0.083426 |
| LNAPE 20:4/N-20:0 | 1.2379 | 0.30785 | 0.2012 | 0.3004 | 0.5223 |
| LNAPE 20:4/N-20:2 | 2.4861 | 1.3139 | 0.093216 | 0.16924 | 0.77151 |
| LNAPE 22:4/N-18:0 | 1.5022 | 0.58703 | 0.022923 | 0.063978 | 1.194 |
| LNAPE 22:4/N-20:0 | 1.5359 | 0.61909 | 0.17179 | 0.26826 | 0.57144 |
| LNAPE 32:0 | 1.0123 | 0.017663 | 0.91448 | 0.93446 | 0.029437 |
| LNAPE 34:1 | 1.0278 | 0.039513 | 0.84669 | 0.8784 | 0.056307 |
| LNAPE 36:0 | 1.2397 | 0.31001 | 0.17179 | 0.26826 | 0.57144 |
| LNAPE 36:1 | 1.0846 | 0.11716 | 0.62087 | 0.68699 | 0.16305 |
| LNAPE 36:2 | 1.3156 | 0.39574 | 0.062953 | 0.12624 | 0.89879 |
| LNAPE 37:4 | 0.94093 | -0.087843 | 0.98286 | 0.9855 | 0.0063434 |
| LNAPE 40:6 | 1.732 | 0.79242 | 0.31014 | 0.41206 | 0.38504 |
| LNAPE 42:5 | 1.7761 | 0.82873 | 0.02025 | 0.060588 | 1.2176 |
| LPC 14:0 | 2.5247 | 1.3361 | 0.00051551 | 0.0086482 | 2.0631 |
| LPC 15:0 | 1.8798 | 0.91058 | 0.0031534 | 0.02016 | 1.6955 |
| LPC 16:0 | 2.3329 | 1.2221 | 0.00034126 | 0.0085011 | 2.0705 |
| LPC 16:1 | 2.2793 | 1.1886 | 0.0016 | 0.015748 | 1.8028 |
| LPC 17:0 | 3.6214 | 1.8565 | 0.0058902 | 0.029972 | 1.5233 |
| LPC 17:1 | 2.6126 | 1.3855 | 0.00051551 | 0.0086482 | 2.0631 |
| LPC 18:0 | 2.8213 | 1.4964 | 0.00027569 | 0.0076376 | 2.117 |
| LPC 18:1 | 2.968 | 1.5695 | 0.00022148 | 0.0075304 | 2.1232 |
| LPC 18:2 | 2.2116 | 1.1451 | 0.0026755 | 0.01888 | 1.724 |
| LPC 18:3 | 2.5041 | 1.3243 | 0.0011149 | 0.012636 | 1.8984 |
| LPC 19:0 | 3.5257 | 1.8179 | 0.00034126 | 0.0085011 | 2.0705 |
| LPC 19:1 | 3.5137 | 1.813 | 0.00022148 | 0.0075304 | 2.1232 |
| LPC 20:0 | 3.9315 | 1.9751 | 8.73E-05 | 0.0046651 | 2.3311 |
| LPC 20:1 | 3.1147 | 1.6391 | 6.82E-05 | 0.0046651 | 2.3311 |
| LPC 20:2 | 4.5858 | 2.1972 | 0.00042051 | 0.0085011 | 2.0705 |
| LPC 20:3 | 3.4791 | 1.7987 | 0.00014069 | 0.0061905 | 2.2083 |
| LPC 20:4 | 2.4976 | 1.3205 | 0.00051551 | 0.0086482 | 2.0631 |

| | | | | | |
|---|---|---|---|---|---|
| LPC 20:5 | 2.9095 | 1.5408 | 0.0019061 | 0.016202 | 1.7904 |
| LPC 22:0 | 3.2839 | 1.7154 | 0.0019061 | 0.016202 | 1.7904 |
| LPC 22:1 | 4.0597 | 2.0214 | 1.76E-05 | 0.0032935 | 2.4823 |
| LPC 22:2 | 5.7336 | 2.5194 | 0.0006293 | 0.0098066 | 2.0085 |
| LPC 22:3 | 6.8242 | 2.7707 | 0.0031534 | 0.02016 | 1.6955 |
| LPC 22:4 | 3.6425 | 1.8649 | 0.00042051 | 0.0085011 | 2.0705 |
| LPC 22:5 | 3.0966 | 1.6307 | 0.00092529 | 0.011163 | 1.9522 |
| LPC 22:6 | 3.0733 | 1.6198 | 0.00022148 | 0.0075304 | 2.1232 |
| LPC 24:0 | 4.3192 | 2.1108 | 0.0022621 | 0.017444 | 1.7584 |
| LPC 24:1 | 4.1746 | 2.0616 | 4.06E-05 | 0.0043427 | 2.3622 |
| LPC 24:2 | 5.5842 | 2.4813 | 0.0058902 | 0.029972 | 1.5233 |
| LPC 24:4 | 4.9911 | 2.3194 | 0.0037044 | 0.022712 | 1.6437 |
| LPC 24:5 | 5.0298 | 2.3305 | 0.00052028 | 0.0086482 | 2.0631 |
| LPC 26:1 | 4.2194 | 2.077 | 0.0019061 | 0.016202 | 1.7904 |
| LPC 26:2 | 5.2516 | 2.3927 | 0.015684 | 0.052844 | 1.277 |
| LPC 26:4 | 8.1029 | 3.0184 | 0.077097 | 0.14787 | 0.83012 |
| LPC O-14:0 | 5.5599 | 2.4751 | 0.0043372 | 0.025346 | 1.5961 |
| LPC O-15:0 | 5.5457 | 2.4714 | 0.00027569 | 0.0076376 | 2.117 |
| LPC O-16:0 | 8.0593 | 3.0107 | 0.0026755 | 0.01888 | 1.724 |
| LPC O-16:1 | 4.3564 | 2.1231 | 0.00027569 | 0.0076376 | 2.117 |
| LPC O-17:0 | 7.2136 | 2.8507 | 0.0031534 | 0.02016 | 1.6955 |
| LPC O-18:0 | 8.8691 | 3.1488 | 0.00042051 | 0.0085011 | 2.0705 |
| LPC O-18:1 | 7.5166 | 2.9101 | 0.00034126 | 0.0085011 | 2.0705 |
| LPC O-18:2 | 5.9466 | 2.5721 | 0.00092529 | 0.011163 | 1.9522 |
| LPC O-19:0 | 4.7626 | 2.2517 | 0.012021 | 0.044075 | 1.3558 |
| LPC O-20:0 | 8.2007 | 3.0357 | 0.00051551 | 0.0086482 | 2.0631 |
| LPC O-20:1 | 11.476 | 3.5205 | 0.00042051 | 0.0085011 | 2.0705 |
| LPC O-20:2 | 12.57 | 3.6519 | 0.00092529 | 0.011163 | 1.9522 |
| LPC O-21:0 | 11.121 | 3.4752 | 0.0068051 | 0.033626 | 1.4733 |
| LPC O-22:0 | 7.4158 | 2.8906 | 0.00027569 | 0.0076376 | 2.117 |
| LPC O-22:1 | 8.6687 | 3.1158 | 0.00014069 | 0.0061905 | 2.2083 |
| LPC O-22:2 | 5.239 | 2.3893 | 8.73E-05 | 0.0046651 | 2.3311 |
| LPC O-23:0 | 6.1461 | 2.6197 | 0.010482 | 0.041049 | 1.3867 |
| LPC O-24:0 | 2.4328 | 1.2826 | 0.00092529 | 0.011163 | 1.9522 |
| LPC O-24:1 | 10.484 | 3.3901 | 0.0011149 | 0.012636 | 1.8984 |
| LPC O-24:2 | 11.803 | 3.5611 | 0.00076461 | 0.010791 | 1.9669 |
| LPC O-26:2 | 7.7173 | 2.9481 | 0.0050625 | 0.02744 | 1.5616 |
| LPE 16:0 | 1.9943 | 0.99586 | 0.0058902 | 0.029972 | 1.5233 |
| LPE 18:0 | 1.9232 | 0.94352 | 0.0019061 | 0.016202 | 1.7904 |
| LPE 18:2 | 1.3335 | 0.4152 | 0.045875 | 0.10004 | 0.99982 |
| LPE 20:3 | 1.8136 | 0.85889 | 0.012021 | 0.044075 | 1.3558 |
| LPE 20:4 | 0.96429 | -0.052467 | 0.62087 | 0.68699 | 0.16305 |
| LPE 22:0 | 2.3147 | 1.2108 | 0.02025 | 0.060588 | 1.2176 |
| LPE 22:4 | 0.82551 | -0.27664 | 0.81318 | 0.8519 | 0.06961 |

| | | | | | |
|---|---|---|---|---|---|
| LPE O-16:0 | 5.2244 | 2.3853 | 0.03277 | 0.080368 | 1.0949 |
| LPE O-16:1 | 2.5752 | 1.3647 | 0.0013383 | 0.014099 | 1.8508 |
| LPE O-17:1 | 2.2153 | 1.1475 | 0.013749 | 0.04874 | 1.3121 |
| LPE O-18:1 | 2.098 | 1.069 | 0.0026755 | 0.01888 | 1.724 |
| LPE O-18:2 | 1.8658 | 0.8998 | 0.0026755 | 0.01888 | 1.724 |
| LPE O-19:0 | 6.7728 | 2.7597 | 0.00042051 | 0.0085011 | 2.0705 |
| LPE O-20:1 | 2.781 | 1.4756 | 6.82E-05 | 0.0046651 | 2.3311 |
| LPE O-22:1 | 2.279 | 1.1884 | 0.00076461 | 0.010791 | 1.9669 |
| LPG 18:0 | 2.3429 | 1.2283 | 0.029159 | 0.073935 | 1.1311 |
| LPG 18:1 | 1.6048 | 0.68242 | 0.12252 | 0.20877 | 0.68034 |
| LPG 18:2 | 1.2298 | 0.29837 | 0.45087 | 0.55378 | 0.25666 |
| LPI 16:0 | 3.2588 | 1.7044 | 0.0013383 | 0.014099 | 1.8508 |
| LPI 18:0 | 2.4736 | 1.3066 | 0.0022621 | 0.017444 | 1.7584 |
| LPI 18:1 | 2.3034 | 1.2037 | 0.0031534 | 0.02016 | 1.6955 |
| LPI 18:2 | 2.6239 | 1.3917 | 0.0019061 | 0.016202 | 1.7904 |
| LPI 20:3 | 2.1305 | 1.0912 | 0.0037044 | 0.022712 | 1.6437 |
| LPI 20:4 | 2.0384 | 1.0274 | 0.0058902 | 0.029972 | 1.5233 |
| LPS 18:0 | 1.751 | 0.80815 | 0.23401 | 0.33468 | 0.47537 |
| Palmitoyl ethanolamide | 4.065 | 2.0232 | 4.06E-05 | 0.0043427 | 2.3622 |
| PC 14:0_16:1 | 1.3515 | 0.43459 | 0.45087 | 0.55378 | 0.25666 |
| PC 14:0_20:4 | 1.5283 | 0.61194 | 0.1337 | 0.22224 | 0.65318 |
| PC 15:0_20:4 | 1.2893 | 0.36656 | 0.2012 | 0.3004 | 0.5223 |
| PC 16:0_18:1 | 1.1824 | 0.24167 | 0.28978 | 0.38984 | 0.40911 |
| PC 16:0_18:2 | 1.5075 | 0.59213 | 0.010482 | 0.041049 | 1.3867 |
| PC 16:0_18:3 | 1.7668 | 0.8211 | 0.015684 | 0.052844 | 1.277 |
| PC 16:0_20:4 | 1.2298 | 0.29844 | 0.2703 | 0.37166 | 0.42985 |
| PC 16:0_22:6 | 1.7774 | 0.82981 | 0.1337 | 0.22224 | 0.65318 |
| PC 16:0_24:0 | 4.4265 | 2.1462 | 0.00071752 | 0.010791 | 1.9669 |
| PC 16:1_18:2 | 1.5592 | 0.64079 | 0.71483 | 0.76494 | 0.11637 |
| PC 16:1_20:4 | 1.3171 | 0.39732 | 0.2012 | 0.3004 | 0.5223 |
| PC 17:0_18:0 | 1.8252 | 0.86803 | 0.010482 | 0.041049 | 1.3867 |
| PC 17:0_18:1 | 1.086 | 0.11896 | 0.31014 | 0.41206 | 0.38504 |
| PC 17:0_20:4 | 1.1177 | 0.1605 | 0.35355 | 0.45675 | 0.34032 |
| PC 17:1_20:4 | 1.0554 | 0.077771 | 0.91448 | 0.93446 | 0.029437 |
| PC 18:0_18:1 | 1.4084 | 0.49408 | 0.022923 | 0.063978 | 1.194 |
| PC 18:0_18:2 | 1.5419 | 0.62468 | 0.010482 | 0.041049 | 1.3867 |
| PC 18:0_20:2 | 2.2364 | 1.1612 | 0.0019061 | 0.016202 | 1.7904 |
| PC 18:0_20:3 | 1.7587 | 0.81448 | 0.029159 | 0.073935 | 1.1311 |
| PC 18:0_20:4 | 1.4192 | 0.50513 | 0.045875 | 0.10004 | 0.99982 |
| PC 18:0_20:5 | 1.7937 | 0.84293 | 0.0026755 | 0.01888 | 1.724 |
| PC 18:0_22:4 | 2.1316 | 1.092 | 0.045875 | 0.10004 | 0.99982 |
| PC 18:0_22:5 | 2.1325 | 1.0925 | 0.015684 | 0.052844 | 1.277 |
| PC 18:0_22:6 | 2.2399 | 1.1634 | 0.00076461 | 0.010791 | 1.9669 |
| PC 18:0_24:4 | 1.8482 | 0.88614 | 0.056773 | 0.11635 | 0.93425 |

| | | | | | |
|---|---|---|---|---|---|
| PC 18:0_24:5 | 1.3169 | 0.39719 | 0.25171 | 0.35325 | 0.45192 |
| PC 18:1_18:1 | 1.11 | 0.15049 | 0.40049 | 0.50348 | 0.29802 |
| PC 18:1_18:2 | 1.2279 | 0.2962 | 0.37659 | 0.47988 | 0.31887 |
| PC 18:1_20:1 | 1.2056 | 0.26981 | 0.18608 | 0.28464 | 0.5457 |
| PC 18:1_22:6 | 1.5526 | 0.63472 | 0.084761 | 0.1585 | 0.79996 |
| PC 18:1_24:4 | 1.2728 | 0.34803 | 0.94862 | 0.95758 | 0.018823 |
| PC 18:2_18:2 | 1.5524 | 0.63453 | 0.1337 | 0.22224 | 0.65318 |
| PC 19:0_20:4 | 1.6049 | 0.68245 | 0.022923 | 0.063978 | 1.194 |
| PC 20:0_20:4 | 1.5417 | 0.62448 | 0.052091 | 0.11038 | 0.95711 |
| PC 20:2_20:4 | 1.3203 | 0.40091 | 0.21718 | 0.31666 | 0.4994 |
| PC 20:3_20:4 | 1.494 | 0.57923 | 0.11112 | 0.1945 | 0.71109 |
| PC 20:4_22:6 | 1.4804 | 0.56602 | 0.022923 | 0.063978 | 1.194 |
| PC 20:4_24:4 | 0.89157 | -0.16558 | 0.47731 | 0.57771 | 0.23829 |
| PC 22:1_20:4 | 1.575 | 0.65539 | 0.0079031 | 0.035398 | 1.451 |
| PC 24:0 | 0.97212 | -0.040792 | 0.45807 | 0.56078 | 0.25121 |
| PC 24:0_18:1 | 2.0777 | 1.055 | 0.025885 | 0.068659 | 1.1633 |
| PC 24:0_18:2 | 2.9689 | 1.5699 | 0.0031534 | 0.02016 | 1.6955 |
| PC 26:0 | 1.5316 | 0.61504 | 0.11112 | 0.1945 | 0.71109 |
| PC 28:0 | 1.6927 | 0.75934 | 0.23401 | 0.33468 | 0.47537 |
| PC 28:1 | 1.2389 | 0.30908 | 0.52679 | 0.61759 | 0.2093 |
| PC 29:0 | 1.4904 | 0.57574 | 0.62087 | 0.68699 | 0.16305 |
| PC 30:0 | 1.0472 | 0.066578 | 0.71483 | 0.76494 | 0.11637 |
| PC 30:2 | 1.61 | 0.68706 | 0.53255 | 0.61759 | 0.2093 |
| PC 31:0 | 1.0356 | 0.05052 | 0.77999 | 0.82523 | 0.083426 |
| PC 31:1 | 1.2459 | 0.31719 | 0.68295 | 0.74144 | 0.12993 |
| PC 32:0 | 1.2146 | 0.28043 | 0.10231 | 0.18134 | 0.7415 |
| PC 32:1 | 1.2067 | 0.2711 | 0.59074 | 0.66648 | 0.17621 |
| PC 32:2 | 1.6484 | 0.72103 | 0.14562 | 0.23526 | 0.62846 |
| PC 33:0 | 1.018 | 0.025746 | 0.71483 | 0.76494 | 0.11637 |
| PC 33:1 | 1.2163 | 0.28254 | 0.2012 | 0.3004 | 0.5223 |
| PC 33:2 | 1.3623 | 0.44602 | 0.3314 | 0.43413 | 0.36238 |
| PC 34:0 | 1.438 | 0.52402 | 0.022923 | 0.063978 | 1.194 |
| PC 34:5 | 1.6038 | 0.68149 | 0.1337 | 0.22224 | 0.65318 |
| PC 35:0 | 1.5409 | 0.62377 | 0.15831 | 0.25088 | 0.60053 |
| PC 35:1 | 1.1703 | 0.22683 | 0.2703 | 0.37166 | 0.42985 |
| PC 35:2 | 1.3734 | 0.45771 | 0.11207 | 0.1945 | 0.71109 |
| PC 36:0 | 2.0659 | 1.0467 | 0.00092529 | 0.011163 | 1.9522 |
| PC 36:3 | 1.6374 | 0.71137 | 0.017844 | 0.056316 | 1.2494 |
| PC 36:5 | 1.9363 | 0.9533 | 0.017844 | 0.056316 | 1.2494 |
| PC 36:6 | 1.9093 | 0.93304 | 0.05109 | 0.10857 | 0.96431 |
| PC 37:1 | 1.7866 | 0.83722 | 0.02025 | 0.060588 | 1.2176 |
| PC 37:2 | 1.2411 | 0.31163 | 0.17179 | 0.26826 | 0.57144 |
| PC 37:6 | 2.0185 | 1.0133 | 0.045875 | 0.10004 | 0.99982 |
| PC 38:1 | 1.8852 | 0.91475 | 0.0013383 | 0.014099 | 1.8508 |

| | | | | | |
|---|---|---|---|---|---|
| PC 38:4 | 1.4405 | 0.52656 | 0.084761 | 0.1585 | 0.79996 |
| PC 38:5 | 1.2353 | 0.30482 | 0.15831 | 0.25088 | 0.60053 |
| PC 39:1 | 1.434 | 0.52009 | 0.47731 | 0.57771 | 0.23829 |
| PC 39:2 | 0.99472 | -0.0076339 | 0.82713 | 0.86531 | 0.062829 |
| PC 39:3 | 1.5379 | 0.62094 | 0.1337 | 0.22224 | 0.65318 |
| PC 39:5 | 1.3283 | 0.4096 | 0.18608 | 0.28464 | 0.5457 |
| PC 39:6 | 2.1584 | 1.11 | 0.0079031 | 0.035398 | 1.451 |
| PC 40:1 | 2.3548 | 1.2356 | 0.0013383 | 0.014099 | 1.8508 |
| PC 40:2 | 1.6697 | 0.73961 | 0.0079031 | 0.035398 | 1.451 |
| PC 40:5 | 1.5993 | 0.67746 | 0.036742 | 0.085617 | 1.0674 |
| PC 40:8 | 1.8962 | 0.92308 | 0.0011149 | 0.012636 | 1.8984 |
| PC 40:9 | 1.8768 | 0.9083 | 0.0077467 | 0.035398 | 1.451 |
| PC 42:2 | 1.9112 | 0.93446 | 0.03277 | 0.080368 | 1.0949 |
| PC 42:3 | 2.1969 | 1.1355 | 0.0037044 | 0.022712 | 1.6437 |
| PC 42:6 | 1.6828 | 0.75086 | 0.02025 | 0.060588 | 1.2176 |
| PC 42:8 | 1.2018 | 0.26525 | 0.65162 | 0.71259 | 0.14716 |
| PC 42:9 | 1.5422 | 0.62497 | 0.015684 | 0.052844 | 1.277 |
| PC 43:9 | 0.96608 | -0.049781 | 0.87858 | 0.91022 | 0.040855 |
| PC 44:10 | 1.4237 | 0.50962 | 0.056773 | 0.11635 | 0.93425 |
| PC 44:11 | 1.5125 | 0.59691 | 0.14363 | 0.23526 | 0.62846 |
| PC 44:12 | 1.8085 | 0.85478 | 0.15831 | 0.25088 | 0.60053 |
| PC 44:3 | 2.9163 | 1.5441 | 0.11207 | 0.1945 | 0.71109 |
| PC 44:5 | 1.8571 | 0.89302 | 0.013749 | 0.04874 | 1.3121 |
| PC 44:6 | 1.5336 | 0.61695 | 0.58529 | 0.66648 | 0.17621 |
| PC 46:8 | 0.7726 | -0.3722 | 0.41938 | 0.52633 | 0.27874 |
| PC O-16:0_20:3 | 1.6448 | 0.71794 | 0.010482 | 0.041049 | 1.3867 |
| PC O-16:0_20:5 | 1.3928 | 0.47797 | 0.42526 | 0.52928 | 0.27632 |
| PC O-16:0_22:4 | 1.8042 | 0.85139 | 0.029159 | 0.073935 | 1.1311 |
| PC O-16:0_22:6 | 1.2332 | 0.30241 | 0.21718 | 0.31666 | 0.4994 |
| PC O-16:1_20:5 | 1.2292 | 0.29776 | 0.88048 | 0.91092 | 0.040518 |
| PC O-16:1_22:4 | 1.5592 | 0.64083 | 0.0026755 | 0.01888 | 1.724 |
| PC O-18:0_16:0 | 1.4042 | 0.48979 | 0.15831 | 0.25088 | 0.60053 |
| PC O-18:0_18:2 | 1.9507 | 0.96398 | 0.02025 | 0.060588 | 1.2176 |
| PC O-18:0_20:4 | 1.866 | 0.89991 | 0.0091146 | 0.038959 | 1.4094 |
| PC O-18:0_22:6 | 1.5789 | 0.65891 | 0.029159 | 0.073935 | 1.1311 |
| PC O-18:1_18:1 | 1.4376 | 0.52369 | 0.25171 | 0.35325 | 0.45192 |
| PC O-18:1_20:4 | 1.3796 | 0.46427 | 0.029159 | 0.073935 | 1.1311 |
| PC O-18:1_22:4 | 1.6386 | 0.71244 | 0.036742 | 0.085617 | 1.0674 |
| PC O-18:1_22:5 | 1.6586 | 0.72998 | 0.0019061 | 0.016202 | 1.7904 |
| PC O-18:1_22:6 | 1.5033 | 0.58814 | 0.041102 | 0.093733 | 1.0281 |
| PC O-20:0_20:4 | 2.149 | 1.1037 | 0.0022621 | 0.017444 | 1.7584 |
| PC O-22:0_20:4 | 2.4041 | 1.2655 | 0.0031534 | 0.02016 | 1.6955 |
| PC O-22:1_20:4 | 2.6784 | 1.4214 | 0.0006293 | 0.0098066 | 2.0085 |
| PC O-24:0_20:4 | 2.3103 | 1.2081 | 0.0022621 | 0.017444 | 1.7584 |

| | | | | | |
|---|---|---|---|---|---|
| PC O-24:1_20:4 | 2.9278 | 1.5498 | 0.00017708 | 0.0073586 | 2.1332 |
| PC O-24:1_22:4 | 3.5207 | 1.8159 | 0.014512 | 0.051202 | 1.2907 |
| PC O-24:2_20:4 | 2.7666 | 1.4681 | 0.00092529 | 0.011163 | 1.9522 |
| PC O-24:3_20:4 | 2.4986 | 1.3211 | 0.0079031 | 0.035398 | 1.451 |
| PC O-28:0 | 1.9138 | 0.93646 | 0.15831 | 0.25088 | 0.60053 |
| PC O-30:0 | 1.656 | 0.72774 | 0.14562 | 0.23526 | 0.62846 |
| PC O-30:1 | 1.3282 | 0.40947 | 0.50454 | 0.59904 | 0.22254 |
| PC O-31:0 | 1.3906 | 0.47573 | 0.62087 | 0.68699 | 0.16305 |
| PC O-31:1 | 1.1358 | 0.18367 | 0.93043 | 0.94947 | 0.022516 |
| PC O-32:0 | 1.6762 | 0.74521 | 0.03277 | 0.080368 | 1.0949 |
| PC O-32:1 | 1.7877 | 0.83806 | 0.022923 | 0.063978 | 1.194 |
| PC O-32:4 | 1.1194 | 0.16267 | 0.40049 | 0.50348 | 0.29802 |
| PC O-33:1 | 1.4942 | 0.57936 | 0.28978 | 0.38984 | 0.40911 |
| PC O-34:1 | 1.4196 | 0.50547 | 0.10231 | 0.18134 | 0.7415 |
| PC O-34:2 | 1.4491 | 0.5352 | 0.05109 | 0.10857 | 0.96431 |
| PC O-34:3 | 1.6927 | 0.75933 | 0.041102 | 0.093733 | 1.0281 |
| PC O-34:4 | 1.6415 | 0.71503 | 0.10231 | 0.18134 | 0.7415 |
| PC O-34:5 | 1.5064 | 0.59108 | 0.91448 | 0.93446 | 0.029437 |
| PC O-35:2 | 1.5953 | 0.67383 | 0.21718 | 0.31666 | 0.4994 |
| PC O-36:0 | 2.5092 | 1.3272 | 0.0043372 | 0.025346 | 1.5961 |
| PC O-36:1 | 2.0422 | 1.0301 | 0.015684 | 0.052844 | 1.277 |
| PC O-36:4 | 1.5896 | 0.66864 | 0.017844 | 0.056316 | 1.2494 |
| PC O-36:5 | 1.1154 | 0.15756 | 0.53255 | 0.61759 | 0.2093 |
| PC O-37:4 | 1.4159 | 0.50174 | 0.069659 | 0.13712 | 0.86291 |
| PC O-37:5 | 1.052 | 0.0731 | 0.71483 | 0.76494 | 0.11637 |
| PC O-38:0 | 3.1325 | 1.6473 | 0.0006293 | 0.0098066 | 2.0085 |
| PC O-38:2 | 2.3015 | 1.2026 | 0.010482 | 0.041049 | 1.3867 |
| PC O-38:3 | 3.0782 | 1.6221 | 8.73E-05 | 0.0046651 | 2.3311 |
| PC O-38:7 | 1.4419 | 0.52797 | 0.03277 | 0.080368 | 1.0949 |
| PC O-39:7 | 1.3103 | 0.38993 | 0.59074 | 0.66648 | 0.17621 |
| PC O-40:1 | 2.8763 | 1.5242 | 0.00092529 | 0.011163 | 1.9522 |
| PC O-40:4 | 2.4214 | 1.2758 | 0.0016 | 0.015748 | 1.8028 |
| PC O-40:5 | 1.9929 | 0.99485 | 0.00092529 | 0.011163 | 1.9522 |
| PC O-40:8 | 1.398 | 0.48337 | 0.056773 | 0.11635 | 0.93425 |
| PC O-42:10 | 1.0496 | 0.069877 | 0.74719 | 0.79615 | 0.099003 |
| PC O-42:6 | 2.4125 | 1.2706 | 0.0058902 | 0.029972 | 1.5233 |
| PC O-42:8 | 1.9373 | 0.95406 | 0.044659 | 0.10004 | 0.99982 |
| PC O-43:5 | 2.8497 | 1.5108 | 0.0013357 | 0.014099 | 1.8508 |
| PC O-44:4 | 2.9404 | 1.556 | 0.00092529 | 0.011163 | 1.9522 |
| PC O-44:6 | 2.1586 | 1.1101 | 0.0050625 | 0.02744 | 1.5616 |
| PC O-44:7 | 3.0601 | 1.6136 | 0.00051551 | 0.0086482 | 2.0631 |
| PC O-44:8 | 2.0813 | 1.0575 | 0.029159 | 0.073935 | 1.1311 |
| PC O-46:5 | 2.453 | 1.2945 | 0.00027569 | 0.0076376 | 2.117 |
| PC O-46:6 | 3.0603 | 1.6136 | 0.00014069 | 0.0061905 | 2.2083 |

| | | | | | |
|---|---|---|---|---|---|
| PC O-46:7 | 3.0626 | 1.6147 | 0.00051551 | 0.0086482 | 2.0631 |
| PC O-46:8 | 2.8936 | 1.5329 | 0.0037044 | 0.022712 | 1.6437 |
| PE 16:0_16:0 | 0.94848 | -0.076311 | 0.56129 | 0.64197 | 0.19249 |
| PE 16:0_18:0 | 1.0716 | 0.099789 | 0.81318 | 0.8519 | 0.06961 |
| PE 16:0_18:1 | 0.76554 | -0.38545 | 0.013749 | 0.04874 | 1.3121 |
| PE 16:0_18:2 | 0.85454 | -0.22677 | 0.2703 | 0.37166 | 0.42985 |
| PE 16:0_20:4 | 1.0548 | 0.076962 | 0.91448 | 0.93446 | 0.029437 |
| PE 16:1_18:1 | 1.4545 | 0.54052 | 0.98286 | 0.9855 | 0.0063434 |
| PE 17:0_18:2 | 0.77656 | -0.36482 | 0.3314 | 0.43413 | 0.36238 |
| PE 18:0_18:1 | 0.77126 | -0.3747 | 0.10231 | 0.18134 | 0.7415 |
| PE 18:0_20:4 | 0.85944 | -0.21853 | 0.14562 | 0.23526 | 0.62846 |
| PE 18:0_22:6 | 1.1155 | 0.1577 | 0.37659 | 0.47988 | 0.31887 |
| PE 18:0_24:4 | 0.87673 | -0.18979 | 0.47731 | 0.57771 | 0.23829 |
| PE 18:1_18:1 | 0.77908 | -0.36016 | 0.076919 | 0.14787 | 0.83012 |
| PE 18:1_20:1 | 0.65731 | -0.60536 | 0.025885 | 0.068659 | 1.1633 |
| PE 18:1_20:4 | 0.88442 | -0.17719 | 0.35355 | 0.45675 | 0.34032 |
| PE 24:0_18:2 | 1.2625 | 0.3363 | 0.42526 | 0.52928 | 0.27632 |
| PE 32:1 | 0.97245 | -0.040307 | 0.53255 | 0.61759 | 0.2093 |
| PE 38:1 | 0.75493 | -0.40559 | 0.12252 | 0.20877 | 0.68034 |
| PE 38:6 | 1.0738 | 0.10272 | 0.94862 | 0.95758 | 0.018823 |
| PE 40:4 | 1.1848 | 0.24467 | 0.91448 | 0.93446 | 0.029437 |
| PE 40:7 | 1.0134 | 0.019215 | 0.74719 | 0.79615 | 0.099003 |
| PE 40:8 | 1.0099 | 0.014155 | 0.62087 | 0.68699 | 0.16305 |
| PE O-15:1_20:4 | 1.0743 | 0.10346 | 0.91448 | 0.93446 | 0.029437 |
| PE O-16:0_22:4 | 0.84293 | -0.24652 | 0.2012 | 0.3004 | 0.5223 |
| PE O-16:1_16:0 | 0.8485 | -0.23702 | 0.45087 | 0.55378 | 0.25666 |
| PE O-16:1_16:1 | 0.7329 | -0.44832 | 0.022923 | 0.063978 | 1.194 |
| PE O-16:1_18:1 | 0.85338 | -0.22873 | 0.2703 | 0.37166 | 0.42985 |
| PE O-16:1_20:3 | 0.77865 | -0.36095 | 0.093216 | 0.16924 | 0.77151 |
| PE O-16:1_20:4 | 0.78857 | -0.34269 | 0.1337 | 0.22224 | 0.65318 |
| PE O-16:1_20:5 | 0.96609 | -0.049776 | 0.2703 | 0.37166 | 0.42985 |
| PE O-16:1_22:4 | 0.92286 | -0.11581 | 0.42526 | 0.52928 | 0.27632 |
| PE O-16:1_22:5 | 0.93697 | -0.093919 | 0.62087 | 0.68699 | 0.16305 |
| PE O-16:1_22:6 | 0.82946 | -0.26976 | 0.56129 | 0.64197 | 0.19249 |
| PE O-16:1_24:4 | 0.56219 | -0.83086 | 0.025885 | 0.068659 | 1.1633 |
| PE O-17:1_18:2 | 0.95306 | -0.069361 | 0.45087 | 0.55378 | 0.25666 |
| PE O-17:1_22:4 | 0.65128 | -0.61864 | 0.0050625 | 0.02744 | 1.5616 |
| PE O-17:1_22:5 | 0.5834 | -0.77744 | 0.036742 | 0.085617 | 1.0674 |
| PE O-17:1_22:6 | 0.7184 | -0.47714 | 0.076919 | 0.14787 | 0.83012 |
| PE O-18:0_18:1 | 0.84954 | -0.23525 | 0.25171 | 0.35325 | 0.45192 |
| PE O-18:0_18:2 | 1.9778 | 0.98389 | 0.017844 | 0.056316 | 1.2494 |
| PE O-18:0_22:4 | 0.7642 | -0.38799 | 0.28978 | 0.38984 | 0.40911 |
| PE O-18:1_18:0 | 0.93628 | -0.094986 | 0.98286 | 0.9855 | 0.0063434 |
| PE O-18:1_18:1 | 0.7125 | -0.48905 | 0.14562 | 0.23526 | 0.62846 |

| | | | | | |
|---|---|---|---|---|---|
| PE O-18:1_18:2 | 0.86726 | -0.20546 | 0.3314 | 0.43413 | 0.36238 |
| PE O-18:1_20:3 | 0.82788 | -0.2725 | 0.25171 | 0.35325 | 0.45192 |
| PE O-18:1_20:4 | 0.78076 | -0.35704 | 0.18608 | 0.28464 | 0.5457 |
| PE O-18:1_22:2 | 0.94253 | -0.085382 | 0.45087 | 0.55378 | 0.25666 |
| PE O-18:1_22:3 | 0.89772 | -0.15567 | 0.50454 | 0.59904 | 0.22254 |
| PE O-18:1_22:4 | 0.88329 | -0.17905 | 0.50454 | 0.59904 | 0.22254 |
| PE O-18:1_22:5 | 0.81633 | -0.29277 | 0.28978 | 0.38984 | 0.40911 |
| PE O-18:1_22:6 | 0.63602 | -0.65286 | 0.025885 | 0.068659 | 1.1633 |
| PE O-18:1_24:4 | 0.52691 | -0.92437 | 0.02025 | 0.060588 | 1.2176 |
| PE O-18:2_18:1 | 0.75531 | -0.40485 | 0.045875 | 0.10004 | 0.99982 |
| PE O-18:2_18:2 | 0.88128 | -0.18232 | 0.47731 | 0.57771 | 0.23829 |
| PE O-18:2_20:5 | 0.85616 | -0.22406 | 0.21718 | 0.31666 | 0.4994 |
| PE O-18:2_22:6 | 0.77896 | -0.36039 | 0.23401 | 0.33468 | 0.47537 |
| PE O-20:0_16:0 | 1.5405 | 0.62338 | 0.23401 | 0.33468 | 0.47537 |
| PE O-20:0_20:4 | 2.1656 | 1.1148 | 0.0016 | 0.015748 | 1.8028 |
| PE O-20:1_20:4 | 0.99664 | -0.004862 | 0.94862 | 0.95758 | 0.018823 |
| PE O-20:1_22:6 | 1.0309 | 0.043885 | 0.56129 | 0.64197 | 0.19249 |
| PE O-20:2_20:4 | 0.7038 | -0.50676 | 0.084761 | 0.1585 | 0.79996 |
| PE O-22:1_20:4 | 1.5829 | 0.66258 | 0.28978 | 0.38984 | 0.40911 |
| PE O-22:2_20:4 | 0.88616 | -0.17436 | 0.68295 | 0.74144 | 0.12993 |
| PE O-24:1_20:4 | 1.3005 | 0.3791 | 0.77999 | 0.82523 | 0.083426 |
| PE O-37:5 | 0.65973 | -0.60006 | 0.022923 | 0.063978 | 1.194 |
| PE O-38:2 | 0.94259 | -0.08529 | 0.84669 | 0.8784 | 0.056307 |
| PE O-38:4 | 0.84241 | -0.24741 | 0.50454 | 0.59904 | 0.22254 |
| PE O-40:8 | 1.9836 | 0.98809 | 0.022923 | 0.063978 | 1.194 |
| PE-Cer (d32:1) | 4.6223 | 2.2086 | 0.010482 | 0.041049 | 1.3867 |
| PE-Cer (d33:1) | 3.4473 | 1.7854 | 0.012021 | 0.044075 | 1.3558 |
| PE-Cer (d34:0) | 3.7501 | 1.9069 | 0.02025 | 0.060588 | 1.2176 |
| PE-Cer (d34:1) | 2.8299 | 1.5008 | 0.0091146 | 0.038959 | 1.4094 |
| PE-Cer (d34:2) | 4.2577 | 2.0901 | 0.0050625 | 0.02744 | 1.5616 |
| PE-Cer (d36:1) | 1.6098 | 0.68684 | 0.1337 | 0.22224 | 0.65318 |
| PE-Cer (d36:2) | 1.6737 | 0.74303 | 0.093216 | 0.16924 | 0.77151 |
| PE-Cer (d38:1) | 2.5748 | 1.3644 | 0.010482 | 0.041049 | 1.3867 |
| PE-Cer (d40:2) | 2.6548 | 1.4086 | 0.0022621 | 0.017444 | 1.7584 |
| PE-Cer (d42:2) | 1.905 | 0.92981 | 0.017844 | 0.056316 | 1.2494 |
| PE-Cer (d42:3) | 2.5259 | 1.3368 | 0.045875 | 0.10004 | 0.99982 |
| PG 16:0_18:2 | 0.63602 | -0.65287 | 0.013749 | 0.04874 | 1.3121 |
| PG 16:0_20:4 | 0.86802 | -0.2042 | 0.53255 | 0.61759 | 0.2093 |
| PG 18:1_18:1 | 1.4562 | 0.54223 | 0.14562 | 0.23526 | 0.62846 |
| PG 18:1_18:2 | 1.6706 | 0.74036 | 0.3314 | 0.43413 | 0.36238 |
| PG 18:1_20:4 | 1.2986 | 0.37698 | 0.59074 | 0.66648 | 0.17621 |
| PG 18:1_22:5 | 1.3592 | 0.44275 | 0.47731 | 0.57771 | 0.23829 |
| PG 18:2_20:4 | 1.1373 | 0.1856 | 0.71483 | 0.76494 | 0.11637 |
| PG 18:2_22:5 | 0.96116 | -0.057149 | 0.28978 | 0.38984 | 0.40911 |

| | | | | | |
|---|---|---|---|---|---|
| PG 18:2_22:6 | 1.2658 | 0.34002 | 0.84669 | 0.8784 | 0.056307 |
| PG 20:4_22:4 | 1.84 | 0.87969 | 0.31014 | 0.41206 | 0.38504 |
| PG 20:4_22:6 | 1.4427 | 0.52881 | 0.77999 | 0.82523 | 0.083426 |
| PG 22:4_22:6 | 2.2878 | 1.194 | 0.23401 | 0.33468 | 0.47537 |
| PG 22:5_22:5 | 3.3496 | 1.744 | 0.013749 | 0.04874 | 1.3121 |
| PG 22:5_22:6 | 1.4489 | 0.53497 | 0.28978 | 0.38984 | 0.40911 |
| PG 22:6_22:6 | 1.7562 | 0.81242 | 0.94862 | 0.95758 | 0.018823 |
| PG 40:7 | 1.6937 | 0.76019 | 0.53255 | 0.61759 | 0.2093 |
| PI 16:0_16:0 | 2.955 | 1.5631 | 0.013749 | 0.04874 | 1.3121 |
| PI 16:0_16:1 | 1.5786 | 0.65862 | 0.25171 | 0.35325 | 0.45192 |
| PI 16:0_20:3 | 0.91607 | -0.12647 | 0.45087 | 0.55378 | 0.25666 |
| PI 16:0_20:4 | 0.93014 | -0.10449 | 0.98286 | 0.9855 | 0.0063434 |
| PI 17:0_20:4 | 0.82461 | -0.27822 | 0.50454 | 0.59904 | 0.22254 |
| PI 18:0_18:1 | 1.0084 | 0.01201 | 1 | 1 | 0 |
| PI 18:0_18:2 | 1.2806 | 0.35676 | 0.2012 | 0.3004 | 0.5223 |
| PI 18:0_20:4 | 0.91035 | -0.1355 | 0.56129 | 0.64197 | 0.19249 |
| PI 18:0_22:4 | 0.95107 | -0.072374 | 0.53255 | 0.61759 | 0.2093 |
| PI 18:0_22:6 | 1.908 | 0.93204 | 0.062953 | 0.12624 | 0.89879 |
| PI 18:1_18:1 | 0.95613 | -0.06472 | 0.74719 | 0.79615 | 0.099003 |
| PI 18:1_18:2 | 1.0318 | 0.045197 | 0.91448 | 0.93446 | 0.029437 |
| PI 18:1_22:6 | 1.1769 | 0.23501 | 0.53255 | 0.61759 | 0.2093 |
| PI 20:0_20:4 | 0.80763 | -0.30823 | 0.35355 | 0.45675 | 0.34032 |
| PI 34:1 | 1.2671 | 0.34153 | 0.23401 | 0.33468 | 0.47537 |
| PI 34:2 | 1.2826 | 0.35912 | 0.37659 | 0.47988 | 0.31887 |
| PI 38:2 | 1.0338 | 0.047941 | 0.94862 | 0.95758 | 0.018823 |
| PI 40:5 | 1.0295 | 0.041938 | 0.84669 | 0.8784 | 0.056307 |
| PI O-18:1_20:4 | 0.76814 | -0.38056 | 0.12252 | 0.20877 | 0.68034 |
| PS 16:0_18:0 | 1.3821 | 0.4669 | 0.12252 | 0.20877 | 0.68034 |
| PS 16:0_18:1 | 0.86462 | -0.20986 | 0.40049 | 0.50348 | 0.29802 |
| PS 18:0_22:5 | 1.0789 | 0.10957 | 0.91448 | 0.93446 | 0.029437 |
| PS 18:0_22:6 | 0.9587 | -0.060846 | 0.91448 | 0.93446 | 0.029437 |
| PS 18:1_18:1 | 0.80362 | -0.31541 | 0.2703 | 0.37166 | 0.42985 |
| PS 18:1_22:1 | 0.70915 | -0.49584 | 0.076919 | 0.14787 | 0.83012 |
| PS 34:2 | 0.9103 | -0.13558 | 0.65162 | 0.71259 | 0.14716 |
| PS 36:1 | 0.81678 | -0.29198 | 0.3314 | 0.43413 | 0.36238 |
| PS 36:2 | 0.95816 | -0.061658 | 0.68295 | 0.74144 | 0.12993 |
| PS 38:1 | 0.69177 | -0.53164 | 0.37659 | 0.47988 | 0.31887 |
| PS 38:2 | 0.57383 | -0.8013 | 0.0091146 | 0.038959 | 1.4094 |
| PS 38:3 | 0.85632 | -0.22378 | 0.45087 | 0.55378 | 0.25666 |
| PS 38:4 | 1.3247 | 0.40572 | 0.40049 | 0.50348 | 0.29802 |
| PS 40:1 | 0.71904 | -0.47586 | 0.37659 | 0.47988 | 0.31887 |
| PS 40:4 | 1.2749 | 0.35041 | 0.35355 | 0.45675 | 0.34032 |
| PS 40:5 | 1.2711 | 0.34606 | 0.42526 | 0.52928 | 0.27632 |
| PS 42:1 | 1.1073 | 0.14704 | 0.68295 | 0.74144 | 0.12993 |

| | | | | | |
|---|---|---|---|---|---|
| PS 42:2 | 0.86791 | -0.20439 | 0.53255 | 0.61759 | 0.2093 |
| SM 28:1;O2 | 2.5687 | 1.361 | 0.0091146 | 0.038959 | 1.4094 |
| SM 29:1;O2 | 2.6648 | 1.414 | 0.029057 | 0.073935 | 1.1311 |
| SM 30:0;O2 | 2.211 | 1.1447 | 0.012021 | 0.044075 | 1.3558 |
| SM 30:1;O2 | 2.6515 | 1.4068 | 0.022923 | 0.063978 | 1.194 |
| SM 30:1;O3 | 2.5763 | 1.3653 | 0.0091146 | 0.038959 | 1.4094 |
| SM 30:2;O2 | 3.3082 | 1.7261 | 0.0048723 | 0.02744 | 1.5616 |
| SM 31:0;O2 | 2.4668 | 1.3027 | 0.022923 | 0.063978 | 1.194 |
| SM 31:1;O2 | 2.3488 | 1.2319 | 0.029159 | 0.073935 | 1.1311 |
| SM 31:2 | 2.4672 | 1.3029 | 0.010482 | 0.041049 | 1.3867 |
| SM 32:0;O2 | 2.5895 | 1.3727 | 0.029159 | 0.073935 | 1.1311 |
| SM 32:1;O2 | 2.5049 | 1.3248 | 0.012021 | 0.044075 | 1.3558 |
| SM 32:1;O3 | 3.6252 | 1.858 | 0.0091146 | 0.038959 | 1.4094 |
| SM 32:2;O2 | 2.7995 | 1.4852 | 0.0079031 | 0.035398 | 1.451 |
| SM 32:2;O3 | 3.4377 | 1.7814 | 0.012079 | 0.044075 | 1.3558 |
| SM 33:0;O2 | 2.0192 | 1.0138 | 0.069659 | 0.13712 | 0.86291 |
| SM 33:1;O2 | 2.218 | 1.1493 | 0.022923 | 0.063978 | 1.194 |
| SM 33:1;O3 | 1.7777 | 0.83 | 0.093216 | 0.16924 | 0.77151 |
| SM 33:2;O2 | 3.1157 | 1.6396 | 0.0091146 | 0.038959 | 1.4094 |
| SM 33:2;O3 | 3.7717 | 1.9152 | 0.029159 | 0.073935 | 1.1311 |
| SM 34:0;O2 | 2.3003 | 1.2018 | 0.076919 | 0.14787 | 0.83012 |
| SM 34:1;O3 | 2.5171 | 1.3318 | 0.093216 | 0.16924 | 0.77151 |
| SM 34:2;O2 | 2.5883 | 1.372 | 0.012021 | 0.044075 | 1.3558 |
| SM 34:2;O3 | 4.1495 | 2.0529 | 0.017844 | 0.056316 | 1.2494 |
| SM 34:3;O2 | 2.9897 | 1.58 | 0.017844 | 0.056316 | 1.2494 |
| SM 35:0;O2 | 1.3512 | 0.43428 | 0.50454 | 0.59904 | 0.22254 |
| SM 35:1;O2 | 1.496 | 0.58109 | 0.056773 | 0.11635 | 0.93425 |
| SM 35:1;O3 | 2.42 | 1.275 | 0.036742 | 0.085617 | 1.0674 |
| SM 35:2;O2 | 2.057 | 1.0405 | 0.022923 | 0.063978 | 1.194 |
| SM 36:0;O2 | 1.1226 | 0.16683 | 0.68295 | 0.74144 | 0.12993 |
| SM 36:0;O3 | 1.6072 | 0.68452 | 0.05109 | 0.10857 | 0.96431 |
| SM 36:1;O2 | 1.1951 | 0.25712 | 0.21718 | 0.31666 | 0.4994 |
| SM 36:2;O2 | 1.4564 | 0.5424 | 0.084761 | 0.1585 | 0.79996 |
| SM 36:2;O3 | 1.6153 | 0.69181 | 0.084761 | 0.1585 | 0.79996 |
| SM 36:3;O2 | 2.7557 | 1.4624 | 0.02025 | 0.060588 | 1.2176 |
| SM 37:0;O2 | 1.2217 | 0.28885 | 0.62087 | 0.68699 | 0.16305 |
| SM 37:1;O2 | 1.4211 | 0.50703 | 0.17179 | 0.26826 | 0.57144 |
| SM 37:2;O2 | 1.8218 | 0.8654 | 0.017844 | 0.056316 | 1.2494 |
| SM 38:0;O2 | 1.8082 | 0.85455 | 0.14562 | 0.23526 | 0.62846 |
| SM 38:1;O2 | 1.8292 | 0.87118 | 0.012021 | 0.044075 | 1.3558 |
| SM 38:2;O2 | 2.0405 | 1.0289 | 0.0043372 | 0.025346 | 1.5961 |
| SM 38:4;O2 | 3.3061 | 1.7251 | 0.02025 | 0.060588 | 1.2176 |
| SM 39:1;O2 | 2.7621 | 1.4658 | 0.0022621 | 0.017444 | 1.7584 |
| SM 39:2;O2 | 2.5996 | 1.3783 | 0.0026755 | 0.01888 | 1.724 |

| | | | | | |
|---|---|---|---|---|---|
| SM 40:0;O2 | 1.683 | 0.75104 | 0.15831 | 0.25088 | 0.60053 |
| SM 40:0;O3 | 2.0544 | 1.0387 | 0.041102 | 0.093733 | 1.0281 |
| SM 40:2;O2_1 | 2.0782 | 1.0553 | 0.0079031 | 0.035398 | 1.451 |
| SM 40:2;O2_2 | 2.48 | 1.3103 | 0.0068331 | 0.033626 | 1.4733 |
| SM 40:3;O2 | 2.515 | 1.3306 | 0.012021 | 0.044075 | 1.3558 |
| SM 41:0;O2 | 2.0565 | 1.0402 | 0.0079031 | 0.035398 | 1.451 |
| SM 41:0;O3 | 1.9761 | 0.98266 | 0.0079031 | 0.035398 | 1.451 |
| SM 41:1;O2 | 2.527 | 1.3374 | 0.0026755 | 0.01888 | 1.724 |
| SM 41:1;O3 | 1.9473 | 0.9615 | 0.0079031 | 0.035398 | 1.451 |
| SM 41:2;O2_1 | 2.7339 | 1.451 | 0.0058902 | 0.029972 | 1.5233 |
| SM 41:2;O2_2 | 2.4343 | 1.2835 | 0.010482 | 0.041049 | 1.3867 |
| SM 41:3;O2 | 1.7791 | 0.83113 | 0.056773 | 0.11635 | 0.93425 |
| SM 42:0;O2 | 1.2099 | 0.27494 | 0.56129 | 0.64197 | 0.19249 |
| SM 42:1;O2 | 1.5905 | 0.66952 | 0.10231 | 0.18134 | 0.7415 |
| SM 42:1;O3 | 2.6171 | 1.388 | 0.017844 | 0.056316 | 1.2494 |
| SM 42:2;O2_1 | 1.4657 | 0.55159 | 0.025885 | 0.068659 | 1.1633 |
| SM 42:2;O2_2 | 2.0176 | 1.0126 | 0.029159 | 0.073935 | 1.1311 |
| SM 42:2;O3 | 2.8632 | 1.5176 | 0.0031534 | 0.02016 | 1.6955 |
| SM 42:3;O2 | 1.805 | 0.85202 | 0.036742 | 0.085617 | 1.0674 |
| SM 42:3;O3 | 2.513 | 1.3294 | 0.0050625 | 0.02744 | 1.5616 |
| SM 42:4;O2 | 1.8411 | 0.88058 | 0.084761 | 0.1585 | 0.79996 |
| SM 43:1;O2_1 | 1.2625 | 0.33626 | 0.62087 | 0.68699 | 0.16305 |
| SM 43:1;O2_2 | 1.9619 | 0.97225 | 0.045875 | 0.10004 | 0.99982 |
| SM 43:2;O2_1 | 1.8672 | 0.90086 | 0.017844 | 0.056316 | 1.2494 |
| SM 43:2;O2_2 | 2.2533 | 1.172 | 0.02025 | 0.060588 | 1.2176 |
| SM 43:3;O2 | 2.553 | 1.3522 | 0.025885 | 0.068659 | 1.1633 |
| SM 44:1;O2 | 1.335 | 0.41685 | 0.40049 | 0.50348 | 0.29802 |
| SM 44:2;O2 | 1.5381 | 0.62117 | 0.25171 | 0.35325 | 0.45192 |
| SM 44:3;O2 | 1.5992 | 0.67734 | 0.3314 | 0.43413 | 0.36238 |
| SM 44:4;O2 | 2.472 | 1.3057 | 0.3314 | 0.43413 | 0.36238 |
| SM 44:5;O2 | 2.1279 | 1.0894 | 0.56129 | 0.64197 | 0.19249 |
| ST 27:1;O;S | 1.0104 | 0.014917 | 0.0079031 | 0.035398 | 1.451 |
| ST 28:1;O;S | 1.9691 | 0.97751 | 0.0068331 | 0.033626 | 1.4733 |
| Stearoyl-EA | 3.4456 | 1.7848 | 5.28E-05 | 0.0046651 | 2.3311 |
| TG 36:0 | 0.12757 | -2.9707 | 0.010482 | 0.041049 | 1.3867 |
| TG 38:0 | 0.14717 | -2.7644 | 0.10231 | 0.18134 | 0.7415 |
| TG 38:1 | 0.023906 | -5.3865 | 0.0019061 | 0.016202 | 1.7904 |
| TG 40:0 | 0.1213 | -3.0434 | 0.093216 | 0.16924 | 0.77151 |
| TG 40:1 | 0.064123 | -3.963 | 0.036742 | 0.085617 | 1.0674 |
| TG 40:2 | 0.062386 | -4.0026 | 0.010482 | 0.041049 | 1.3867 |
| TG 41:0 | 0.1101 | -3.1831 | 0.062953 | 0.12624 | 0.89879 |
| TG 41:1 | 0.04979 | -4.328 | 0.036742 | 0.085617 | 1.0674 |
| TG 41:2 | 0.033774 | -4.8879 | 0.010482 | 0.041049 | 1.3867 |
| TG 42:0 | 0.21744 | -2.2013 | 0.076919 | 0.14787 | 0.83012 |

| | | | | | |
|---|---|---|---|---|---|
| TG 43:0 | 0.17958 | -2.4773 | 0.056773 | 0.11635 | 0.93425 |
| TG 43:1 | 0.076732 | -3.704 | 0.025885 | 0.068659 | 1.1633 |
| TG 43:2 | 0.071964 | -3.7966 | 0.045875 | 0.10004 | 0.99982 |
| TG 44:0 | 0.47189 | -1.0835 | 0.2703 | 0.37166 | 0.42985 |
| TG 44:3 | 0.086468 | -3.5317 | 0.03277 | 0.080368 | 1.0949 |
| TG 45:0 | 0.26793 | -1.9001 | 0.036742 | 0.085617 | 1.0674 |
| TG 45:1 | 0.10488 | -3.2533 | 0.022923 | 0.063978 | 1.194 |
| TG 45:2 | 0.074386 | -3.7488 | 0.015684 | 0.052844 | 1.277 |
| TG 45:3 | 0.1007 | -3.3119 | 0.03277 | 0.080368 | 1.0949 |
| TG 46:0 | 0.7294 | -0.45523 | 0.42526 | 0.52928 | 0.27632 |
| TG 46:1 | 0.29635 | -1.7546 | 0.062953 | 0.12624 | 0.89879 |
| TG 47:0 | 0.43852 | -1.1893 | 0.062953 | 0.12624 | 0.89879 |
| TG 47:1 | 0.17856 | -2.4856 | 0.041102 | 0.093733 | 1.0281 |
| TG 47:2 | 0.10695 | -3.225 | 0.017844 | 0.056316 | 1.2494 |
| TG 47:3 | 0.098334 | -3.3462 | 0.029159 | 0.073935 | 1.1311 |
| TG 47:4 | 0.1176 | -3.0881 | 0.069659 | 0.13712 | 0.86291 |
| TG 48:6 | 0.099137 | -3.3344 | 0.069659 | 0.13712 | 0.86291 |
| TG 49:1 | 0.27644 | -1.8549 | 0.076919 | 0.14787 | 0.83012 |
| TG 49:3 | 0.1169 | -3.0967 | 0.015684 | 0.052844 | 1.277 |
| TG 49:4 | 0.15065 | -2.7307 | 0.036742 | 0.085617 | 1.0674 |
| TG 50:5 | 0.15537 | -2.6863 | 0.05109 | 0.10857 | 0.96431 |
| TG 50:6 | 0.24079 | -2.0541 | 0.14562 | 0.23526 | 0.62846 |
| TG 51:0 | 0.40525 | -1.3031 | 0.036742 | 0.085617 | 1.0674 |
| TG 51:1 | 0.30738 | -1.7019 | 0.12252 | 0.20877 | 0.68034 |
| TG 51:2 | 0.22462 | -2.1545 | 0.045875 | 0.10004 | 0.99982 |
| TG 51:3 | 0.18217 | -2.4566 | 0.03277 | 0.080368 | 1.0949 |
| TG 51:5 | 0.3244 | -1.6242 | 0.12252 | 0.20877 | 0.68034 |
| TG 51:6 | 0.26122 | -1.9367 | 0.10231 | 0.18134 | 0.7415 |
| TG 52:1 | 0.39179 | -1.3518 | 0.18608 | 0.28464 | 0.5457 |
| TG 52:7 | 0.12522 | -2.9975 | 0.084761 | 0.1585 | 0.79996 |
| TG 52:8 | 0.10813 | -3.2091 | 0.14562 | 0.23526 | 0.62846 |
| TG 53:1 | 0.17901 | -2.4819 | 0.03277 | 0.080368 | 1.0949 |
| TG 53:2 | 0.29859 | -1.7438 | 0.093216 | 0.16924 | 0.77151 |
| TG 53:3 | 0.19803 | -2.3362 | 0.025885 | 0.068659 | 1.1633 |
| TG 53:4 | 0.21051 | -2.248 | 0.03277 | 0.080368 | 1.0949 |
| TG 53:5 | 0.41598 | -1.2654 | 0.093216 | 0.16924 | 0.77151 |
| TG 53:6 | 0.25326 | -1.9813 | 0.093216 | 0.16924 | 0.77151 |
| TG 54:1 | 0.31468 | -1.6681 | 0.31014 | 0.41206 | 0.38504 |
| TG 54:4 | 0.26594 | -1.9108 | 0.036742 | 0.085617 | 1.0674 |
| TG 54:5 | 0.22347 | -2.1619 | 0.025885 | 0.068659 | 1.1633 |
| TG 54:6 | 0.24743 | -2.0149 | 0.10231 | 0.18134 | 0.7415 |
| TG 55:1 | 0.34641 | -1.5295 | 0.2703 | 0.37166 | 0.42985 |
| TG 55:2 | 0.094236 | -3.4076 | 0.015684 | 0.052844 | 1.277 |
| TG 55:3 | 0.13797 | -2.8576 | 0.029159 | 0.073935 | 1.1311 |

| | | | | | |
|---|---|---|---|---|---|
| TG 55:4 | 0.17627 | -2.5042 | 0.017844 | 0.056316 | 1.2494 |
| TG 55:6 | 0.41232 | -1.2782 | 0.15831 | 0.25088 | 0.60053 |
| TG 55:7 | 0.39859 | -1.327 | 0.18608 | 0.28464 | 0.5457 |
| TG 55:8 | 0.32115 | -1.6387 | 0.1337 | 0.22224 | 0.65318 |
| TG 56:1 | 0.3332 | -1.5855 | 0.21718 | 0.31666 | 0.4994 |
| TG 56:10 | 0.28901 | -1.7908 | 0.17179 | 0.26826 | 0.57144 |
| TG 56:2 | 0.15275 | -2.7108 | 0.084761 | 0.1585 | 0.79996 |
| TG 56:3 | 0.1426 | -2.81 | 0.05109 | 0.10857 | 0.96431 |
| TG 56:4 | 0.25136 | -1.9922 | 0.041102 | 0.093733 | 1.0281 |
| TG 56:7 | 0.42585 | -1.2316 | 0.21718 | 0.31666 | 0.4994 |
| TG 57:1 | 0.36262 | -1.4635 | 0.35355 | 0.45675 | 0.34032 |
| TG 57:2 | 0.2088 | -2.2598 | 0.21718 | 0.31666 | 0.4994 |
| TG 57:6 | 0.35708 | -1.4857 | 0.11207 | 0.1945 | 0.71109 |
| TG 57:7 | 0.36276 | -1.4629 | 0.18608 | 0.28464 | 0.5457 |
| TG 57:8 | 0.39077 | -1.3556 | 0.17179 | 0.26826 | 0.57144 |
| TG 58:1 | 0.40536 | -1.3027 | 0.31014 | 0.41206 | 0.38504 |
| TG 58:12 | 1.9173 | 0.93911 | 0.59074 | 0.66648 | 0.17621 |
| TG 58:2 | 0.15687 | -2.6724 | 0.056773 | 0.11635 | 0.93425 |
| TG 58:3 | 0.12115 | -3.0451 | 0.03277 | 0.080368 | 1.0949 |
| TG 58:4 | 0.14754 | -2.7609 | 0.017844 | 0.056316 | 1.2494 |
| TG 58:5 | 0.20126 | -2.3129 | 0.069659 | 0.13712 | 0.86291 |
| TG 58:7 | 0.1982 | -2.3349 | 0.056773 | 0.11635 | 0.93425 |
| TG 58:8_1 | 0.27625 | -1.856 | 0.2012 | 0.3004 | 0.5223 |
| TG 58:8_2 | 0.22231 | -2.1694 | 0.12252 | 0.20877 | 0.68034 |
| TG 58:9 | 0.24078 | -2.0542 | 0.15831 | 0.25088 | 0.60053 |
| TG 59:1 | 0.37907 | -1.3995 | 0.28978 | 0.38984 | 0.40911 |
| TG 59:2 | 0.21126 | -2.2429 | 0.2703 | 0.37166 | 0.42985 |
| TG 59:3 | 0.16965 | -2.5593 | 0.062953 | 0.12624 | 0.89879 |
| TG 59:4 | 0.15382 | -2.7006 | 0.0091146 | 0.038959 | 1.4094 |
| TG 60:1 | 0.421 | -1.2481 | 0.31014 | 0.41206 | 0.38504 |
| TG 60:10 | 0.52163 | -0.93889 | 0.25171 | 0.35325 | 0.45192 |
| TG 60:11 | 0.57959 | -0.78689 | 0.62087 | 0.68699 | 0.16305 |
| TG 60:13 | 0.74981 | -0.41541 | 0.69444 | 0.75281 | 0.12331 |
| TG 60:2 | 0.21935 | -2.1887 | 0.23401 | 0.33468 | 0.47537 |
| TG 60:3 | 0.086686 | -3.5281 | 0.025885 | 0.068659 | 1.1633 |
| TG 60:4 | 0.10314 | -3.2773 | 0.015684 | 0.052844 | 1.277 |
| TG 60:6 | 0.21754 | -2.2007 | 0.025885 | 0.068659 | 1.1633 |
| TG 60:7 | 0.28722 | -1.7998 | 0.02025 | 0.060588 | 1.2176 |
| TG 61:1 | 0.37165 | -1.428 | 0.35355 | 0.45675 | 0.34032 |
| TG 61:2 | 0.19504 | -2.3582 | 0.21718 | 0.31666 | 0.4994 |
| TG 61:3 | 0.13742 | -2.8634 | 0.056773 | 0.11635 | 0.93425 |
| TG 62:1 | 0.51511 | -0.95705 | 0.28978 | 0.38984 | 0.40911 |
| TG 62:10 | 0.60911 | -0.71522 | 0.47731 | 0.57771 | 0.23829 |
| TG 62:11 | 0.51899 | -0.94623 | 0.62087 | 0.68699 | 0.16305 |

| | | | | | |
|---|---|---|---|---|---|
| TG 62:12 | 0.40225 | -1.3138 | 0.28978 | 0.38984 | 0.40911 |
| TG 62:13 | 0.79868 | -0.32431 | 0.53255 | 0.61759 | 0.2093 |
| TG 62:14 | 0.65003 | -0.62142 | 0.50454 | 0.59904 | 0.22254 |
| TG 62:2 | 0.24823 | -2.0102 | 0.2012 | 0.3004 | 0.5223 |
| TG 62:3 | 0.24139 | -2.0506 | 0.23401 | 0.33468 | 0.47537 |
| TG 62:4 | 0.16349 | -2.6127 | 0.045875 | 0.10004 | 0.99982 |
| TG 62:5 | 0.18524 | -2.4325 | 0.045875 | 0.10004 | 0.99982 |
| TG 62:6 | 0.15821 | -2.6601 | 0.010482 | 0.041049 | 1.3867 |
| TG 63:2 | 0.26129 | -1.9363 | 0.3314 | 0.43413 | 0.36238 |
| TG 64:1 | 0.581 | -0.78339 | 0.59074 | 0.66648 | 0.17621 |
| TG 64:2 | 0.35328 | -1.5011 | 0.2703 | 0.37166 | 0.42985 |
| TG 64:3 | 0.36677 | -1.4471 | 0.37659 | 0.47988 | 0.31887 |
| TG 64:6 | 0.27408 | -1.8673 | 0.084761 | 0.1585 | 0.79996 |
| TG 66:1 | 0.56034 | -0.83562 | 0.55569 | 0.64197 | 0.19249 |
| TG O-52:0 | 0.98692 | -0.018993 | 0.77999 | 0.82523 | 0.083426 |
| TG O-54:1 | 0.2783 | -1.8453 | 0.62087 | 0.68699 | 0.16305 |
| TG O-54:2 | 0.23453 | -2.0922 | 0.2012 | 0.3004 | 0.5223 |
| TG O-58:3 | 0.13923 | -2.8444 | 0.40049 | 0.50348 | 0.29802 |

**Supplementary Table 2:** Plasma Lipidome – Fold Change CAVD patients vs. Controls

| Lipid | Fold Change (FC) | log2(FC) | raw.pval | -log10(p) |
|---|---|---|---|---|
| CAR 10:0 | 1.5327 | 0.61607 | 0.040689 | 1.3905 |
| CAR 11:0 | 1.1076 | 0.14741 | 0.70451 | 0.15211 |
| CAR 11:1 | 1.2753 | 0.35081 | 0.43189 | 0.36462 |
| CAR 12:0 | 1.5763 | 0.65658 | 0.052643 | 1.2787 |
| CAR 12:1 | 1.4393 | 0.52539 | 0.059596 | 1.2248 |
| CAR 13:0 | 1.7378 | 0.79724 | 0.27398 | 0.56228 |
| CAR 13:1 | 1.313 | 0.39283 | 0.3744 | 0.42667 |
| CAR 14:0 | 1.2671 | 0.34148 | 0.046354 | 1.3339 |
| CAR 14:1 | 1.7089 | 0.77304 | 0.040689 | 1.3905 |
| CAR 14:2 | 1.7054 | 0.77009 | 106 | 0.97469 |
| CAR 15:1 | 1.2589 | 0.33214 | 0.1598 | 0.79643 |
| CAR 16:0 | 1.114 | 0.15572 | 0.43189 | 0.36462 |
| CAR 16:1 | 1.4555 | 0.54151 | 0.017335 | 1.7611 |
| CAR 16:3 | 1.7092 | 0.77332 | 0.010767 | 1.9679 |
| CAR 16:4 | 1.3082 | 0.38755 | 0.1598 | 0.79643 |
| CAR 18:3 | 1.4858 | 0.57128 | 0.040689 | 1.3905 |
| CAR 20:1 | 1.7112 | 0.77502 | 0.046354 | 1.3339 |
| CAR 20:4 | 1.4405 | 0.52656 | 0.13084 | 0.88324 |
| CAR 20:5 | 1.1513 | 0.20322 | 0.78101 | 0.10734 |
| CAR 22:0 | 1.0609 | 0.085319 | 0.97984 | 0.0088434 |
| CAR 22:1 | 1.504 | 0.58879 | 106 | 0.97469 |
| CAR 22:6 | 1.3654 | 0.44931 | 0.52671 | 0.27843 |
| CAR 24:0 | 0.94448 | -0.082413 | 0.93959 | 0.027062 |
| CAR 24:1 | 1.2047 | 0.2687 | 0.32173 | 0.49251 |
| CAR 26:0 | 1.1085 | 0.1486 | 0.52671 | 0.27843 |
| CAR 26:1 | 1.2495 | 0.32131 | 0.29723 | 0.5269 |
| CAR 8:0 | 1.6432 | 0.71649 | 0.035598 | 1.4486 |
| CAR 8:1 | 1.3019 | 0.38065 | 0.11793 | 0.92839 |
| CE 16:1 | 1.0253 | 0.0361 | 0.74245 | 0.12933 |
| CE 17:0 | 0.7286 | -0.45679 | 0.1759 | 0.75473 |
| CE 18:3 | 1.041 | 0.057924 | 1 | 0 |
| CE 20:2 | 1.0917 | 0.12657 | 0.52671 | 0.27843 |
| CE 20:4 | 1.0277 | 0.039474 | 1 | 0 |
| CE 20:5 | 1.0961 | 0.13243 | 0.93959 | 0.027062 |
| CE 22:4 | 1.2166 | 0.28291 | 0.27398 | 0.56228 |
| CE 22:5 | 1.0409 | 0.057891 | 0.66725 | 0.17571 |
| CE 22:6 | 0.9586 | -0.060995 | 0.93959 | 0.027062 |
| Cer 16:0;O2/24:1 | 0.72648 | -0.46101 | 0.3744 | 0.42667 |
| Cer 16:1;O2/24:1 | 0.9464 | -0.079481 | 0.93959 | 0.027062 |
| Cer 18:0;O2/18:0 | 0.98112 | -0.027496 | 1 | 0 |
| Cer 18:0;O2/20:0 | 0.93576 | -0.095786 | 0.93959 | 0.027062 |

| | | | | |
|---|---|---|---|---|
| Cer 18:0;O2/22:0 | 1.0591 | 0.082812 | 0.63079 | 0.20012 |
| Cer 18:0;O2/23:0 | 1.0156 | 0.022286 | 0.74245 | 0.12933 |
| Cer 18:0;O2/24:0 | 1.063 | 0.08818 | 0.49401 | 0.30626 |
| Cer 18:0;O2/24:2 | 1.3286 | 0.40994 | 0.29723 | 0.5269 |
| Cer 18:0;O3/22:0 | 0.82359 | -0.27999 | 0.095012 | 1.0222 |
| Cer 18:0;O3/23:0 | 0.89321 | -0.16293 | 0.3744 | 0.42667 |
| Cer 18:0;O3/24:0 | 0.79354 | -0.33362 | 0.023364 | 1.6315 |
| Cer 18:0;O3/24:1 | 0.86347 | -0.21178 | 0.29723 | 0.5269 |
| Cer 18:1;O2/16:0 | 1.027 | 0.038426 | 0.82009 | 0.086138 |
| Cer 18:1;O2/18:0 | 0.99057 | -0.013672 | 0.97984 | 0.0088434 |
| Cer 18:1;O2/20:0 | 0.97981 | -0.029425 | 0.74245 | 0.12933 |
| Cer 18:1;O2/22:0;(2OH) | 1.104 | 0.14273 | 0.59516 | 0.22537 |
| Cer 18:1;O2/23:0 | 1.0429 | 0.060596 | 0.97984 | 0.0088434 |
| Cer 18:1;O2/24:0 | 0.92076 | -0.1191 | 0.66725 | 0.17571 |
| Cer 18:1;O2/24:0;(2OH) | 1.1108 | 0.15154 | 0.52671 | 0.27843 |
| Cer 18:1;O2/24:1 | 1.0483 | 0.068112 | 0.74245 | 0.12933 |
| Cer 18:1;O2/24:1;(2OH) | 1.2765 | 0.35221 | 0.21156 | 0.67458 |
| Cer 18:1;O2/24:2 | 1.0013 | 0.0018129 | 0.66725 | 0.17571 |
| Cer 18:1;O2/25:0 | 0.99458 | -0.0078391 | 0.70451 | 0.15211 |
| Cer 18:1;O2/26:1 | 1.2958 | 0.37382 | 0.43189 | 0.36462 |
| Cer 18:1;O3/24:0 | 1.0827 | 0.11465 | 0.49401 | 0.30626 |
| Cer 18:2;O2/20:0 | 0.94807 | -0.076933 | 0.46239 | 0.33499 |
| Cer 18:2;O2/21:2 | 0.99575 | -0.0061489 | 0.78101 | 0.10734 |
| Cer 18:2;O2/22:0 | 1.1177 | 0.16053 | 0.40255 | 0.39518 |
| Cer 18:2;O2/23:0 | 1.0999 | 0.13734 | 0.52671 | 0.27843 |
| Cer 18:2;O2/24:0 | 0.98294 | -0.024829 | 0.82009 | 0.086138 |
| Cer 18:2;O2/24:0;(2OH) | 1.2508 | 0.32284 | 0.25195 | 0.59868 |
| Cer 18:2;O2/24:1 | 1.1158 | 0.15804 | 0.21156 | 0.67458 |
| Cer 18:2;O2/25:0 | 0.98718 | -0.018619 | 0.52671 | 0.27843 |
| Cer 19:1;O2/24:0 | 0.84648 | -0.24045 | 0.82009 | 0.086138 |
| Cer 19:1;O2/24:1 | 0.82948 | -0.26972 | 0.49401 | 0.30626 |
| Cer 20:0;O2/24:0 | 1.0234 | 0.033384 | 0.97984 | 0.0088434 |
| Cer 39:1;O2 | 0.93452 | -0.097698 | 0.70451 | 0.15211 |
| CoQ10 | 815 | -0.29512 | 0.82009 | 0.086138 |
| CoQ9 | 0.77833 | -0.36155 | 0.97984 | 0.0088434 |
| DG 16:0_16:0 | 0.58658 | -0.76961 | 0.59516 | 0.22537 |
| DG 16:0_16:1 | 0.39807 | -1.3289 | 0.63079 | 0.20012 |
| DG 16:0_18:0 | 0.56812 | -0.81574 | 0.63079 | 0.20012 |
| DG 16:0_18:1 | 0.66511 | -0.58834 | 0.66725 | 0.17571 |
| DG 16:0_18:2 | 0.52496 | -0.92971 | 1 | 0 |
| DG 16:0_20:4 | 0.75993 | -0.39605 | 0.93959 | 0.027062 |
| DG 16:0_22:6 | 0.5252 | -0.92905 | 0.70451 | 0.15211 |
| DG 16:1_18:1 | 0.41004 | -1.2862 | 0.63079 | 0.20012 |
| DG 17:1_18:1 | 0.50436 | -0.98748 | 0.89948 | 0.046011 |

| | | | | |
|---|---|---|---|---|
| DG 18:0_18:1 | 0.56598 | -0.82117 | 0.63079 | 0.20012 |
| DG 18:0_18:2 | 0.65588 | -0.6085 | 0.70451 | 0.15211 |
| DG 18:0_18:3 | 0.65086 | -0.61958 | 0.66725 | 0.17571 |
| DG 18:0_20:4 | 0.67592 | -0.56507 | 0.66725 | 0.17571 |
| DG 18:1_18:1 | 0.59905 | -0.73924 | 0.74245 | 0.12933 |
| DG 18:1_18:2 | 0.58244 | -0.77981 | 0.82009 | 0.086138 |
| DG 18:1_20:1 | 0.69286 | -0.52937 | 0.78101 | 0.10734 |
| DG 18:1_20:3 | 0.54887 | -0.86547 | 0.74245 | 0.12933 |
| DG 18:1_20:4 | 0.68732 | -0.54094 | 0.82009 | 0.086138 |
| DG 18:1_22:4 | 0.74303 | -0.42851 | 0.97984 | 0.0088434 |
| DG 18:1_22:5 | 0.47945 | -1.0605 | 0.49401 | 0.30626 |
| DG 18:2_18:3 | 0.73053 | -0.45299 | 0.78101 | 0.10734 |
| DG 18:2_22:6 | 727 | -0.45997 | 0.52671 | 0.27843 |
| DG 36:4 | 0.72976 | -0.4545 | 0.82009 | 0.086138 |
| FA 16:0-2OH | 1.0873 | 0.12074 | 0.89948 | 0.046011 |
| FA 18:0 | 1.428 | 0.51396 | 0.1759 | 0.75473 |
| FA 18:1 | 1.6939 | 0.76031 | 0.27398 | 0.56228 |
| FA 18:2 | 1.7662 | 0.82068 | 0.59516 | 0.22537 |
| FA 20:1 | 2.001 | 1.0008 | 0.052643 | 1.2787 |
| FA 20:4 | 1.9934 | 0.9952 | 0.059596 | 1.2248 |
| FA 22:5 | 2.9549 | 1.5631 | 0.1598 | 0.79643 |
| FA 22:6 | 1.598 | 0.67622 | 0.43189 | 0.36462 |
| GM3 32:1;2O | 0.83791 | -0.25513 | 0.1598 | 0.79643 |
| GM3 33:1;2O | 0.90734 | -0.14029 | 0.63079 | 0.20012 |
| GM3 34:1;2O | 1.006 | 0.0085799 | 0.89948 | 0.046011 |
| GM3 34:2;2O | 0.99252 | -0.010828 | 0.89948 | 0.046011 |
| GM3 35:1;2O | 0.83851 | -0.2541 | 0.32173 | 0.49251 |
| GM3 36:1;2O | 0.99163 | -0.012123 | 0.89948 | 0.046011 |
| GM3 36:2;2O | 0.99519 | -0.0069611 | 0.66725 | 0.17571 |
| GM3 38:1;2O | 1.0064 | 0.0092072 | 0.70451 | 0.15211 |
| GM3 40:2;2O | 0.99945 | -0.0007958 | 0.52671 | 0.27843 |
| GM3 42:3;2O | 1.0665 | 0.092841 | 0.74245 | 0.12933 |
| Hex2Cer 18:1;O2/16:0 | 0.9974 | -0.0037488 | 0.97984 | 0.0088434 |
| Hex2Cer 18:1;O2/22:0 | 0.97047 | -0.043241 | 0.66725 | 0.17571 |
| Hex2Cer 18:1;O2/23:0 | 1.0357 | 0.050627 | 0.74245 | 0.12933 |
| Hex2Cer 18:1;O2/24:0 | 0.97986 | -0.029353 | 0.97984 | 0.0088434 |
| Hex2Cer 18:1;O2/24:1 | 0.9128 | -0.13164 | 0.66725 | 0.17571 |
| Hex3Cer 18:1;O2/14:0 | 0.93093 | -0.10325 | 0.56045 | 0.25146 |
| Hex3Cer 18:1;O2/16:0 | 0.89879 | -0.15395 | 0.34745 | 0.45911 |
| Hex3Cer 18:1;O2/22:0 | 0.88733 | -0.17246 | 0.32173 | 0.49251 |
| Hex3Cer 18:1;O2/24:0 | 0.92936 | -0.1057 | 0.46239 | 0.33499 |
| Hex3Cer 18:1;O2/24:1 | 1.0736 | 0.10245 | 0.74245 | 0.12933 |
| Hex3Cer 18:2;O2/16:0 | 1.1124 | 0.15372 | 0.59516 | 0.22537 |
| Hex3Cer 18:2;O2/24:1 | 1.1268 | 0.17217 | 0.52671 | 0.27843 |

| | | | | |
|---|---|---|---|---|
| HexCer 18:1;O2/20:0 | 1.0338 | 0.048003 | 0.59516 | 0.22537 |
| HexCer 18:1;O2/22:0 | 1.0149 | 0.021376 | 0.85962 | 0.065695 |
| HexCer 18:1;O2/23:0 | 0.92058 | -0.11939 | 0.89948 | 0.046011 |
| HexCer 18:1;O2/24:0 | 0.86847 | -0.20345 | 0.25195 | 0.59868 |
| HexCer 18:1;O2/24:1 | 1.0587 | 0.082257 | 0.63079 | 0.20012 |
| HexCer 18:2;O2/22:0 | 1.0927 | 0.12791 | 0.89948 | 0.046011 |
| HexCer 18:2;O2/23:0 | 1.0162 | 0.023233 | 1 | 0 |
| HexCer 18:2;O2/24:1 | 1.0918 | 0.12665 | 0.70451 | 0.15211 |
| LNAPE 16:0/N-18:0 | 1.0973 | 134 | 0.34745 | 0.45911 |
| LNAPE 18:1/N-21:0 | 1.0378 | 0.053594 | 0.66725 | 0.17571 |
| LNAPE 18:2/N-2:0 | 1.3037 | 0.38256 | 0.32173 | 0.49251 |
| LNAPE 18:3/N-18:0 | 0.98782 | -0.017679 | 0.85962 | 0.065695 |
| LNAPE 20:3/N-18:0 | 1.1539 | 0.20651 | 0.34745 | 0.45911 |
| LNAPE 20:3/N-20:0 | 1.0285 | 0.0406 | 0.82009 | 0.086138 |
| LNAPE 20:4/N-16:0 | 1.234 | 0.30332 | 0.46239 | 0.33499 |
| LNAPE 20:4/N-17:0 | 0.93984 | -0.089511 | 1 | 0 |
| LNAPE 20:4/N-18:1 | 0.90965 | -0.13661 | 0.89948 | 0.046011 |
| LNAPE 20:4/N-2:0 | 1.1232 | 0.16763 | 0.74245 | 0.12933 |
| LNAPE 32:0 | 0.96356 | -0.053547 | 0.89948 | 0.046011 |
| LNAPE 34:2 | 0.8465 | -0.24042 | 0.27398 | 0.56228 |
| LNAPE 35:2 | 0.84405 | -0.24461 | 0.19315 | 0.7141 |
| LNAPE 36:0 | 1.0573 | 0.080438 | 0.78101 | 0.10734 |
| LNAPE 38:1 | 1.0303 | 0.043035 | 0.93959 | 0.027062 |
| LNAPE 38:5 | 1.005 | 0.0071658 | 0.97984 | 0.0088434 |
| LNAPE 40:6 | 0.98041 | -0.028547 | 0.93959 | 0.027062 |
| LNAPE 42:4 | 1.3754 | 0.45984 | 0.29723 | 0.5269 |
| LNAPE 42:6 | 1.039 | 0.055216 | 0.74245 | 0.12933 |
| LPC 12:0 | 0.99269 | -0.010589 | 0.89948 | 0.046011 |
| LPC 14:0 | 0.86224 | -0.21384 | 0.56045 | 0.25146 |
| LPC 14:1 | 0.79929 | -0.3232 | 0.49401 | 0.30626 |
| LPC 15:0 | 0.84143 | -0.24908 | 0.27398 | 0.56228 |
| LPC 16:0 | 0.94621 | -0.079762 | 0.70451 | 0.15211 |
| LPC 16:1 | 0.88042 | -0.18373 | 0.89948 | 0.046011 |
| LPC 17:0 | 1.1772 | 0.23533 | 0.70451 | 0.15211 |
| LPC 17:1 | 0.87321 | -0.19559 | 0.59516 | 0.22537 |
| LPC 17:2 | 0.97906 | -0.030529 | 0.93959 | 0.027062 |
| LPC 18:0 | 0.88534 | -0.17569 | 0.40255 | 0.39518 |
| LPC 18:1 | 0.96572 | -0.050328 | 0.85962 | 0.065695 |
| LPC 18:2 | 0.90863 | -0.13823 | 0.49401 | 0.30626 |
| LPC 18:3 | 1.0175 | 0.025051 | 0.93959 | 0.027062 |
| LPC 19:1 | 0.93163 | -0.10217 | 0.59516 | 0.22537 |
| LPC 20:1 | 0.9858 | -0.020634 | 0.85962 | 0.065695 |
| LPC 20:2 | 0.86195 | -0.21433 | 0.59516 | 0.22537 |
| LPC 20:3 | 0.92137 | -0.11815 | 0.52671 | 0.27843 |

| | | | | |
|---|---|---|---|---|
| LPC 20:4 | 1.0131 | 0.01877 | 0.93959 | 0.027062 |
| LPC 20:5 | 1.0187 | 0.026769 | 0.3744 | 0.42667 |
| LPC 22:0 | 0.87175 | -0.19801 | 0.40255 | 0.39518 |
| LPC 22:3 | 1.1789 | 0.2374 | 0.85962 | 0.065695 |
| LPC 22:4 | 1.1609 | 0.21519 | 0.49401 | 0.30626 |
| LPC 22:5 | 1.012 | 0.017196 | 0.40255 | 0.39518 |
| LPC 22:6 | 0.97534 | -0.03603 | 0.3744 | 0.42667 |
| LPC 23:0 | 0.90274 | -0.14761 | 0.43189 | 0.36462 |
| LPC 24:0 | 0.87749 | -0.18855 | 0.32173 | 0.49251 |
| LPC 24:1 | 1.2003 | 0.26336 | 0.56045 | 0.25146 |
| LPC 24:5 | 0.86236 | -0.21364 | 0.43189 | 0.36462 |
| LPC O-16:1 | 0.86314 | -0.21233 | 0.34745 | 0.45911 |
| LPC O-17:0 | 0.72047 | -0.47298 | 0.11793 | 0.92839 |
| LPC O-18:1 | 1.0703 | 0.09802 | 0.85962 | 0.065695 |
| LPC O-20:0 | 0.87029 | -0.20043 | 0.3744 | 0.42667 |
| LPC O-22:0 | 0.9295 | -0.10547 | 0.29723 | 0.5269 |
| LPC O-22:1 | 0.85103 | -0.23271 | 0.23115 | 0.6361 |
| LPC O-24:0 | 0.95537 | -0.065867 | 0.25195 | 0.59868 |
| LPC O-24:1 | 1.0407 | 0.057491 | 1 | 0 |
| LPC O-24:2 | 0.68318 | -0.54966 | 0.1598 | 0.79643 |
| LPC O-26:1 | 0.89018 | -0.16783 | 0.49401 | 0.30626 |
| LPE 16:0 | 0.94813 | -0.07685 | 0.89948 | 0.046011 |
| LPE 18:0 | 1.0644 | 0.09 | 0.49401 | 0.30626 |
| LPE 18:2 | 1.08 | 0.11101 | 0.52671 | 0.27843 |
| LPE 20:3 | 1.104 | 0.14274 | 0.89948 | 0.046011 |
| LPE 20:4 | 1.243 | 0.31384 | 0.23115 | 0.6361 |
| LPE 20:5 | 0.90171 | -0.14927 | 0.56045 | 0.25146 |
| LPE 22:6 | 1.0537 | 0.075465 | 0.93959 | 0.027062 |
| LPE O-16:1 | 0.9766 | -0.034164 | 1 | 0 |
| LPE O-18:1 | 1.0429 | 0.060621 | 0.89948 | 0.046011 |
| LPE O-18:2 | 1.0044 | 0.0063625 | 0.97984 | 0.0088434 |
| LPE O-22:1 | 0.93863 | -0.091365 | 0.43189 | 0.36462 |
| LPG 18:1 | 1.2028 | 0.26635 | 0.084927 | 1.071 |
| LPI 16:0 | 0.92356 | -0.11473 | 0.70451 | 0.15211 |
| LPI 16:1 | 1.096 | 0.1323 | 0.78101 | 0.10734 |
| LPI 18:0 | 0.9647 | -0.051851 | 1 | 0 |
| LPI 18:1 | 1.1623 | 0.21702 | 0.46239 | 0.33499 |
| LPI 18:2 | 1.2842 | 0.3609 | 0.49401 | 0.30626 |
| LPI 20:3 | 1.0261 | 0.037222 | 0.82009 | 0.086138 |
| LPI 20:4 | 1.3177 | 398 | 0.70451 | 0.15211 |
| LPI 22:5 | 1.2205 | 0.28753 | 0.49401 | 0.30626 |
| LPI 22:6 | 1.1881 | 0.24861 | 0.70451 | 0.15211 |
| PC 14:0_20:4 | 0.83803 | -0.25493 | 0.97984 | 0.0088434 |
| PC 15:0_16:0 | 0.91496 | -0.12822 | 0.52671 | 0.27843 |

| | | | | |
|---|---|---|---|---|
| PC 15:0_18:0 | 0.84683 | -0.23986 | 0.52671 | 0.27843 |
| PC 16:0_17:0 | 0.82474 | -0.27798 | 0.40255 | 0.39518 |
| PC 16:0_18:1 | 0.97781 | -0.032379 | 0.93959 | 0.027062 |
| PC 16:0_18:2 | 0.97264 | -0.040025 | 0.43189 | 0.36462 |
| PC 16:0_20:4 | 1.0186 | 0.026578 | 0.93959 | 0.027062 |
| PC 16:1_18:2 | 0.92875 | -0.10664 | 0.56045 | 0.25146 |
| PC 16:1_18:3 | 0.79805 | -0.32545 | 0.70451 | 0.15211 |
| PC 16:1_20:5 | 0.66546 | -0.58757 | 0.32173 | 0.49251 |
| PC 17:0_20:3 | 0.86222 | -0.21387 | 0.21156 | 0.67458 |
| PC 17:1_18:2 | 0.96612 | -0.049732 | 0.97984 | 0.0088434 |
| PC 18:0_18:1 | 0.9676 | -0.047516 | 0.89948 | 0.046011 |
| PC 18:0_18:2 | 0.9523 | -0.070516 | 0.43189 | 0.36462 |
| PC 18:0_20:3 | 0.93273 | -0.10047 | 0.59516 | 0.22537 |
| PC 18:0_20:4 | 0.98455 | -0.022465 | 0.97984 | 0.0088434 |
| PC 18:0_20:5 | 1.0391 | 0.055312 | 0.63079 | 0.20012 |
| PC 18:0_22:6 | 0.93872 | -0.091231 | 0.63079 | 0.20012 |
| PC 18:1_18:2 | 0.99111 | -0.012885 | 0.56045 | 0.25146 |
| PC 18:1_20:1 | 1.13 | 0.17632 | 0.49401 | 0.30626 |
| PC 18:1_20:3 | 0.88736 | -0.17241 | 0.3744 | 0.42667 |
| PC 18:1_22:1 | 1.339 | 0.4212 | 0.25195 | 0.59868 |
| PC 18:1_22:6 | 0.86957 | -0.20163 | 0.19315 | 0.7141 |
| PC 18:2_18:2 | 1.0764 | 0.10618 | 0.63079 | 0.20012 |
| PC 18:2_20:4 | 0.90549 | -0.14322 | 0.46239 | 0.33499 |
| PC 18:2_22:5 | 0.93258 | -0.10069 | 0.49401 | 0.30626 |
| PC 19:0_20:4 | 0.91299 | -0.13133 | 0.59516 | 0.22537 |
| PC 20:3_20:4 | 0.94455 | -0.082301 | 0.85962 | 0.065695 |
| PC 20:4_22:4 | 0.97576 | -0.035403 | 0.66725 | 0.17571 |
| PC 20:4_22:5 | 0.79626 | -0.32869 | 0.25195 | 0.59868 |
| PC 20:4_22:6 | 0.75858 | -0.39863 | 0.19315 | 0.7141 |
| PC 20:4_24:1 | 1.203 | 0.26659 | 0.059596 | 1.2248 |
| PC 22:0_18:2 | 0.93864 | -0.091358 | 0.32173 | 0.49251 |
| PC 22:0_20:4 | 1.0044 | 0.0063406 | 0.89948 | 0.046011 |
| PC 22:1_18:2 | 1.471 | 0.55675 | 0.19315 | 0.7141 |
| PC 22:1_20:4 | 1.3512 | 0.43424 | 0.075691 | 1.121 |
| PC 24:0_18:2 | 0.87906 | -0.18596 | 0.70451 | 0.15211 |
| PC 26:0 | 1.2392 | 0.30939 | 0.93959 | 0.027062 |
| PC 26:2 | 1.8083 | 0.85463 | 0.68067 | 0.16706 |
| PC 28:0 | 0.87151 | -0.19841 | 0.59516 | 0.22537 |
| PC 29:0 | 0.8256 | -0.27648 | 0.40255 | 0.39518 |
| PC 30:0 | 1.0124 | 0.01772 | 0.59516 | 0.22537 |
| PC 30:1 | 0.90459 | -0.14467 | 0.82009 | 0.086138 |
| PC 30:2 | 1.1255 | 0.1706 | 0.78101 | 0.10734 |
| PC 31:1 | 0.84113 | -0.2496 | 0.66725 | 0.17571 |
| PC 32:0 | 0.96334 | -0.053883 | 0.93959 | 0.027062 |

| | | | | |
|---|---|---|---|---|
| PC 32:1 | 1.002 | 0.0029041 | 0.93959 | 0.027062 |
| PC 32:2 | 0.8093 | -0.30525 | 0.46239 | 0.33499 |
| PC 33:1 | 0.92147 | -118 | 0.52671 | 0.27843 |
| PC 33:2 | 0.84403 | -0.24464 | 0.14478 | 0.83928 |
| PC 33:3 | 0.80372 | -0.31523 | 0.56045 | 0.25146 |
| PC 34:0 | 0.93202 | -0.10157 | 0.66725 | 0.17571 |
| PC 34:3 | 0.97241 | -0.040359 | 1 | 0 |
| PC 34:5 | 0.57166 | -0.80676 | 0.32173 | 0.49251 |
| PC 35:0 | 0.83116 | -0.26681 | 0.70451 | 0.15211 |
| PC 35:1 | 0.92871 | -0.1067 | 0.59516 | 0.22537 |
| PC 35:2 | 0.8116 | -0.30116 | 0.25195 | 0.59868 |
| PC 35:3 | 0.90428 | -0.14515 | 0.43189 | 0.36462 |
| PC 35:4 | 0.83464 | -0.26077 | 0.63079 | 0.20012 |
| PC 35:5 | 0.67098 | -0.57566 | 0.25195 | 0.59868 |
| PC 36:0 | 0.97817 | -0.031839 | 0.66725 | 0.17571 |
| PC 36:2 | 1.0174 | 0.024908 | 0.97984 | 0.0088434 |
| PC 36:3 | 0.92377 | -0.1144 | 0.43189 | 0.36462 |
| PC 36:5 | 0.99976 | -0.0003435 | 0.89948 | 0.046011 |
| PC 37:1 | 0.93329 | -0.099604 | 0.89948 | 0.046011 |
| PC 37:3 | 0.96456 | -0.052059 | 0.85962 | 0.065695 |
| PC 37:4 | 0.89664 | -0.1574 | 0.85962 | 0.065695 |
| PC 37:6 | 0.73773 | -0.43884 | 0.27398 | 0.56228 |
| PC 38:1 | 1.2099 | 0.27484 | 0.14478 | 0.83928 |
| PC 38:2 | 0.89659 | -0.15748 | 0.52671 | 0.27843 |
| PC 38:3 | 1.1092 | 0.14957 | 0.85962 | 0.065695 |
| PC 38:6 | 0.87133 | -0.19871 | 0.19315 | 0.7141 |
| PC 38:7 | 0.73689 | -0.44047 | 0.27398 | 0.56228 |
| PC 39:3 | 0.92427 | -0.11361 | 0.56045 | 0.25146 |
| PC 39:5 | 0.96212 | -0.055712 | 0.89948 | 0.046011 |
| PC 39:6 | 0.94663 | -0.079124 | 0.56045 | 0.25146 |
| PC 40:0 | 0.91352 | -0.13049 | 0.70451 | 0.15211 |
| PC 40:1 | 1.3772 | 0.46173 | 0.11793 | 0.92839 |
| PC 40:3 | 0.98093 | -0.027783 | 1 | 0 |
| PC 40:4 | 1.1679 | 0.22396 | 0.49401 | 0.30626 |
| PC 40:5 | 0.99985 | -0.0002114 | 1 | 0 |
| PC 40:8 | 0.82325 | -0.2806 | 0.25195 | 0.59868 |
| PC 40:9 | 0.95102 | -0.072457 | 0.97984 | 0.0088434 |
| PC 41:6 | 0.84387 | -0.2449 | 0.21156 | 0.67458 |
| PC 41:7 | 0.89148 | -0.16573 | 0.40255 | 0.39518 |
| PC 42:1 | 0.93591 | -0.095565 | 0.97984 | 0.0088434 |
| PC 42:11 | 0.74087 | -0.4327 | 0.23115 | 0.6361 |
| PC 42:2 | 1.2158 | 0.28192 | 0.27398 | 0.56228 |
| PC 42:3 | 1.1339 | 0.18133 | 0.32173 | 0.49251 |
| PC 42:4 | 1.2383 | 0.30836 | 0.13084 | 0.88324 |

| | | | | |
|---|---|---|---|---|
| PC 42:6 | 1.0355 | 0.050304 | 0.49401 | 0.30626 |
| PC 43:9 | 0.65279 | -0.61532 | 0.66725 | 0.17571 |
| PC 44:10 | 0.94451 | -0.082355 | 0.85962 | 0.065695 |
| PC 44:11 | 0.76777 | -0.38125 | 0.40255 | 0.39518 |
| PC 44:12 | 0.74562 | -0.4235 | 0.21156 | 0.67458 |
| PC 44:7 | 1.8043 | 0.85144 | 0.70451 | 0.15211 |
| PC O-16:0_20:3 | 0.81886 | -0.28832 | 0.059596 | 1.2248 |
| PC O-16:0_20:5 | 0.74059 | -0.43326 | 0.46239 | 0.33499 |
| PC O-16:0_22:6 | 0.95419 | -0.067647 | 0.43189 | 0.36462 |
| PC O-16:1_20:5 | 1.0065 | 0.0092764 | 0.46239 | 0.33499 |
| PC O-16:1_22:6 | 0.83318 | -0.2633 | 0.32173 | 0.49251 |
| PC O-18:0_16:0 | 0.93074 | -0.10354 | 0.46239 | 0.33499 |
| PC O-18:0_18:2 | 0.90316 | -0.14694 | 0.23115 | 0.6361 |
| PC O-18:0_20:4 | 0.98945 | -0.015305 | 0.93959 | 0.027062 |
| PC O-18:1_18:1 | 0.95527 | -0.06602 | 0.46239 | 0.33499 |
| PC O-18:1_18:2 | 0.95515 | -0.066197 | 0.70451 | 0.15211 |
| PC O-18:1_22:4 | 1.1179 | 0.16076 | 0.59516 | 0.22537 |
| PC O-18:1_22:5 | 0.92642 | -0.11026 | 0.63079 | 0.20012 |
| PC O-18:2_18:1 | 0.95336 | -0.068904 | 0.66725 | 0.17571 |
| PC O-20:0_20:4 | 0.96251 | -0.055133 | 0.78101 | 0.10734 |
| PC O-20:4_24:1 | 1.0064 | 0.0091683 | 0.89948 | 0.046011 |
| PC O-22:0_20:4 | 0.96052 | -0.058106 | 0.89948 | 0.046011 |
| PC O-22:1_20:3 | 1.084 | 0.11642 | 0.32173 | 0.49251 |
| PC O-22:1_22:6 | 0.91423 | -0.12937 | 0.49401 | 0.30626 |
| PC O-22:2_20:4 | 892 | -0.16488 | 0.89948 | 0.046011 |
| PC O-24:0_20:4 | 1.0094 | 0.013494 | 0.97984 | 0.0088434 |
| PC O-24:1_20:3 | 0.97234 | -0.040461 | 0.25195 | 0.59868 |
| PC O-24:1_22:6 | 0.97281 | -0.03977 | 0.89948 | 0.046011 |
| PC O-24:2_20:4 | 0.94724 | -0.078198 | 0.82009 | 0.086138 |
| PC O-28:0 | 0.78106 | -0.35649 | 0.059596 | 1.2248 |
| PC O-30:0 | 0.87237 | -0.19699 | 0.21156 | 0.67458 |
| PC O-30:1 | 0.96358 | -0.05352 | 0.40255 | 0.39518 |
| PC O-31:0 | 0.77865 | -0.36096 | 0.11793 | 0.92839 |
| PC O-32:0 | 0.98883 | -0.016205 | 0.66725 | 0.17571 |
| PC O-32:1 | 0.98855 | -0.016614 | 0.82009 | 0.086138 |
| PC O-32:2 | 0.90596 | -0.14247 | 0.32173 | 0.49251 |
| PC O-33:2 | 1.1706 | 0.2272 | 0.85962 | 0.065695 |
| PC O-34:1 | 0.97763 | -0.032645 | 0.56045 | 0.25146 |
| PC O-34:2 | 0.91923 | -0.1215 | 0.21156 | 0.67458 |
| PC O-34:3 | 0.9928 | -0.010419 | 0.52671 | 0.27843 |
| PC O-35:2 | 0.86869 | -0.20309 | 0.29723 | 0.5269 |
| PC O-36:0 | 0.99598 | -0.0058146 | 0.82009 | 0.086138 |
| PC O-36:1 | 0.98907 | -0.015857 | 0.63079 | 0.20012 |
| PC O-36:2 | 0.83017 | -0.26852 | 0.29723 | 0.5269 |

| | | | | |
|---|---|---|---|---|
| PC O-36:4 | 0.97371 | -0.03843 | 0.89948 | 0.046011 |
| PC O-36:5 | 0.96978 | -0.044278 | 0.85962 | 0.065695 |
| PC O-37:4 | 0.91511 | -0.12799 | 0.85962 | 0.065695 |
| PC O-37:5 | 0.81442 | -0.29616 | 0.70451 | 0.15211 |
| PC O-38:0 | 0.85419 | -0.22737 | 0.34745 | 0.45911 |
| PC O-38:2 | 0.97378 | -0.038334 | 0.59516 | 0.22537 |
| PC O-38:4 | 0.99994 | -9.34E-05 | 0.97984 | 0.0088434 |
| PC O-38:5 | 0.98287 | -0.024931 | 0.97984 | 0.0088434 |
| PC O-38:6 | 0.85488 | -0.2262 | 0.21156 | 0.67458 |
| PC O-39:6 | 0.73921 | -0.43594 | 0.29723 | 0.5269 |
| PC O-39:7 | 0.71276 | -0.48852 | 0.21156 | 0.67458 |
| PC O-40:1 | 1.1484 | 0.19964 | 0.70451 | 0.15211 |
| PC O-40:3 | 0.90498 | -0.14405 | 0.1759 | 0.75473 |
| PC O-40:4 | 1.1329 | 0.17998 | 0.43189 | 0.36462 |
| PC O-40:5 | 0.95577 | -0.065271 | 0.63079 | 0.20012 |
| PC O-40:6 | 0.81903 | -0.28801 | 0.3744 | 0.42667 |
| PC O-40:7 | 0.90055 | -0.15113 | 0.59516 | 0.22537 |
| PC O-40:8 | 1.0446 | 0.062956 | 0.89948 | 0.046011 |
| PC O-42:2 | 1.0214 | 0.030494 | 0.93959 | 0.027062 |
| PC O-42:5 | 0.90236 | -0.14822 | 0.46239 | 0.33499 |
| PC O-42:6 | 0.91947 | -0.12112 | 0.82009 | 0.086138 |
| PC O-42:7 | 0.90173 | -0.14923 | 0.59516 | 0.22537 |
| PC O-42:8 | 0.70933 | -0.49547 | 0.40255 | 0.39518 |
| PC O-43:4 | 0.96778 | -0.047242 | 0.85962 | 0.065695 |
| PC O-43:5 | 0.92909 | -0.10611 | 0.66725 | 0.17571 |
| PC O-44:5 | 1.0163 | 0.023372 | 0.56045 | 0.25146 |
| PC O-44:7 | 0.87138 | -0.19862 | 0.93959 | 0.027062 |
| PC O-46:6 | 0.93304 | -0.099983 | 0.63079 | 0.20012 |
| PC O-46:7 | 0.90278 | -0.14755 | 1 | 0 |
| PC O-48:8 | 0.73631 | -0.44162 | 0.52671 | 0.27843 |
| PE 16:0_16:1 | 1.2027 | 0.26632 | 0.66725 | 0.17571 |
| PE 16:0_18:1 | 1.0988 | 0.13598 | 0.82009 | 0.086138 |
| PE 16:0_18:2 | 1.0958 | 0.13204 | 0.63079 | 0.20012 |
| PE 16:0_20:3 | 0.96835 | -0.046393 | 0.63079 | 0.20012 |
| PE 16:0_20:4 | 1.0612 | 0.085667 | 0.78101 | 0.10734 |
| PE 16:0_20:5 | 0.8628 | -0.21291 | 0.63079 | 0.20012 |
| PE 17:0_20:4 | 0.92237 | -0.11658 | 0.89948 | 0.046011 |
| PE 18:0_18:1 | 1.0319 | 0.045316 | 0.70451 | 0.15211 |
| PE 18:0_20:3 | 0.93308 | -0.099932 | 0.70451 | 0.15211 |
| PE 18:0_20:4 | 1.0594 | 0.083228 | 0.40255 | 0.39518 |
| PE 18:0_22:6 | 0.89402 | -0.16163 | 0.93959 | 0.027062 |
| PE 18:1_18:1 | 1.1431 | 0.19293 | 0.32173 | 0.49251 |
| PE 18:1_18:2 | 1.0199 | 0.028473 | 0.85962 | 0.065695 |
| PE 30:0 | 0.91013 | -0.13585 | 0.63079 | 0.20012 |

| | | | | |
|---|---|---|---|---|
| PE 36:2 | 0.9034 | -0.14656 | 0.74245 | 0.12933 |
| PE 38:5 | 1.0757 | 0.10524 | 0.32173 | 0.49251 |
| PE 38:6 | 0.8586 | -0.21995 | 0.82009 | 0.086138 |
| PE O-16:0_18:2 | 0.9971 | -0.0041842 | 0.32173 | 0.49251 |
| PE O-16:1_18:1 | 1.0965 | 0.13293 | 0.49401 | 0.30626 |
| PE O-16:1_18:2 | 1.1462 | 0.19686 | 0.89948 | 0.046011 |
| PE O-16:1_20:3 | 1.0114 | 0.016297 | 1 | 0 |
| PE O-16:1_20:4 | 1.0697 | 0.097177 | 0.59516 | 0.22537 |
| PE O-16:1_20:5 | 0.96764 | -0.047463 | 0.89948 | 0.046011 |
| PE O-16:1_22:6 | 0.97413 | -0.037808 | 0.93959 | 0.027062 |
| PE O-18:0_18:1 | 0.95142 | -0.071847 | 0.59516 | 0.22537 |
| PE O-18:0_20:4 | 0.78396 | -0.35115 | 0.74245 | 0.12933 |
| PE O-18:1_18:1 | 0.96542 | -0.050777 | 0.93959 | 0.027062 |
| PE O-18:1_18:2 | 1.0262 | 0.037374 | 0.82009 | 0.086138 |
| PE O-18:1_20:3 | 1.0112 | 0.016077 | 0.93959 | 0.027062 |
| PE O-18:1_20:4 | 1.1076 | 0.14744 | 0.52671 | 0.27843 |
| PE O-18:1_22:4 | 1.2517 | 0.32392 | 0.1759 | 0.75473 |
| PE O-18:1_22:5 | 1.099 | 0.13619 | 0.52671 | 0.27843 |
| PE O-18:1_22:6 | 0.98307 | -0.024641 | 0.85962 | 0.065695 |
| PE O-18:2_18:1 | 0.99987 | -0.0001823 | 0.70451 | 0.15211 |
| PE O-18:2_22:6 | 1.0339 | 0.048085 | 0.89948 | 0.046011 |
| PE O-19:1_20:4 | 0.9326 | -0.10068 | 0.93959 | 0.027062 |
| PE O-20:1_18:1 | 0.86972 | -0.20137 | 1 | 0 |
| PE O-20:1_18:2 | 1.1057 | 0.14497 | 0.78101 | 0.10734 |
| PE O-20:1_20:4 | 1.049 | 0.069076 | 0.82009 | 0.086138 |
| PE O-20:1_22:6 | 0.97627 | -0.034649 | 0.74245 | 0.12933 |
| PE O-20:2_22:6 | 1.2355 | 0.30511 | 0.46239 | 0.33499 |
| PE O-22:1_18:2 | 1.0562 | 0.078941 | 0.78101 | 0.10734 |
| PE O-22:1_20:4 | 1.1033 | 0.14186 | 0.93959 | 0.027062 |
| PE O-24:2_20:4 | 1.1536 | 0.20618 | 0.3744 | 0.42667 |
| PE O-24:2_22:6 | 1.104 | 0.14273 | 0.89948 | 0.046011 |
| PE O-36:4 | 0.97347 | -0.038793 | 0.56045 | 0.25146 |
| PE O-38:6 | 1.1047 | 0.14368 | 0.63079 | 0.20012 |
| PE O-38:7 | 1.0335 | 0.047492 | 1 | 0 |
| PE-Cer (d32:1) | 0.86904 | -0.2025 | 0.49401 | 0.30626 |
| PE-Cer (d34:1) | 1.0478 | 0.067392 | 0.78101 | 0.10734 |
| PE-Cer (d40:1) | 1.0514 | 0.07236 | 0.66725 | 0.17571 |
| PE-Cer (d42:2) | 1.1453 | 0.19573 | 0.49401 | 0.30626 |
| PE-Cer (d42:3) | 1.2271 | 0.29528 | 0.40255 | 0.39518 |
| PI 16:0_16:1 | 0.90971 | -0.13653 | 0.49401 | 0.30626 |
| PI 16:0_20:4 | 0.93727 | -0.093466 | 0.52671 | 0.27843 |
| PI 18:0_18:2 | 0.91484 | -0.1284 | 0.66725 | 0.17571 |
| PI 18:0_18:3 | 0.94136 | -0.087176 | 0.49401 | 0.30626 |
| PI 18:0_20:4 | 0.91604 | -0.12652 | 0.63079 | 0.20012 |

| | | | | |
|---|---|---|---|---|
| PI 18:0_22:4 | 0.93885 | -0.091033 | 0.46239 | 0.33499 |
| PI 18:0_22:5 | 0.88283 | -0.17979 | 0.43189 | 0.36462 |
| PI 18:0_22:6 | 0.80951 | -0.30488 | 0.27398 | 0.56228 |
| PI 18:1_18:2 | 1.0229 | 0.03264 | 0.78101 | 0.10734 |
| PI 32:0 | 0.98979 | -0.014801 | 0.82009 | 0.086138 |
| PI 34:2 | 0.90426 | -0.1452 | 0.82009 | 0.086138 |
| PI 37:4 | 0.82909 | -0.2704 | 0.49401 | 0.30626 |
| PI 38:2 | 0.74004 | -0.43432 | 0.19315 | 0.7141 |
| PI 38:4 | 0.81029 | -0.30348 | 0.23115 | 0.6361 |
| PS 18:1_20:4 | 0.94813 | -0.076844 | 0.29723 | 0.5269 |
| PS 18:2_20:3 | 1.1201 | 0.16362 | 0.52671 | 0.27843 |
| PS 38:4 | 1.2109 | 0.27607 | 0.075691 | 1.121 |
| SM 29:1;O2 | 0.70665 | -0.50093 | 0.21156 | 0.67458 |
| SM 30:1;O2 | 0.81815 | -0.28956 | 0.34745 | 0.45911 |
| SM 31:0;O2 | 0.59732 | -0.74342 | 0.21156 | 0.67458 |
| SM 31:1;O2 | 0.70831 | -0.49755 | 0.095012 | 1.0222 |
| SM 31:2 | 0.8187 | -0.2886 | 0.56045 | 0.25146 |
| SM 32:0;O2 | 0.75759 | -0.4005 | 0.11793 | 0.92839 |
| SM 32:1;O2 | 0.85216 | -0.23081 | 0.23115 | 0.6361 |
| SM 32:1;O3 | 0.82464 | -0.27816 | 0.1759 | 0.75473 |
| SM 32:2;O2 | 0.88614 | -0.17439 | 0.78101 | 0.10734 |
| SM 32:2;O3 | 0.70073 | -0.51306 | 0.01267 | 1.8972 |
| SM 33:0;O2 | 0.7212 | -0.47152 | 0.23115 | 0.6361 |
| SM 33:1;O2 | 0.81421 | -0.29653 | 0.23115 | 0.6361 |
| SM 33:1;O3 | 0.61379 | -0.70419 | 0.031043 | 1.508 |
| SM 33:2;O2 | 0.89684 | -0.15707 | 0.70451 | 0.15211 |
| SM 34:0;O2 | 0.88451 | -0.17705 | 0.1598 | 0.79643 |
| SM 34:1;O2 | 1.0304 | 0.043192 | 0.78101 | 0.10734 |
| SM 34:1;O3 | 1.0548 | 0.076949 | 0.85962 | 0.065695 |
| SM 34:2;O2 | 0.97099 | -0.042475 | 0.89948 | 0.046011 |
| SM 34:2;O3 | 0.95218 | -0.070686 | 0.70451 | 0.15211 |
| SM 34:3;O2 | 0.88318 | -0.17921 | 0.63079 | 0.20012 |
| SM 35:0;O2 | 0.87482 | -0.19293 | 0.78101 | 0.10734 |
| SM 35:0;O2_2 | 0.73213 | -0.44983 | 0.1759 | 0.75473 |
| SM 35:1;O2 | 0.90295 | -0.14728 | 0.40255 | 0.39518 |
| SM 35:2;O2 | 0.86427 | -0.21044 | 0.3744 | 0.42667 |
| SM 36:1;O2 | 0.93994 | -0.089366 | 0.78101 | 0.10734 |
| SM 36:2;O2 | 0.95929 | -0.059956 | 0.49401 | 0.30626 |
| SM 36:2;O3 | 1.0595 | 0.083394 | 0.70451 | 0.15211 |
| SM 36:3;O2 | 0.8439 | -0.24485 | 0.34745 | 0.45911 |
| SM 37:1;O2 | 0.88416 | -0.17761 | 0.29723 | 0.5269 |
| SM 37:1;O2_2 | 0.81574 | -0.29382 | 0.70451 | 0.15211 |
| SM 37:2;O2 | 0.96431 | -0.052424 | 0.56045 | 0.25146 |
| SM 38:0;O2 | 0.86708 | -0.20576 | 0.59516 | 0.22537 |

| | | | | |
|---|---|---|---|---|
| SM 38:1;O2 | 0.86824 | -0.20383 | 0.3744 | 0.42667 |
| SM 38:2;O2 | 1.0093 | 0.013376 | 0.82009 | 0.086138 |
| SM 38:3;O2 | 1.0422 | 0.059662 | 0.89948 | 0.046011 |
| SM 39:0;O2 | 0.77399 | -0.36962 | 0.66725 | 0.17571 |
| SM 39:1;O2 | 0.77361 | -0.37032 | 0.70451 | 0.15211 |
| SM 39:2;O2 | 0.97008 | -0.043825 | 0.85962 | 0.065695 |
| SM 39:2;O2_2 | 0.9082 | -0.13892 | 0.63079 | 0.20012 |
| SM 40:0;O2 | 0.82521 | -0.27717 | 0.27398 | 0.56228 |
| SM 40:0;O3 | 0.65057 | -0.62022 | 0.0044802 | 2.3487 |
| SM 40:1;O2 | 0.85153 | -0.23188 | 0.46239 | 0.33499 |
| SM 40:2;O2 | 0.96576 | -0.05026 | 0.63079 | 0.20012 |
| SM 40:2;O2_2 | 0.96444 | -0.052231 | 0.82009 | 0.086138 |
| SM 40:3;O2 | 0.97843 | -0.031453 | 0.82009 | 0.086138 |
| SM 41:1;O2 | 0.87833 | -0.18717 | 0.34745 | 0.45911 |
| SM 41:2;O2 | 0.91726 | -0.12459 | 0.43189 | 0.36462 |
| SM 41:2;O2_2 | 0.96233 | -0.055403 | 0.82009 | 0.086138 |
| SM 41:3;O2 | 0.92134 | -0.11819 | 0.74245 | 0.12933 |
| SM 42:0;O2 | 0.73625 | -0.44174 | 0.059596 | 1.2248 |
| SM 42:0;O3 | 0.69837 | -0.51793 | 0.023364 | 1.6315 |
| SM 42:1;O2 | 0.83982 | -0.25185 | 0.34745 | 0.45911 |
| SM 42:1;O3 | 0.87588 | -0.19119 | 0.27398 | 0.56228 |
| SM 42:2;O2 | 0.98831 | -0.016964 | 0.93959 | 0.027062 |
| SM 42:2;O2_2 | 0.96003 | -0.05885 | 0.85962 | 0.065695 |
| SM 42:2;O3 | 1.1003 | 0.13794 | 0.63079 | 0.20012 |
| SM 42:3;O2 | 1.0428 | 0.060523 | 0.66725 | 0.17571 |
| SM 42:3;O3 | 1.2906 | 0.36803 | 106 | 0.97469 |
| SM 42:4;O2_2 | 0.88701 | -0.17298 | 0.70451 | 0.15211 |
| SM 43:1;O2 | 0.75908 | -0.39767 | 0.13084 | 0.88324 |
| SM 43:2;O2 | 0.82543 | -0.27679 | 0.74245 | 0.12933 |
| SM 43:3;O2 | 0.84783 | -0.23815 | 1 | 0 |
| SM 43:3;O2_2 | 0.78767 | -0.34434 | 0.59516 | 0.22537 |
| SM 44:1;O2 | 0.83704 | -0.25664 | 0.43189 | 0.36462 |
| SM 44:2;O2 | 1.11 | 0.15062 | 0.74245 | 0.12933 |
| SM 44:3;O2 | 1.0612 | 0.085763 | 0.63079 | 0.20012 |
| ST 24:1;O3;S | 2.2796 | 1.1888 | 0.34745 | 0.45911 |
| ST 27:1;O;S | 0.93127 | -0.10272 | 0.93959 | 0.027062 |
| ST 28:1;O;S | 1.1027 | 0.14107 | 0.97984 | 0.0088434 |
| TG 27:0 | 0.99181 | -0.011859 | 0.74245 | 0.12933 |
| TG 28:0 | 0.97758 | -0.032715 | 0.66725 | 0.17571 |
| TG 29:0 | 0.94205 | -0.08613 | 0.59516 | 0.22537 |
| TG 30:0 | 972 | -0.040971 | 0.59516 | 0.22537 |
| TG 32:0 | 0.77863 | -361 | 0.63079 | 0.20012 |
| TG 36:1 | 1.5287 | 0.6123 | 0.095012 | 1.0222 |
| TG 36:2 | 1.9482 | 0.96211 | 0.035598 | 1.4486 |

| | | | | |
|---|---|---|---|---|
| TG 38:0 | 1.266 | 0.34031 | 0.25195 | 0.59868 |
| TG 38:1 | 1.7083 | 0.77258 | 0.059596 | 1.2248 |
| TG 38:2 | 2.3029 | 1.2035 | 0.01267 | 1.8972 |
| TG 40:0 | 0.74202 | -0.43048 | 0.78101 | 0.10734 |
| TG 41:0 | 0.50661 | -0.98105 | 0.63079 | 0.20012 |
| TG 42:0 | 0.6288 | -0.66933 | 0.78101 | 0.10734 |
| TG 42:1 | 0.85 | -0.23447 | 0.56045 | 0.25146 |
| TG 42:2 | 2.2625 | 1.1779 | 0.46239 | 0.33499 |
| TG 42:3 | 1.7324 | 0.79278 | 0.25195 | 0.59868 |
| TG 43:1 | 0.41818 | -1.2578 | 0.46239 | 0.33499 |
| TG 44:0 | 0.85643 | -0.2236 | 0.82009 | 0.086138 |
| TG 44:1 | 0.55196 | -0.85737 | 0.89948 | 0.046011 |
| TG 44:2 | 1.0399 | 0.056437 | 0.40255 | 0.39518 |
| TG 44:3 | 2.3878 | 1.2557 | 0.11793 | 0.92839 |
| TG 44:5 | 2.7361 | 1.4521 | 0.11793 | 0.92839 |
| TG 45:0 | 0.6816 | -0.55301 | 0.63079 | 0.20012 |
| TG 45:1 | 0.56978 | -0.81152 | 0.63079 | 0.20012 |
| TG 46:0 | 0.87332 | -0.19542 | 0.52671 | 0.27843 |
| TG 46:1 | 0.65543 | -0.60949 | 0.78101 | 0.10734 |
| TG 46:2 | 0.47787 | -1.0653 | 0.85962 | 0.065695 |
| TG 46:5 | 1.3809 | 0.46565 | 0.74245 | 0.12933 |
| TG 47:1 | 0.55953 | -0.83771 | 0.59516 | 0.22537 |
| TG 47:2 | 0.48714 | -1.0376 | 0.52671 | 0.27843 |
| TG 47:3 | 0.54937 | -0.86415 | 0.66725 | 0.17571 |
| TG 48:1 | 0.7508 | -0.41349 | 0.78101 | 0.10734 |
| TG 49:1 | 0.62466 | -0.67886 | 0.49401 | 0.30626 |
| TG 49:3 | 0.46879 | -1.093 | 0.40255 | 0.39518 |
| TG 51:1 | 0.67641 | -0.56403 | 0.78101 | 0.10734 |
| TG 51:2 | 0.65699 | -0.60606 | 0.59516 | 0.22537 |
| TG 51:4 | 0.55215 | -0.85686 | 0.1598 | 0.79643 |
| TG 52:7 | 0.46262 | -1.1121 | 0.66725 | 0.17571 |
| TG 52:8 | 0.56626 | -0.82047 | 0.29723 | 0.5269 |
| TG 53:1 | 0.7854 | -0.3485 | 1 | 0 |
| TG 53:2 | 0.65363 | -0.61346 | 0.66725 | 0.17571 |
| TG 53:3 | 0.6471 | -0.62794 | 0.34745 | 0.45911 |
| TG 53:4 | 0.71254 | -0.48896 | 0.27398 | 0.56228 |
| TG 53:5 | 0.65861 | -0.60251 | 0.19315 | 0.7141 |
| TG 54:1 | 0.74153 | -0.43142 | 0.93959 | 0.027062 |
| TG 54:3 | 0.8083 | -0.30704 | 0.93959 | 0.027062 |
| TG 54:4 | 0.8963 | -0.15795 | 0.46239 | 0.33499 |
| TG 54:5 | 0.86509 | -0.20907 | 0.82009 | 0.086138 |
| TG 54:7 | 0.56443 | -0.82513 | 0.1598 | 0.79643 |
| TG 54:8 | 0.44599 | -1.1649 | 0.1598 | 0.79643 |
| TG 54:9 | 0.75457 | -0.40628 | 0.85962 | 0.065695 |

| | | | | |
|---|---|---|---|---|
| TG 55:2 | 0.75769 | -0.40032 | 0.82009 | 0.086138 |
| TG 55:3 | 0.73044 | -0.45316 | 0.66725 | 0.17571 |
| TG 55:6 | 0.75035 | -0.41437 | 0.32173 | 0.49251 |
| TG 55:8 | 0.43901 | -1.1877 | 0.059596 | 1.2248 |
| TG 56:1 | 0.64982 | -0.62188 | 0.89948 | 0.046011 |
| TG 56:10 | 0.49098 | -1.0263 | 0.19315 | 0.7141 |
| TG 56:2 | 0.81421 | -0.29653 | 0.85962 | 0.065695 |
| TG 56:3 | 0.77577 | -0.3663 | 0.82009 | 0.086138 |
| TG 56:5 | 0.83082 | -0.2674 | 0.66725 | 0.17571 |
| TG 56:7 | 0.67405 | -0.56907 | 0.46239 | 0.33499 |
| TG 56:8 | 0.5051 | -0.98535 | 0.067266 | 1.1722 |
| TG 56:9 | 0.45404 | -1.1391 | 0.21156 | 0.67458 |
| TG 57:2 | 0.9772 | -0.033276 | 0.3744 | 0.42667 |
| TG 57:4 | 0.81476 | -0.29555 | 0.74245 | 0.12933 |
| TG 57:7 | 0.74311 | -0.42836 | 0.56045 | 0.25146 |
| TG 58:1 | 0.69187 | -0.53144 | 0.78101 | 0.10734 |
| TG 58:10 | 0.78784 | -0.34403 | 0.3744 | 0.42667 |
| TG 58:11 | 0.57281 | -0.80388 | 0.43189 | 0.36462 |
| TG 58:4 | 0.98333 | -0.024252 | 0.56045 | 0.25146 |
| TG 58:5 | 0.91342 | -0.13065 | 0.59516 | 0.22537 |
| TG 58:6 | 0.79995 | -0.32201 | 0.49401 | 0.30626 |
| TG 58:7 | 0.80138 | -0.31944 | 0.82009 | 0.086138 |
| TG 58:8 | 0.7525 | -0.41023 | 0.34745 | 0.45911 |
| TG 58:9 | 0.69482 | -0.5253 | 0.1598 | 0.79643 |
| TG 59:2 | 737 | -0.44027 | 0.93959 | 0.027062 |
| TG 59:3 | 1.0299 | 0.042465 | 0.59516 | 0.22537 |
| TG 59:4 | 1.0283 | 0.040298 | 0.82009 | 0.086138 |
| TG 60:11 | 0.73781 | -0.43868 | 0.27398 | 0.56228 |
| TG 60:13 | 0.95832 | -0.061425 | 0.52671 | 0.27843 |
| TG 60:3 | 0.94006 | -0.08917 | 0.59516 | 0.22537 |
| TG 60:4 | 0.93224 | -0.10122 | 0.63079 | 0.20012 |
| TG 60:5 | 0.8759 | -0.19116 | 0.93959 | 0.027062 |
| TG 60:7 | 0.78852 | -0.34279 | 0.63079 | 0.20012 |
| TG 61:4 | 0.8503 | -0.23395 | 0.85962 | 0.065695 |
| TG 62:12 | 0.71976 | -0.47442 | 0.1598 | 0.79643 |
| TG 62:13 | 0.87713 | -0.18914 | 0.3744 | 0.42667 |
| TG 62:2 | 0.70008 | -0.5144 | 0.97984 | 0.0088434 |
| TG 62:3 | 0.70369 | -0.50698 | 0.97984 | 0.0088434 |
| TG O-50:1 | 0.69608 | -0.52267 | 0.56045 | 0.25146 |
| TG O-58:3 | 0.67711 | -0.56254 | 0.19315 | 0.7141 |

**Supplementary Table 3:** Full annotation of Fig. 3D

| Position | Lipid |
|---|---|
| 1 | Cer 18:0;O2/16:0 |
| 2 | Cer 18:0;O2/22:0 |
| 3 | Cer 18:0;O2/23:0 |
| 4 | Cer 18:0;O2/24:2 |
| 5 | Cer 18:0;O3/20:0 |
| 6 | Cer 18:0;O3/23:0 |
| 7 | Cer 18:1;O2/26:0 |
| 8 | Hex2Cer 18:1;O2/16:0 |
| 9 | Hex2Cer 18:1;O2/22:0 |
| 10 | Hex2Cer 18:1;O2/24:0 |
| 11 | Hex2Cer 18:1;O2/24:1 |
| 12 | Hex3Cer 18:1;O2/16:0 |
| 13 | Hex3Cer 18:1;O2/22:0 |
| 14 | Hex3Cer 18:1;O2/24:0 |
| 15 | Hex3Cer 18:1;O2/24:1 |
| 16 | HexCer 18:1;O2/16:0 |
| 17 | HexCer 18:1;O2/20:0 |
| 18 | HexCer 18:1;O2/22:0 |
| 19 | HexCer 18:1;O2/24:0 |
| 20 | HexCer 18:1;O2/24:1 |
| 21 | SM 28:1;O2 |
| 22 | SM 30:1;O2 |
| 23 | SM 32:0;O2 |
| 24 | SM 32:1;O2 |
| 25 | SM 32:1;O3 |
| 26 | SM 32:2;O2 |
| 27 | SM 33:0;O2 |
| 28 | SM 33:1;O2 |
| 29 | SM 33:1;O3 |
| 30 | SM 33:2;O2 |
| 31 | SM 34:0;O2 |
| 32 | SM 34:0;O3 |
| 33 | SM 34:1;O2 |
| 34 | SM 34:1;O3 |
| 35 | SM 34:2;O2 |
| 36 | SM 34:2;O3 |
| 37 | SM 35:0;O2 |
| 38 | SM 35:1;O2 |
| 39 | SM 35:2;O2 |
| 40 | SM 39:2;O2_2 |
| 41 | SM 40:0;O2 |
| 42 | SM 40:2;O2_2 |
| 43 | SM 40:3;O2 |
| 44 | SM 41:0;O2 |

| | |
|---|---|
| 45 | SM 41:0;O3 |
| 46 | SM 41:1;O2_1 |
| 47 | SM 41:1;O2_2 |
| 48 | SM 41:2;O2_2 |
| 49 | SM 41:3;O2 |
| 50 | SM 42:0;O2 |
| 51 | SM 42:1;O2 |
| 52 | SM 42:1;O3 |
| 53 | SM 42:2;O2 |
| 54 | SM 42:3;O2 |
| 55 | SM 43:2;O2 |
| 56 | SM 44:1;O2 |
| 57 | SM 44:2;O2 |
| 58 | SM 44:3;O2 |
| 59 | SM 44:4;O2 |

**Supplementary Table 4:** Full annotation of Fig. 4C

| Position | Lipid |
|---|---|
| 1 | Cer 17:1;O2/16:0 |
| 2 | Cer 18:0;O2/16:0 |
| 3 | Cer 18:0;O2/18:0 |
| 4 | Cer 18:0;O2/20:0 |
| 5 | Cer 18:0;O2/21:0 |
| 6 | Cer 18:0;O2/22:0 |
| 7 | Cer 18:0;O2/23:0 |
| 8 | Cer 18:0;O2/24:0 |
| 9 | Cer 18:0;O2/24:1 |
| 10 | Cer 18:0;O2/25:0 |
| 11 | Cer 18:0;O3/16:0 |
| 12 | Cer 18:0;O3/16:0;(2OH) |
| 13 | Cer 18:0;O3/20:0 |
| 14 | Cer 18:0;O3/21:0 |
| 15 | Cer 18:0;O3/22:0;(2OH) |
| 16 | Cer 18:0;O3/22:1 |
| 17 | Cer 18:0;O3/24:0;(2OH) |
| 18 | Cer 18:0;O3/24:1;(2OH) |
| 19 | Cer 18:1;O2/16:0 |
| 20 | Cer 18:1;O2/16:0;(2OH) |
| 21 | Cer 18:1;O2/18:0 |
| 22 | Cer 18:1;O2/20:0 |
| 23 | Cer 18:1;O2/22:0 |
| 24 | Cer 18:1;O2/22:1 |
| 25 | Cer 18:1;O2/23:0 |
| 26 | Cer 18:1;O2/24:0 |
| 27 | Cer 18:1;O2/24:1 |
| 28 | Cer 18:1;O2/24:2 |
| 29 | Cer 18:1;O2/25:0 |
| 30 | Cer 18:1;O2/34:3 |
| 31 | Cer 18:2;O2/22:0 |
| 32 | Hex2Cer 18:1;O2/16:0 |
| 33 | Hex2Cer 18:1;O2/22:0 |
| 34 | Hex2Cer 18:1;O2/24:0 |
| 35 | Hex2Cer 18:1;O2/24:1 |
| 36 | Hex3Cer 18:1;O2/16:0 |
| 37 | Hex3Cer 18:1;O2/24:1 |
| 38 | HexCer 18:0;O2/22:0 |
| 39 | HexCer 18:0;O3/18:0;(2OH) |
| 40 | HexCer 18:0;O3/22:0;(2OH) |
| 41 | HexCer 18:0;O3/24:0;(2OH) |
| 42 | HexCer 18:0;O3/24:1;(2OH) |
| 43 | HexCer 18:1;O2/16:1 |
| 44 | HexCer 18:1;O2/18:0 |

| | |
|---|---|
| 45 | HexCer 18:1;O2/20:0 |
| 46 | HexCer 18:1;O2/22:0 |
| 47 | HexCer 18:1;O2/22:1 |
| 48 | HexCer 18:1;O2/23:0 |
| 49 | HexCer 18:1;O2/24:0 |
| 50 | HexCer 18:1;O2/24:1 |
| 51 | HexCer 18:1;O2/25:0 |
| 52 | HexCer 18:2;O2/22:0 |
| 53 | HexCer 18:2;O2/24:0 |
| 54 | PE-Cer (d34:0) |
| 55 | PE-Cer (d42:2) |
| 56 | SM 30:1;O2 |
| 57 | SM 30:1;O3 |
| 58 | SM 31:1;O2 |
| 59 | SM 31:1;O3 |
| 60 | SM 32:0;O2 |
| 61 | SM 32:1;O2 |
| 62 | SM 32:1;O3 |
| 63 | SM 33:0;O2 |
| 64 | SM 33:0;O3 |
| 65 | SM 33:1;O2 |
| 66 | SM 33:1;O3 |
| 67 | SM 33:2;O2 |
| 68 | SM 34:1;O2 |
| 69 | SM 34:1;O3 |
| 70 | SM 34:2;O2 |
| 71 | SM 34:2;O3 |
| 72 | SM 35:1;O2 |
| 73 | SM 35:2;O2 |
| 74 | SM 36:0;O2 |
| 75 | SM 36:1;O2 |
| 76 | SM 36:2;O2 |
| 77 | SM 36:2;O3 |
| 78 | SM 36:3;O2 |
| 79 | SM 37:1;O2 |
| 80 | SM 38:0;O2 |
| 81 | SM 38:1;O2 |
| 82 | SM 38:2;O2 |
| 83 | SM 39:1;O2 |
| 84 | SM 39:2;O2_A |
| 85 | SM 39:2;O2_B |
| 86 | SM 40:0;O2 |
| 87 | SM 40:0;O3 |
| 88 | SM 40:1;O2 |
| 89 | SM 40:1;O3 |
| 90 | SM 40:2;O2_A |
| 91 | SM 40:2;O2_B |

| | |
|---|---|
| 92 | SM 40:3;O2 |
| 93 | SM 40:4;O2 |
| 94 | SM 41:0;O3 |
| 95 | SM 41:1;O2 |
| 96 | SM 41:2;O2_A |
| 97 | SM 41:2;O2_B |
| 98 | SM 41:3;O2 |
| 99 | SM 42:0;O2 |
| 100 | SM 42:0;O3 |
| 101 | SM 42:1;O2_A |
| 102 | SM 42:1;O2_B |
| 103 | SM 42:1;O3 |
| 104 | SM 42:2;O2_A |
| 105 | SM 42:2;O2_B |
| 106 | SM 42:2;O3 |
| 107 | SM 42:3;O2 |
| 108 | SM 42:3;O3 |
| 109 | SM 42:5;O2 |
| 110 | SM 43:1;O2 |
| 111 | SM 43:2;O2 |
| 112 | SM 43:3;O2 |
| 113 | SM 44:1;O2 |
| 114 | SM 44:2;O2 |
| 115 | SM 44:3;O2 |
| 116 | SM 44:5;O2 |

**Supplementary Table 5**: Baseline characteristics of Transcatheter Aortic Valve Implantation (TAVI) patients for plasma Lipidomics analysis

| **Parameter** | **n=70** |
|---|---|
| **Age** | 80.1 ± 8.6 |
| **Female** | 60% |
| **BMI** | 26.1 ± 4.6 |
| **Hypercholesterolemia** | 21% |
| **Hypertension** | 65% |

**Supplementary Table 6:** Target proteins and lipids for partial correlation analysis between lipidomics and proteomics data

| Gene Name | Assay |
|---|---|
| ACAN | protein |
| APOC4 | protein |
| APOL4 | protein |
| ASB6 | protein |
| CRP | protein |
| CTHRC1 | protein |
| DES | protein |
| EPB42 | protein |
| ERAP2 | protein |
| FABP4 | protein |
| FBXO16 | protein |
| GFAP | protein |
| GNG5 | protein |
| GPD1 | protein |
| GSTM1 | protein |
| HBG1 | protein |
| HSPA2 | protein |
| IGHD | protein |
| IGHV1-45 | protein |
| IGHV1-69 | protein |
| IGKV1-13 | protein |
| IGLV2-14 | protein |
| IGLV8-61 | protein |
| KMT2D | protein |
| KRT1 | protein |
| LPA | protein |
| MAPK15 | protein |
| MYL1 | protein |
| MYL2 | protein |
| MYL3 | protein |
| PF4 | protein |
| PLIN1 | protein |
| PLIN4 | protein |
| POTEF | protein |
| PZP | protein |
| RALYL | protein |
| RHAG | protein |
| S100A8 | protein |
| S100A9 | protein |
| SERPINE1 | protein |

| | |
|---|---|
| SLC2A1 | protein |
| SLC4A1 | protein |
| SMG8 | protein |
| SMIM28 | protein |
| SPTA1 | protein |
| STRBP | protein |
| THBS2 | protein |
| TIMP1 | protein |
| TKTL2 | protein |
| TMPRSS6 | protein |
| TNC | protein |
| TNFRSF11B | protein |
| TPSAB1 | protein |
| Cer 16:0;O2/24:1 | lipid |
| Cer 16:0;O3/24:1 | lipid |
| Cer 16:1;O2/23:0 | lipid |
| Cer 16:1;O2/24:0;(2OH) | lipid |
| Cer 16:1;O2/24:1 | lipid |
| Cer 16:1;O2/24:2 | lipid |
| Cer 17:1;O2/16:0 | lipid |
| Cer 17:1;O2/24:1 | lipid |
| Cer 18:0;O2/20:0 | lipid |
| Cer 18:0;O2/22:0 | lipid |
| Cer 18:0;O2/23:0 | lipid |
| Cer 18:0;O2/24:0 | lipid |
| Cer 18:0;O2/24:2 | lipid |
| Cer 18:0;O3/16:0 | lipid |
| Cer 18:0;O3/22:0 | lipid |
| Cer 18:0;O3/23:0 | lipid |
| Cer 18:0;O3/24:0 | lipid |
| Cer 18:0;O3/24:1 | lipid |
| Cer 18:1;O2/16:0;(2OH) | lipid |
| Cer 18:1;O2/18:0 | lipid |
| Cer 18:1;O2/20:0 | lipid |
| Cer 18:1;O2/22:0 | lipid |
| Cer 18:1;O2/23:0 | lipid |
| Cer 18:1;O2/24:0;(2OH) | lipid |
| Cer 18:1;O2/24:1 | lipid |
| Cer 18:1;O2/24:1;(2OH) | lipid |
| Cer 18:1;O2/26:0 | lipid |
| Cer 18:1;O2/26:1 | lipid |
| Cer 18:1;O3/22:0 | lipid |
| Cer 18:1;O3/23:0 | lipid |
| Cer 18:1;O3/24:0 | lipid |

| | |
|---|---|
| Cer 18:1;O3/24:1 | lipid |
| Cer 18:2;O2/16:0 | lipid |
| Cer 18:2;O2/18:0 | lipid |
| Cer 18:2;O2/20:0 | lipid |
| Cer 18:2;O2/22:0 | lipid |
| Cer 18:2;O2/23:0 | lipid |
| Cer 18:2;O2/24:0 | lipid |
| Cer 18:2;O2/24:2 | lipid |
| Cer 18:2;O2/26:0 | lipid |
| Cer 19:1;O2/22:0 | lipid |
| Cer 19:1;O2/24:0 | lipid |
| Cer 37:1;O2 | lipid |
| Cer 42:3;O2 | lipid |
| Cer 44:3;O2 | lipid |
| GM3 33:1;O2 | lipid |
| Hex2Cer 18:1;O2/24:0 | lipid |
| Hex2Cer 18:1;O2/24:1 | lipid |
| Hex2Cer 18:2;O2/16:0 | lipid |
| Hex2Cer 18:2;O2/24:0 | lipid |
| Hex2Cer 18:2;O2/24:1 | lipid |
| Hex2Cer 32:1;O2 | lipid |
| Hex3Cer 16:1;O2/24:1 | lipid |
| Hex3Cer 18:1;O2/16:0 | lipid |
| Hex3Cer 18:1;O2/18:0 | lipid |
| Hex3Cer 18:1;O2/20:0 | lipid |
| Hex3Cer 18:1;O2/22:0 | lipid |
| Hex3Cer 18:1;O2/24:0 | lipid |
| Hex3Cer 18:1;O2/24:1 | lipid |
| Hex3Cer 18:2;O2/24:0 | lipid |
| Hex3Cer 32:1;O2 | lipid |
| Hex3Cer 41:1;O2 | lipid |
| Hex3Cer 44:1;O2 | lipid |
| HexCer 17:1;O2/22:0 | lipid |
| HexCer 18:0;O3/16:0;(2OH) | lipid |
| HexCer 18:1;O2/20:0 | lipid |
| HexCer 18:1;O2/22:0 | lipid |
| HexCer 18:1;O2/23:0 | lipid |
| HexCer 18:1;O2/24:0 | lipid |
| HexCer 18:1;O2/24:1 | lipid |
| HexCer 18:2;O2/22:0 | lipid |
| HexCer 18:2;O2/24:0 | lipid |
| PE-Cer D32:1 | lipid |
| PE-Cer D33:1 | lipid |
| PE-Cer D34:0 | lipid |

| | |
|---|---|
| PE-Cer D34:1 | lipid |
| PE-Cer D34:2 | lipid |
| PE-Cer D36:1 | lipid |
| PE-Cer D36:2 | lipid |
| PE-Cer D38:1 | lipid |
| PE-Cer D40:2 | lipid |
| PE-Cer D42:2 | lipid |
| PE-Cer D42:3 | lipid |
| SM 32:1;O2 | lipid |
| SM 38:2;O2 | lipid |
| SM 39:1;O2 | lipid |
| SM 39:2;O2 | lipid |
| SM 40:2;O2 | lipid |
| SM 40:3;O2 | lipid |
| SM 41:1;O2 | lipid |
| SM 41:1;O3 | lipid |
| SM 41:2;O2 | lipid |
| SM 42:2;O3 | lipid |

### References


1. Achner, L., T. Klersy, B. Fels, T. Reinberger, C.X. Schmidt, N. Groß, et al., *AFM-based nanoindentation indicates an impaired cortical stiffness in the AAV-PCSK9(DY) atherosclerosis mouse model.* Pflugers Arch, 2022. **474**(9): p. 993-1002.
2. Wu, Q., L. Sun, X. Hu, X. Wang, F. Xu, B. Chen, et al., *Suppressing the intestinal farnesoid X receptor/sphingomyelin phosphodiesterase 3 axis decreases atherosclerosis.* J Clin Invest, 2021. **131**(9).
3. Honda, S., T. Miyamoto, T. Watanabe, T. Narumi, S. Kadowaki, Y. Honda, et al., *A novel mouse model of aortic valve stenosis induced by direct wire injury.* Arterioscler Thromb Vasc Biol, 2014. **34**(2): p. 270-8.
4. Pachon, R.E., B.A. Scharf, D.E. Vatner, and S.F. Vatner, *Best anesthetics for assessing left ventricular systolic function by echocardiography in mice.* Am J Physiol Heart Circ Physiol, 2015. **308**(12): p. H1525-9.
5. Lee, S.H., N. Kim, M. Kim, S.-H. Woo, I. Han, J. Park, et al., *Single-cell transcriptomics reveal cellular diversity of aortic valve and the immunomodulation by PPARγ during hyperlipidemia.* Nature Communications, 2022. **13**(1): p. 5461.
6. Hao, Y., S. Hao, E. Andersen-Nissen, W.M. Mauck, 3rd, S. Zheng, A. Butler, et al., *Integrated analysis of multimodal single-cell data.* Cell, 2021. **184**(13): p. 3573-3587.e29.
7. Wolock, S.L., R. Lopez, and A.M. Klein, *Scrublet: Computational Identification of Cell Doublets in Single-Cell Transcriptomic Data.* Cell Syst, 2019. **8**(4): p. 281-291.e9.
8. Korsunsky, I., N. Millard, J. Fan, K. Slowikowski, F. Zhang, K. Wei, et al., *Fast, sensitive and accurate integration of single-cell data with Harmony.* Nature Methods, 2019. **16**(12): p. 1289-1296.
9. Liberzon, A., C. Birger, H. Thorvaldsdóttir, M. Ghandi, J.P. Mesirov, and P. Tamayo, *The Molecular Signatures Database (MSigDB) hallmark gene set collection.* Cell Syst, 2015. **1**(6): p. 417-425.

10. Ge, S.X., D. Jung, and R. Yao, *ShinyGO: a graphical gene-set enrichment tool for animals and plants.* Bioinformatics, 2020. **36**(8): p. 2628-2629.
11. Borcherding, N., A. Vishwakarma, A.P. Voigt, A. Bellizzi, J. Kaplan, K. Nepple, et al., *Mapping the immune environment in clear cell renal carcinoma by single-cell genomics.* Communications Biology, 2021. **4**(1): p. 122.
12. Subramanian, A., P. Tamayo, V.K. Mootha, S. Mukherjee, B.L. Ebert, M.A. Gillette, et al., *Gene set enrichment analysis: A knowledge-based approach for interpreting genome-wide expression profiles.* Proceedings of the National Academy of Sciences, 2005. **102**(43): p. 15545-15550.
13. Aherrahrou, R., A.E. Kulle, N. Alenina, R. Werner, S. Vens-Cappell, M. Bader, et al., *CYP17A1 deficient XY mice display susceptibility to atherosclerosis, altered lipidomic profile and atypical sex development.* Scientific Reports, 2020. **10**(1): p. 8792.
14. Karsai, G., F. Kraft, N. Haag, G.C. Korenke, B. Hänisch, A. Othman, et al., *DEGS1-associated aberrant sphingolipid metabolism impairs nervous system function in humans.* J Clin Invest, 2019. **129**(3): p. 1229-1239.
15. Kind, T., K.-H. Liu, D.Y. Lee, B. DeFelice, J.K. Meissen, and O. Fiehn, *LipidBlast in silico tandem mass spectrometry database for lipid identification.* Nature Methods, 2013. **10**(8): p. 755-758.
16. Meier, F., A.D. Brunner, M. Frank, A. Ha, I. Bludau, E. Voytik, et al., *diaPASEF: parallel accumulation-serial fragmentation combined with data-independent acquisition.* Nat Methods, 2020. **17**(12): p. 1229-1236.
17. Demichev, V., L. Szyrwiel, F. Yu, G.C. Teo, G. Rosenberger, A. Niewienda, et al., *dia-PASEF data analysis using FragPipe and DIA-NN for deep proteomics of low sample amounts.* Nat Commun, 2022. **13**(1): p. 3944.
18. Tyanova, S., T. Temu, P. Sinitcyn, A. Carlson, M.Y. Hein, T. Geiger, et al., *The Perseus computational platform for comprehensive analysis of (prote)omics data.* Nature Methods, 2016. **13**(9): p. 731-740.
19. Love, M.I., W. Huber, and S. Anders, *Moderated estimation of fold change and dispersion for RNA-seq data with DESeq2.* Genome Biology, 2014. **15**(12): p. 550.
20. Robinson, M.D., D.J. McCarthy, and G.K. Smyth, *edgeR: a Bioconductor package for differential expression analysis of digital gene expression data.* Bioinformatics, 2010. **26**(1): p. 139-140.
21. Ritchie, M.E., B. Phipson, D. Wu, Y. Hu, C.W. Law, W. Shi, et al., *limma powers differential expression analyses for RNA-sequencing and microarray studies.* Nucleic Acids Research, 2015. **43**(7): p. e47-e47.
22. Langfelder, P. and S. Horvath, *WGCNA: an R package for weighted correlation network analysis.* BMC Bioinformatics, 2008. **9**(1): p. 559.
23. Kim, S., *ppcor: An R Package for a Fast Calculation to Semi-partial Correlation Coefficients.* Communications for Statistical Applications and Methods, 2015. **22**(6): p. 665-674.
24. Sud, M., E. Fahy, D. Cotter, K. Azam, I. Vadivelu, C. Burant, et al., *Metabolomics Workbench: An international repository for metabolomics data and metadata, metabolite standards, protocols, tutorials and training, and analysis tools.* Nucleic Acids Res, 2016. **44**(D1): p. D463-70.
25. Perez-Riverol, Y., J. Bai, C. Bandla, D. García-Seisdedos, S. Hewapathirana, S. Kamatchinathan, et al., *The PRIDE database resources in 2022: a hub for mass spectrometry-based proteomics evidences.* Nucleic Acids Res, 2022. **50**(D1): p. D543-d552.

Supp Figure 1

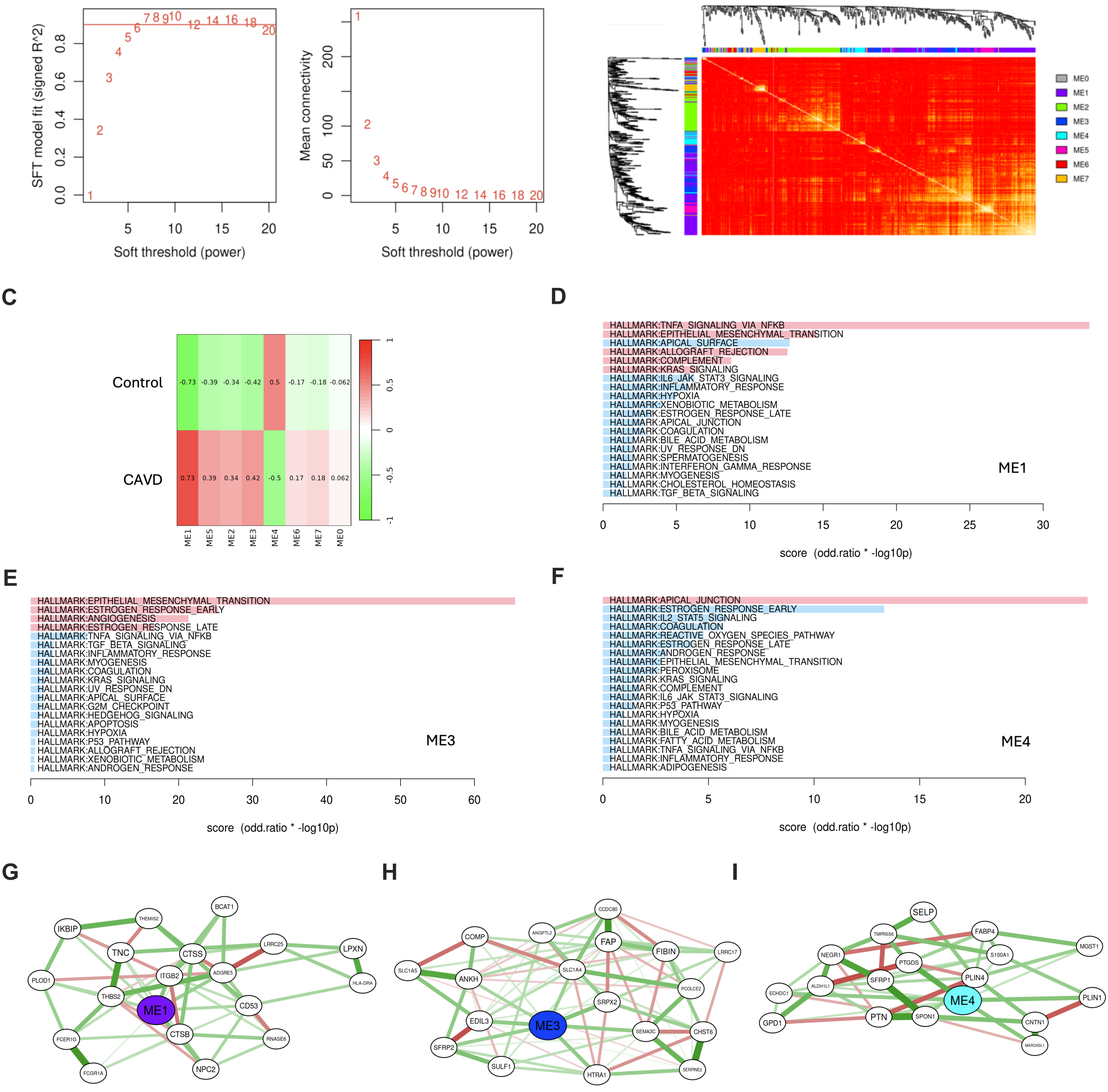

### Supp Figure 2

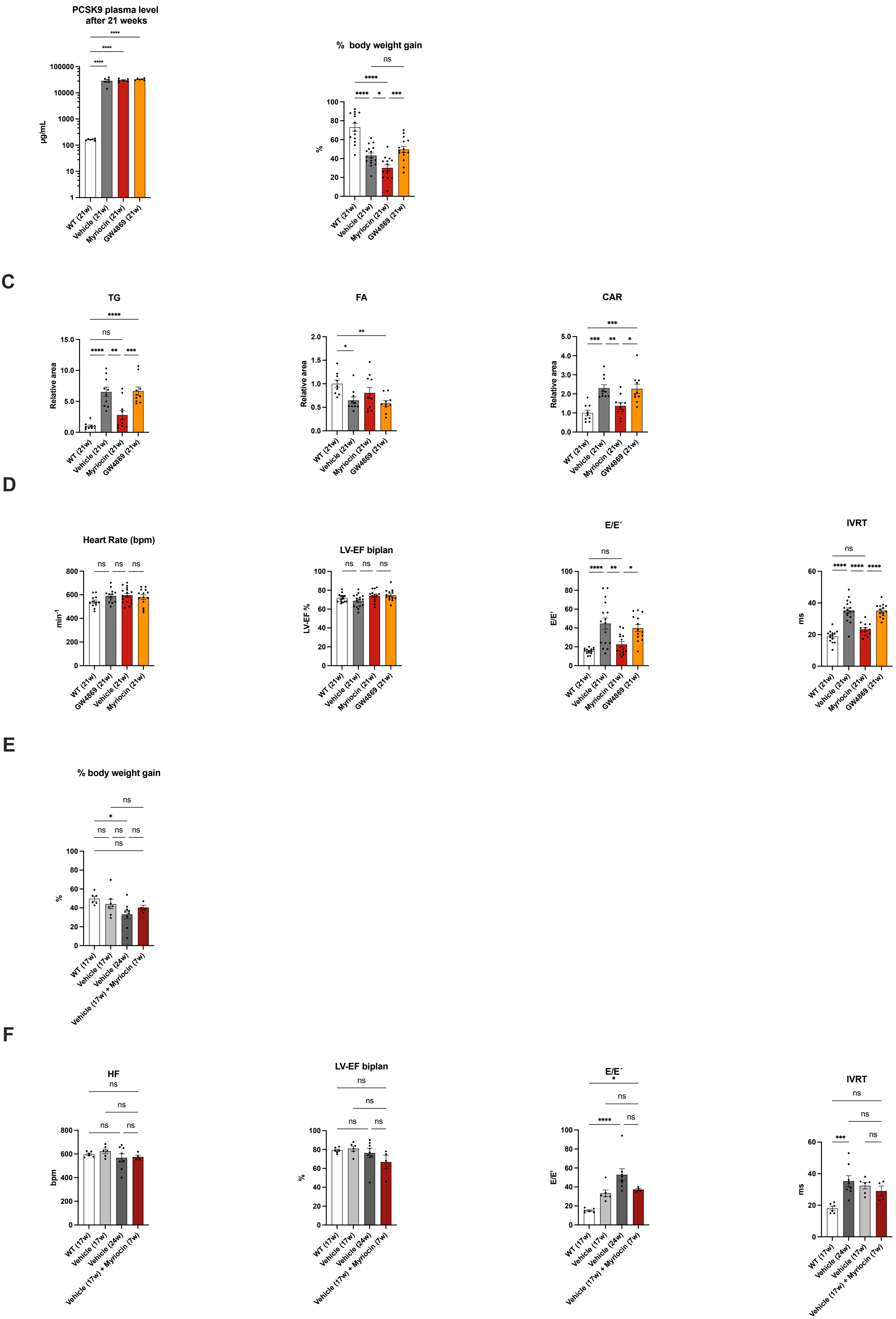

**A** Supp Figure 3

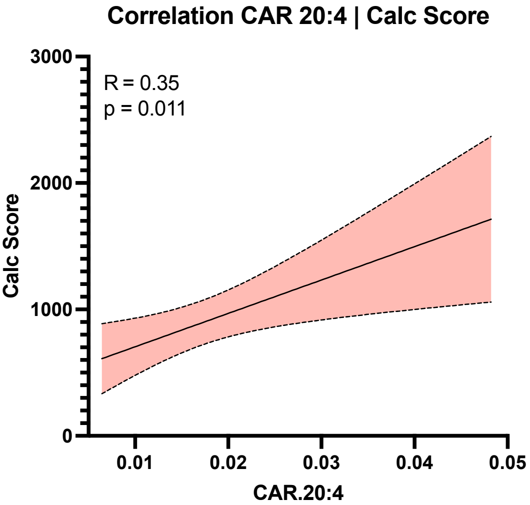

Supp Figure 4

A

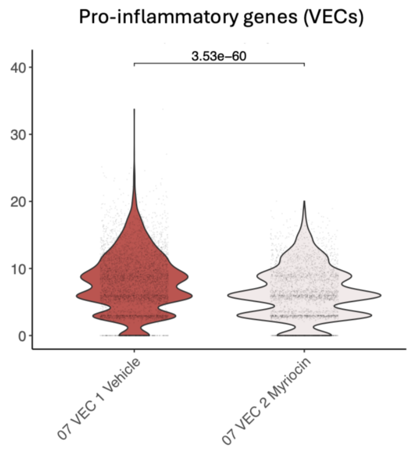


B

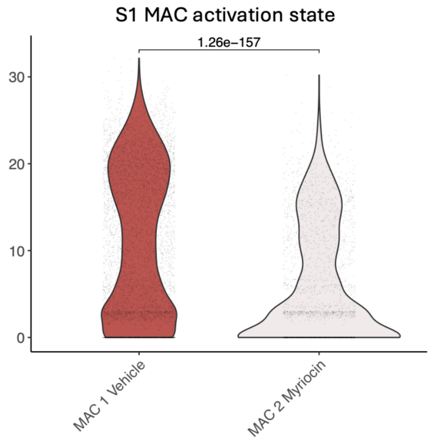


C

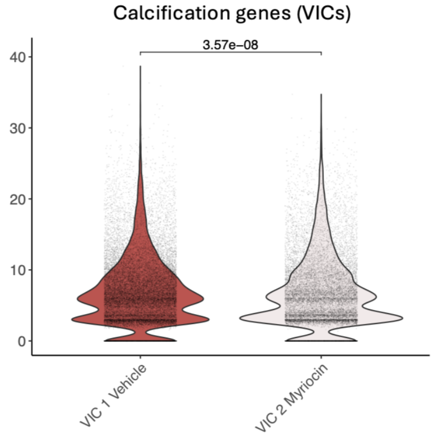


D

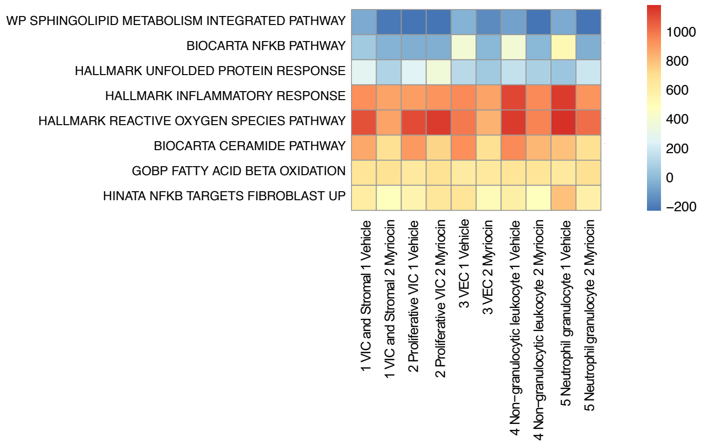

A

Supp Figure 5

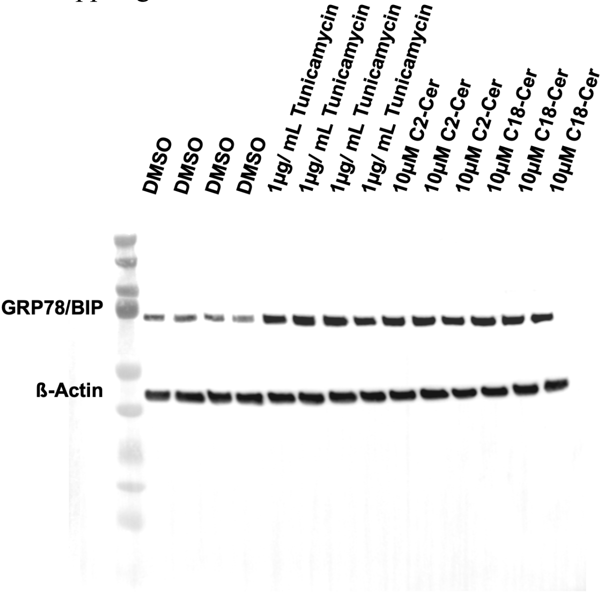

Supp Figure 6

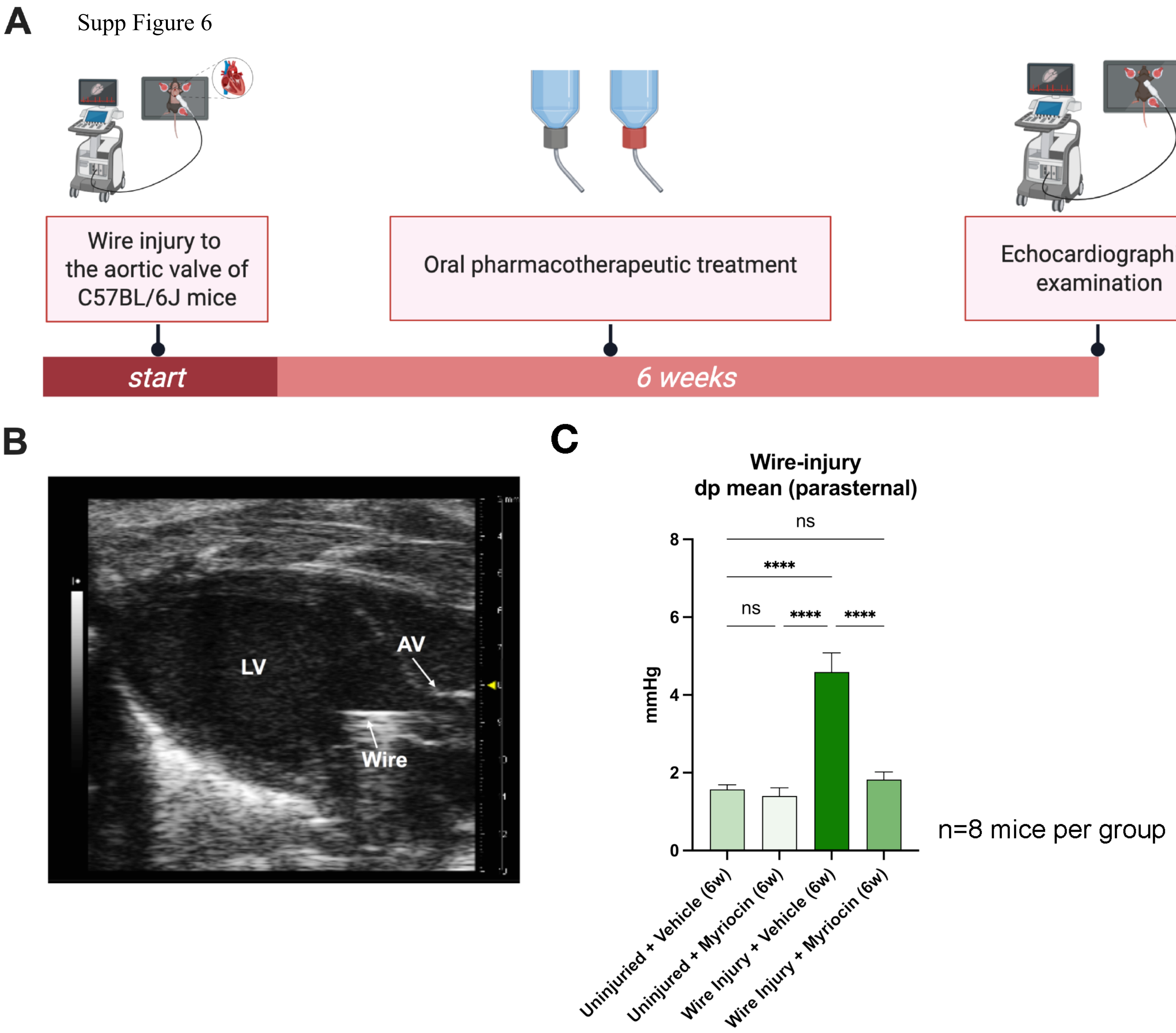
